# Screening cell-cell communication in spatial transcriptomics via collective optimal transport

**DOI:** 10.1101/2022.08.24.505185

**Authors:** Zixuan Cang, Yanxiang Zhao, Axel A. Almet, Adam Stabell, Raul Ramos, Maksim Plikus, Scott X. Atwood, Qing Nie

**Affiliations:** Department of Mathematics and Center for Research in Scientific Computation, North Carolina State University, Raleigh, NC 27695, USA; Department of Mathematics, The George Washington University, Washington, DC 20052, USA; Department of Mathematics, University of California, Irvine, CA 92697, USA; Department of Developmental and Cell Biology, University of California, Irvine, CA 92697, USA; The NSF-Simons Center for Multiscale Cell Fate Research, University of California, Irvine, CA 92697, USA

## Abstract

Spatial transcriptomic technologies and spatially annotated single cell RNA-sequencing (scRNA-seq) datasets provide unprecedented opportunities to dissect cell-cell communication (CCC). How to incorporate the spatial information and complex biochemical processes in reconstructing CCC remains a major challenge. Here we present COMMOT to infer CCC in spatial transcriptomics, which accounts for the competition among different ligand and receptor species as well as spatial distances between cells. A novel collective optimal transport method is developed to handle complex molecular interactions and spatial constraints. We introduce downstream analysis tools on spatial directionality of signalings and genes regulated by such signalings using machine learning models. We apply COMMOT to simulation data and eight spatial datasets acquired with five different technologies, showing its effectiveness and robustness in identifying spatial CCC in data with varying spatial resolutions and gene coverages. Finally, COMMOT reveals new CCCs during skin morphogenesis in a case study of human epidermal development. Both the method and the computational package have broad applications in inferring cell-cell interactions within spatial genomics datasets.

## Introduction

The complex structures and functions of multicellularity are achieved through the coordinated activities of various cells. Cells make decisions and accomplish their goals by interacting with an environment consisting of external stimuli and other cells. A major form of cell-cell interaction is cell-cell communication (CCC), which is mainly mediated by biochemical signaling through ligand-receptor binding that further induces downstream responses in shaping development, structure, and function.

Traditionally, CCC studies were often restricted to a few cell types and a small number of selected genes at the resolution of cell groups. Recently, the emergence of single-cell transcriptomics (scRNA-seq) has enabled us to examine tissues at single-cell resolution and with unprecedented genomic coverage^1^. Computational tools have been developed to estimate CCC activities from scRNA-seq data^2,3^ often using knowledge databases of signaling^4–6^. Most of these methods rely on the expression levels of ligand and receptor pairs and explicitly defined functions. In particular, the products of ligand and receptor gene expression^5,7^ or dedicated nonlinear models like Hill function-based formulas^6^ are used. In addition, these methods emphasize different aspects of CCC. To name a few, CellPhoneDB^5^, ICELLNET^7^, and CellChat^6^ account for the multi-subunit composition of protein complexes; SoptSC^8^, NicheNet^9^, and CytoTalk^10^ utilize downstream intracellular gene-gene interactions; and scTensor^11^ examines higher-order CCC, which is represented as hypergraphs. These CCC inference methods designed for scRNA-seq data have shed light on various biological systems based on non-spatial transcriptomic datasets^2,12,13^. However, the CCC activities inferred from scRNA-seq data often contain significant false positives since CCC only takes place within a certain range of spatial distance that is not measured in scRNA-seq datasets. Improvement can be made by filtering inferred CCC using spatial annotations^14^.

Spatial transcriptomics technologies^15–20^ provide distance information among cells or spots containing multiple or fractions of cells. With various cellular resolutions, these technologies measure spatial expression of hundreds to tens of thousands of genes in a 2-dimensional or a 3-dimensional tissue sample^21^. Methods and software packages^22–24^ have been developed for analysis of spatial data similar to scRNA-seq data. Computationally, only a small number of methods have been developed to analyze CCC specifically for spatial data. Giotto builds a spatial proximity graph to identify the interactions through membrane-bound ligand-receptor pairs^23^; CellPhoneDB v3 restricts interactions to cell clusters within the same microenvironment defined based on spatial information^25^; stLearn relates the co-expression of ligand and receptor genes to the spatial diversity of cell types^24^; SVCA^26^ and MISTy^27^ use probabilistic and machine learning models, respectively, to identify the spatially constrained intercellular gene-gene interactions; and NCEM fits a function to relate cell type and spatial context to gene expression^28^. However, current methods examine CCC pairwisely and locally, focusing on information between cells or in the neighborhoods of individual cells, as a result, the collective or global information in CCC, such as the competition between cells, are neglected.

Optimal transport, which globally connects two distributions by minimizing a total coupling cost function, has been recently used for transcriptomic data analysis, including batch effect correction^29^, developmental trajectory reconstruction^30^, and spatial annotation of scRNA-seq data^31,32^. Naturally, one can view ligand and receptor expression as two distributions to be coupled with a cost based on spatial distance to form an Optimal Transport (OT) problem^31,33,34^. However, using classical OT, when different molecule species with significantly different expression levels are normalized to ensure the same total mass, it makes the units of distributions incomparable. In addition, multiple ligand species can often bind to multiple receptor species, resulting in competition. Among the 1735 (secreted) ligand-receptor pairs in the Fantom5 database^35^, 72% (372/516) of ligands and 60% (309/512) of receptors bind to multiple species. Such competition between multiple molecule species is ubiquitous, a critical physical process ignored in the existing CCC inference methods. While recent OT variants such as unbalanced OT and partial OT can deal with unnormalized distributions and avoid certain coupling due to signaling spatial range and simultaneous consideration of multiple species^33,36–38^, they introduce some other issues. Specifically, unbalanced OT^38^ in its common form uses KL divergence as a soft constraint on marginal distribution preservation. This approach may result in the total coupled signaling molecule species significantly exceeding the total amount of either ligand or receptor initially available. On the other hand, partial OT^36^, requires an additional parameter, the total coupled mass, which is usually difficult to estimate in the context of CCC inference.

To adapt OT theory for the application of CCC inference, we present a method called collective optimal transport, which is capable of 1) preserving the original units, 2) ensuring the total signal not exceed the amount of an individual species (ligand or receptor), 3) enforcing spatial range limits of signaling, and 4) handling multiple competing species. The collective optimal transport method achieves these properties by optimizing the total transported mass and the ligand-receptor coupling simultaneously, unlike the existing optimal transport methods. By introducing an entropy regularization to enforce the inequalities for marginal distributions, the collective optimal transport can be reformulated as a special case of the general unbalanced optimal transport framework^38^. An efficient algorithm is developed specifically for solving the collective optimal transport problem.

Based on collective optimal transport, we develop COMMOT (COMMunication analysis by Optimal Transport), a package which 1) infers CCC by simultaneously considering numerous ligand-receptor pairs for both spatial transcriptomics data and the spatially annotated scRNA-seq data with estimated spatial distance between cells based on paired spatial imaging data; 2) summarizes and compares the spatial signaling directions; 3) identifies CCC downstream effects on gene expressions using ensemble of trees models; and 4) provides visualization utility for the various analyses.

We show that COMMOT accurately reconstructs CCC on simulated data generated by partial differential equation models and outperforms three related optimal transport methods. We then apply COMMOT to analyze scRNA-seq data spatially annotated using paired spatial datasets and five types of spatial transcriptomics data which have different properties in terms of spatial resolution and gene coverage. Finally, we examine a specific system of human epidermal development and reveal connections between CCC and skin development.

## Results

### Overview of COMMOT

In ligand-receptor interactions, multiple ligand species can bind to the same receptor species, or a receptor species can accommodate multiple ligand species in space (Fig. 1a). To incorporate this into an inference model, we develop a method named collective optimal transport (Fig. 1b). This method has three important properties: 1) taking unnormalized distributions and controlling the marginals of the transport plan by the original distributions to maintain the species units for more accurate estimate of the ligand-receptor coupling, 2) allowing infinity entries in the cost matrix to enforce spatial distance limits of CCC to avoid connecting cells that too far apart spatially, and 3) transporting multi-species distributions (ligands) to multi-species distributions (receptors) to account for the multi-species interactions among ligands and receptors (Fig. 1c).

**Fig. 1.**
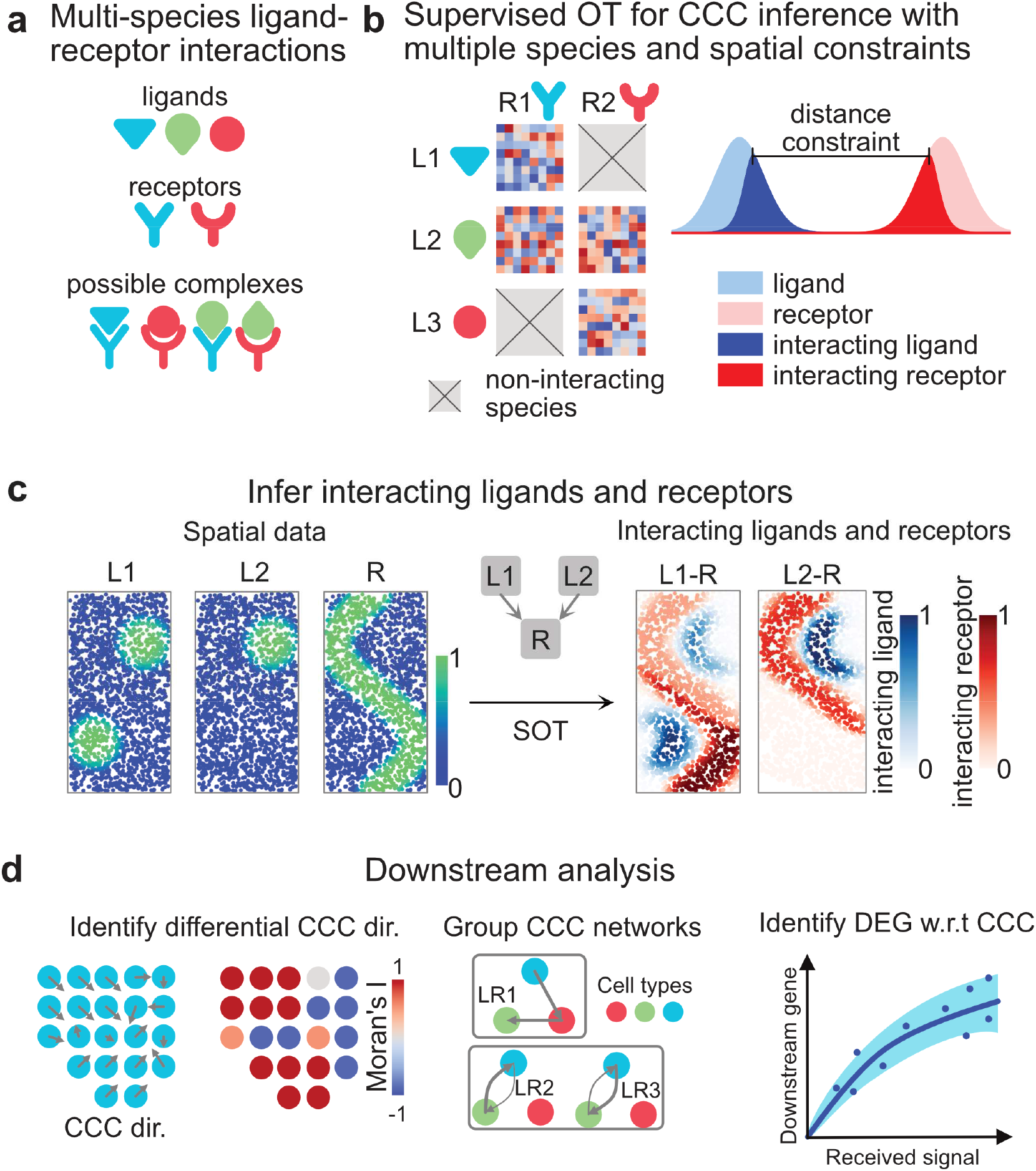
Overview of COMMOT. **a** COMMOT infers cell-cell communication (CCC) in space considering the competition between different ligand and receptor species. **b** Collective optimal transport infers CCC in space by introducing multi-species distributions and enforcing limited spatial ranges. **c** An example of inferring CCC for spatial distributions of ligand-receptor complexes from spatial distributions of the ligands and receptor where two ligand species compete for one receptor species. **d** Based on the inferred CCC network among cells or spots, we further 1) construct the spatial signaling direction trends, 2) summarize and group into spatial clusters, and 3) identify differentially expressed genes due to CCC.

Specifically, given a spatial transcriptomics (ST) dataset of *n*_*s*_ cells or spots of cells and *n*_*l*_ ligand species and *n*_*r*_ receptor species, the collective optimal transport determines an optimal multi-species coupling 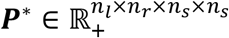 where 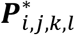 scores the signaling strength from sender cell *k* to receiver cell *l* through ligand *i* and receptor *j*. This is achieved by solving a minimization problem,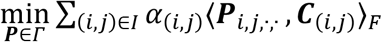 where 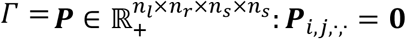 for 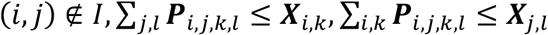, *I* is the index set for ligand and receptor species that can bind, and ***X***_*i,k*_ is the expression level of gene *i* on spot *k*. The species-specific cost matrix 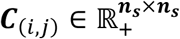 is the modified distance matrix among the spots where a pair of spots whose distance exceeds the spatial range of ligand *i* will result in an infinity entry. This construction differs from traditional optimal transport methods in that the exact marginal distribution matching restriction is relaxed to inequalities and infinity entries are allows in the cost matrix. The competitions between molecule species and cells are addressed in the optimization problem that a given receptor species or a cell have limited capacity for interaction, so higher inferred interaction with one particular ligand species or given cell will reduce the interaction score with other ligand species or cells. See Methods – COMMOT model and Methods – collective optimal transport algorithm, and Supplementary Information for details of formulations and derivation of algorithms.

For each ligand-receptor pair and each pair of cells or spots, the inferred CCC quantifies the ligand contributed by one spot to the ligand-receptor complex in another spot. Based on the inferred CCC, we further perform several downstream analyses: 1) interpolating the spatial signaling direction and identifying regions of difference in CCC, 2) summarizing and grouping CCC at the level of spatial clusters, and 3) identifying downstream genes whose expression is impacted by CCC (Fig. 1d). The spatial signaling direction is obtained by interpolating the cell-by-cell CCC matrix to a vector field in the intact tissue space to describe from or to what direction is the signal received or sent. For downstream genes, we first identify genes that are differentially expressed with the received signal. We then quantify the impact of CCC on these genes, while also considering the impact from other genes by incorporating a machine learning model that predicts the expression of a target gene based on both the received signal and other correlated genes. See Methods – Spatial signaling direction and cluster level CCC and Methods – Downstream gene analysis for the exact algorithms that perform the downstream tasks.

In summary, the COMMOT package performs ligand/receptor amount- and spatial-constrained inference of CCC considering the competition among different ligand and receptor species. Several useful downstream analyses include 1) summarizing the inferred CCC in terms of spatial signaling directions and generating cluster-level CCC, 2) comparing and grouping signaling pathways using quantitative metrics, and 3) screening and quantitative exploration of potential downstream genes.

### Accurate reconstruction of simulated spatial CCC

Ideally, CCC inference methods for spatial data should be validated by examining the spatial co-localization of ligand and receptor proteins. However, such data is often lacking. We therefore built partial differential equation (PDE) models to simulate CCC in space. The PDE model describes the essential physical processes in ligand-receptor interaction including the diffusion of ligands, binding and dissociation of ligand-receptor complexes, and degradation of molecules^39^ in 1-dimensional (1D) or 2-dimensional (2D) spaces (Fig. 2a).

**Fig. 2.**
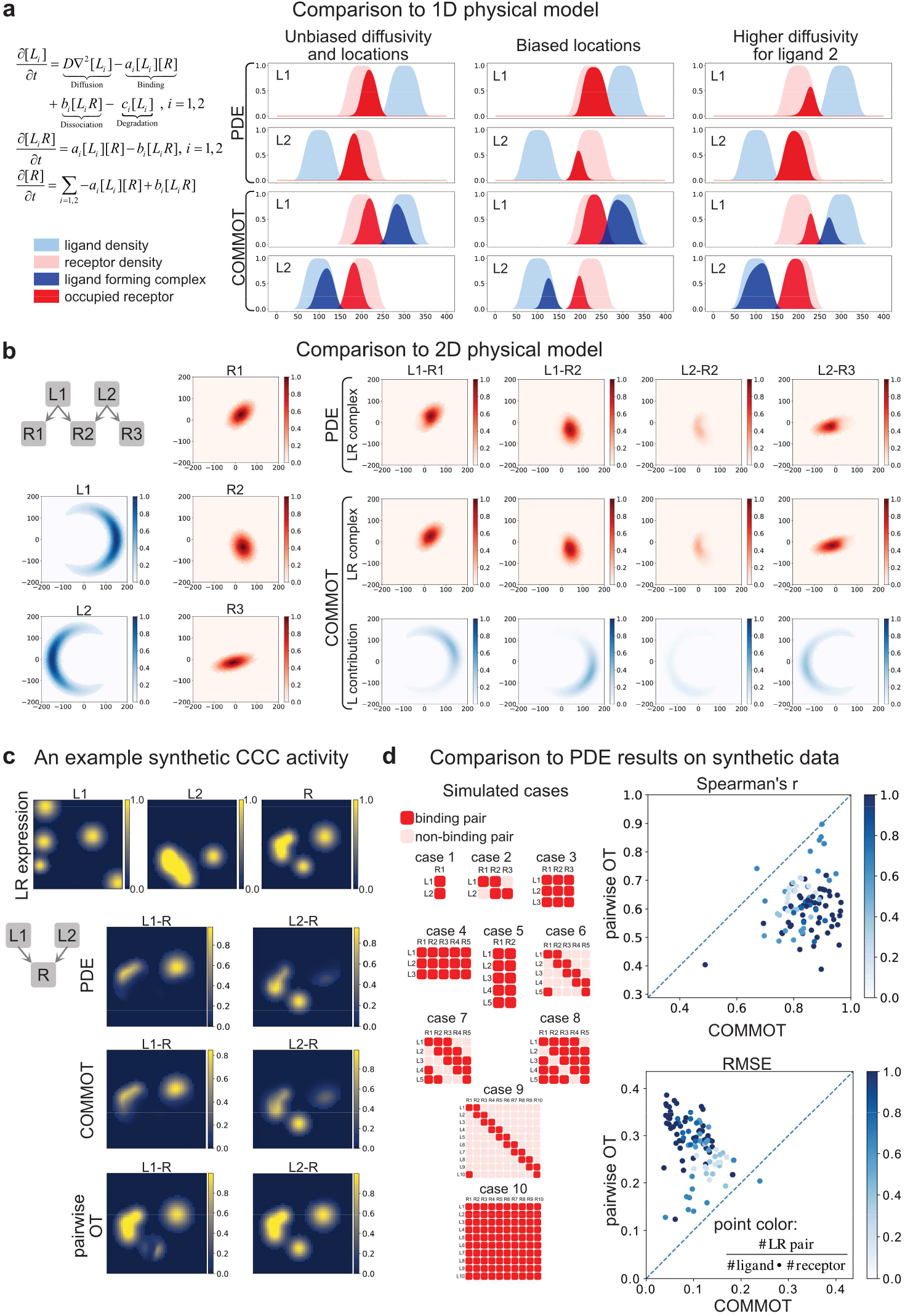
Validation using simulated data by partial differential equations (PDE) model. **a** The example PDE model where two ligand species can bind to the same receptor. The inference by COMMOT is compared to the simulation results in several 1-dimensional cases. **b** Comparison to simulated results in a 2-dimensional case with three ligand species and two receptor species. **c** An example of randomly generated 2-dimensional benchmark with two ligand species that binds to the same receptor. The simulated result, inference by COMMOT, and inference by pairwise method are shown. **d** Ten different cases of ligand-receptor binding and the performance of COMMOT and pairwise OT (with the same spatial limit as COMMOT but each LR pair examined separately) obtained by comparing to simulated results.

As an example, in the case of two ligand species and one receptor species (Fig. 2a, b), the two ligands could be Wnt2 and Wnt5a and the receptor could be Fzd5^35^. Then [*L*_1_] and [*L*_2_] describe the spatial distribution of free Wnt2 and Wnt5a whose production rates are observed from spatial data. Similarly, [*R*] describes the spatial distribution of unoccupied Fzd5 whose initial distribution is taken from spatial data. Then [*L*_1_*R*] and [*L*_2_*R*] describe the spatial distribution of Wnt2-Fzd5 and Wnt5a-Fzd5 complexes, respectively, which reconstruct the CCC activity in space. The quality of the inferred CCC will be assessed based on how accurate it reproduces the simulated spatial distributions of the ligand-receptor complexes.

We first illustrate COMMOT in two simulated examples in 1D and 2D. Specifically, the inferred spatial distribution of bound ligand-receptor complexes is compared with the simulated one. In the 1D physical model, the two ligand species are located on the two sides of the location of receptor, respectively and will compete to bind to the receptor with a limit spatial range of interactions. The effects of symmetric ligand locations, biased ligand locations, and different ligand diffusivities are correctly captured by collective optimal transport (Fig. 2a). In the 2D physical model, five interacting species are modeled, and the interaction patterns are also recapitulated (Fig. 2b). In addition to revealing ligand-receptor complexes, collective optimal transport also annotates the source of ligands that contributed to the ligand-receptor complexes (Fig. 2a,b).

Further quantitative comparison is carried out using simulations with randomly generated spatial expression of ligands and receptors in various cases of ligand-receptor binding (Fig. 2c). To account to different ligand-receptor binding situations, ten different cases are considered with different numbers of ligand and receptor species and different levels of competition (the ratio of interacting pairs over total number of species) among the species (Fig. 2d). For each case, ten random initializations are generated. The levels of bound ligand-receptor complex inferred by collective optimal transport are compared to the PDE simulation results using the root mean square error and Spearman’s correlation coefficient. Our results show that collective optimal transport consistently derives accurate predictions across the different cases and outperforms the results by evaluating each ligand-receptor pair independently using traditional optimal transport. The improvement over the results from the pairwise approach is especially significant when competition among species is high (Fig. 2d, Supplementary Figs. 1-4), for example, cases 4, 5 and 10. A comparison with simulated data demonstrates that collective optimal transport used by COMMOT accurately reconstructs CCC in space. The ability to handle non-probability distribution is especially useful when the total levels of ligand and receptor expressions are significantly different (Supplementary Fig. 5). Also, incorporating spatial information helps prioritizing interactions among cells that are spatially close (Supplementary Fig. 6). Moreover, our method outperforms two other variants of optimal transport that also do not require the input data to be probability distributions: unbalanced optimal transport and partial optimal transport (Supplementary Figs. 7-8). In unbalanced optimal transport, the commonly used KL divergence is adapted in this comparison while other divergence forms may be incorporated as well. The differences among these optimal transport variants are further illustrated by both a one-dimensional example (Supplementary Fig. 9) and with real ST data (Supplementary Fig. 10-11), showing again the advantages in using COMMOT.

**Fig. 3.**
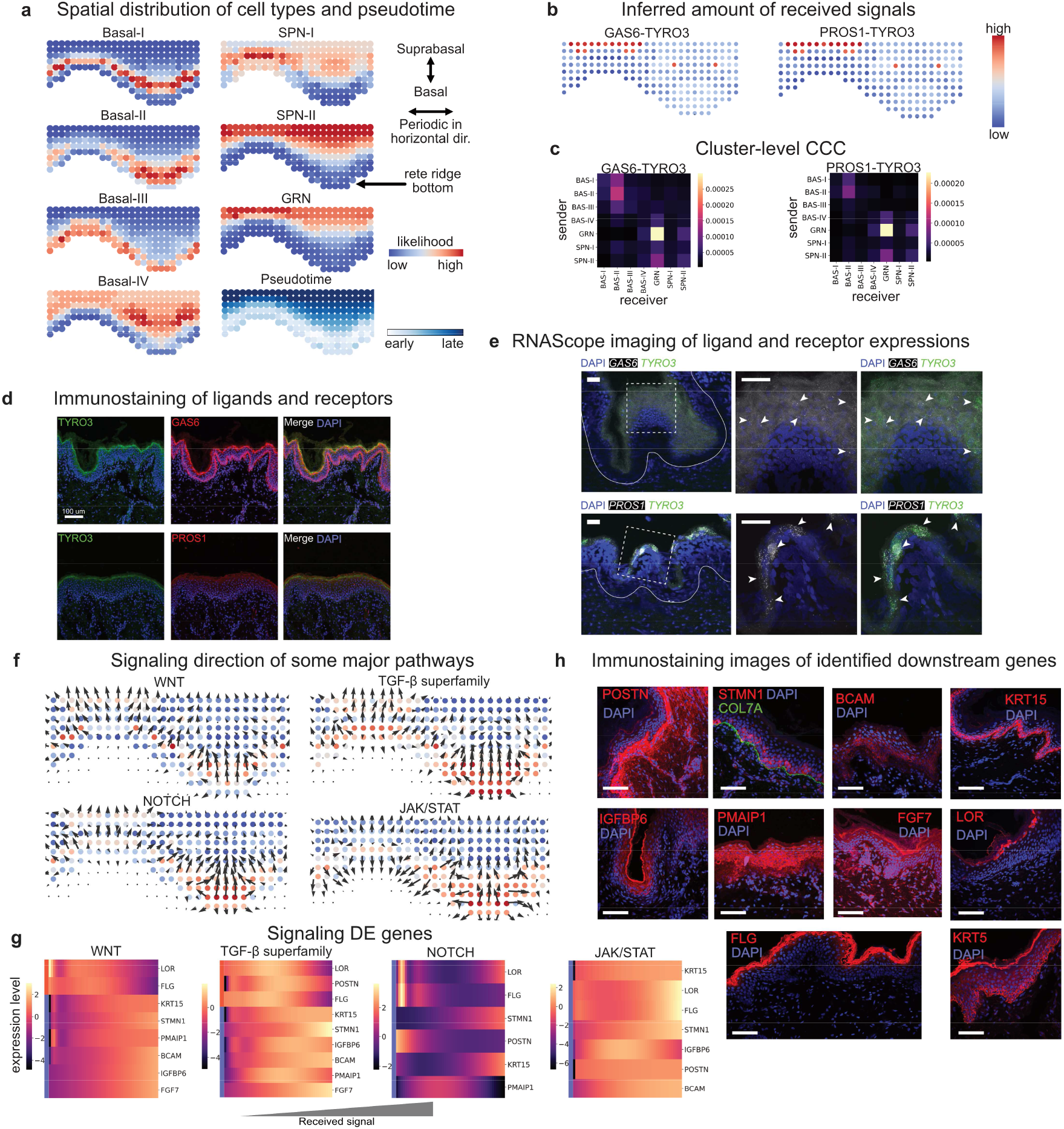
Role of CCC in human skin development. **a** Predicted spatial origin of the skin subtypes of cells in intact tissue and the pseudotime projected to space. **b-c** The inferred amount of received signals of two example ligand-receptor pairs, GAS6-TYRO3 and PROS1-TYRO3 at cell level and cluster level. **d** Immunostaining of proteins for GAS6, TYRO3, and PROS1. **e** Fluorescent in situ hybridization against RNA molecules for predicted L-R interactions in human epidermis (solid white outline; regions of interest are marked by a white dashed square). Top row shows expression patterns of Gas6 (white) and Tyro3 (green); bottom row shows expression patterns for Pros1 (white) and Tyro3 (green). In both cases, L-R signals that localize to stratum granulosum and are in close proximity to each other are indicated with white arrowheads. **f** The signaling directions of four major signaling pathways. **g** Heatmaps of selected signaling DE genes of the four signaling pathways respectively. The color represents gene expression level, and the horizontal axis depicts the amount of received signal. **h** Immunofluorescence staining images of the identified signaling DE genes supporting the identified correlation between WNT signaling and expression of these genes. Scale bars: d:100μm; e:100μm; h:100μm

**Fig. 4.**
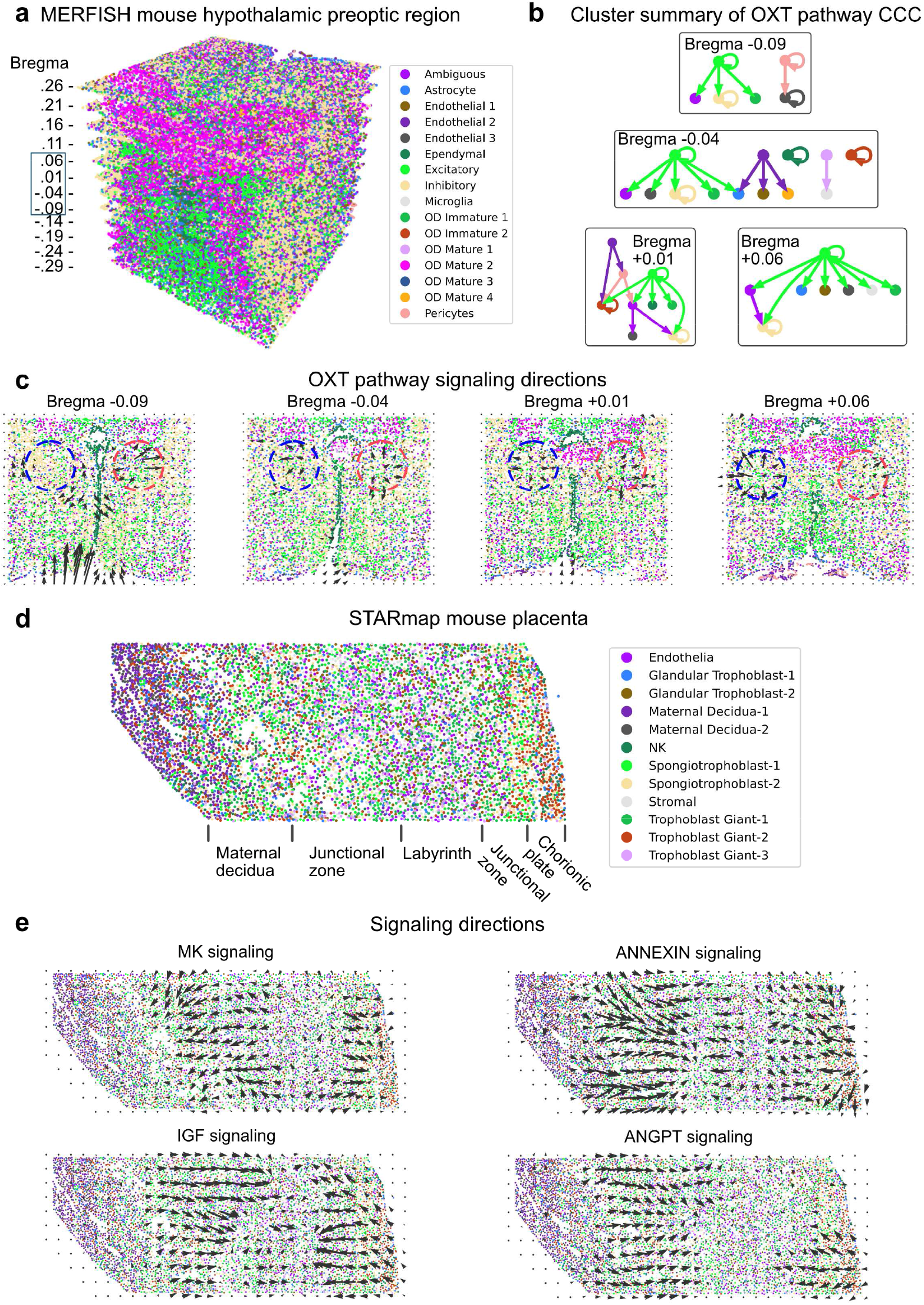
Inference of signaling direction in single-cell resolution spatial transcriptomics data. **a** MERFISH data of mouse hypothalamic preoptic region with multiple slices across the anterior-posterior axis^44^. **b** Cluster-level summary of CCC through oxytocin (OXT) signaling pathway. **c** Signaling directions of OXT pathway. **d** STARmap data of mouse placenta^46^. **e** Signaling directions of Midkine (MK), insulin-like growth factor (IGF), Annexin, and angiopoietin (ANGPT) pathways.

**Fig. 5.**
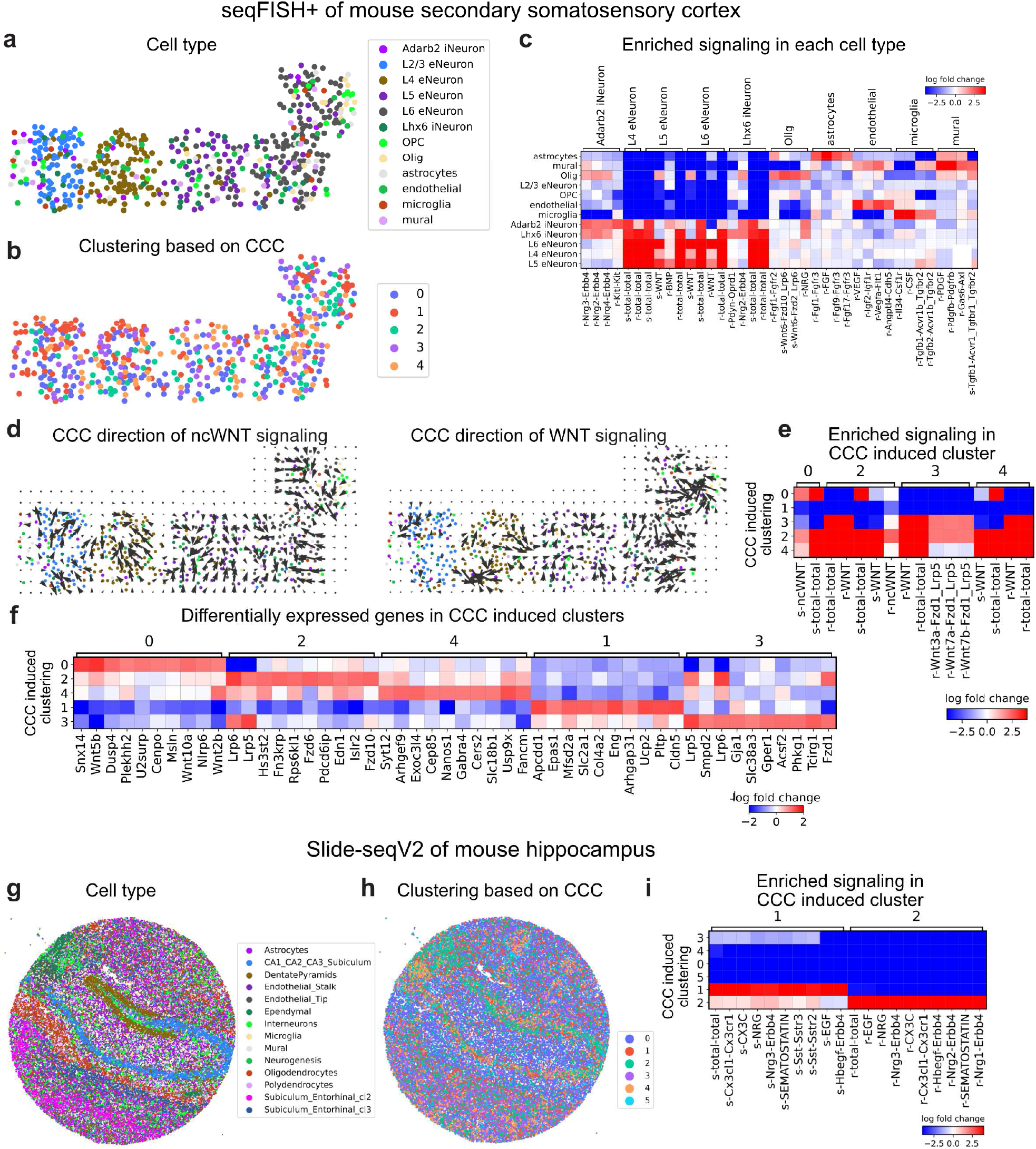
Downstream analysis of inferred CCC in single-cell resolution spatial transcriptomics data. **a-f** CCC analysis of a seqFISH+ data mouse secondary somatosensory cortex. **a** Clustering of cell type based on gene expression. **b** Clustering based on inferred CCC. **c** Enriched signaling in each cell type. **d** Spatial signaling direction of ncWnt and Wnt signaling pathways. **e** Enriched signaling in CCC induced clusters. **f** Differentially expressed genes in the CCC induced clusters. **g-i** CCC analysis of a Slide-seq (v2) data of mouse hippocampus. **g** Clustering of cell type based on gene expression. **h** Clustering based on inferred CCC. **i** Enriched signaling in CCC induced clusters.

**Fig. 6.**
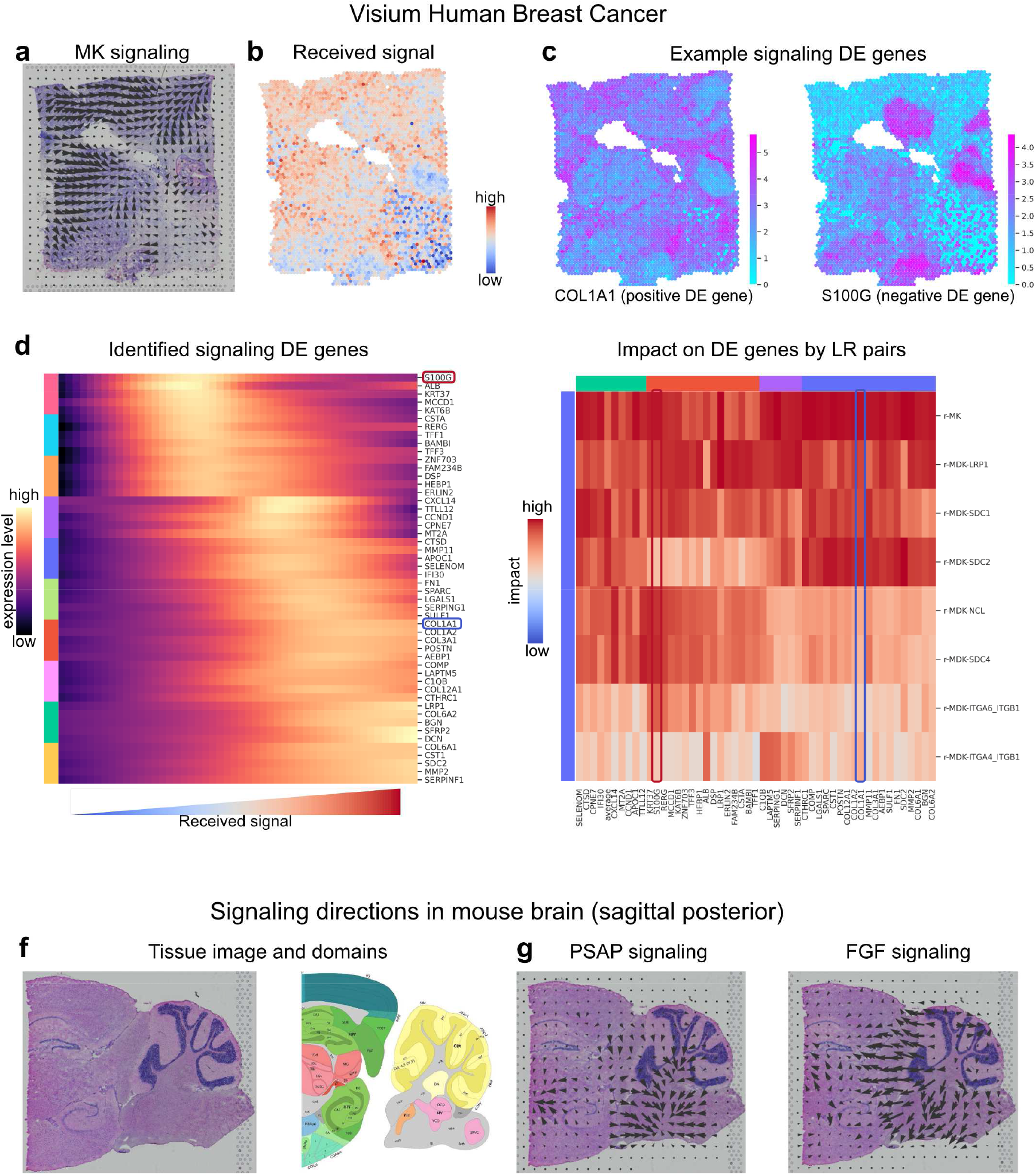
CCC inference on Visium spatial transcriptomics data. **a**-**e** Midkine (MK) signaling in breast cancer tissue. **a** Spatial signaling direction. **b** Amount of received signal by each spot. **c** Two example differentially expressed genes due to signaling. **d** The differentially expressed genes due to the total amount of received signal of MK signaling pathway. **e** Unique impact on the identified differentially expressed genes by the individual ligand-receptor pairs. **f**-**g** Signaling in mouse brain tissue. **f** The H&E image of the Visium data of the mouse brain tissue and the reference tissue domain annotation by Allen Brain Atlas. **g** Spatial signaling direction for prosaposin (PSAP) and fibroblast growth factor (FGF) signaling pathways, respectively.

### The roles of CCC in human epidermal development

Next, we applied COMMOT to examine development of epidermis in human skin. Our recent work profiled neonatal human epidermis on scRNA-seq and found four stem cell clusters that reside in the innermost basal layer of the epidermis, a differentiating spinous (SPN) cell cluster in the intermediate layer, and a granular (GRN) cell cluster in the outermost living layers^40^. The four basal cell clusters were spatially resolved by immunofluorescence staining and shown to reside either at the tips of the so-called epidermal ridges (Basal-III), ridge valleys (Basal-IV), or heterogeneously, dispersed throughout the ridge space (Basal-I and -II). To validate and further refine spatial relationships between diverse epidermal cells, we constructed an *in situ* spatial transcriptomic map using the tool called SpaOTsc^31^ by integrating spatial data digitalized from immunofluorescence staining images of 208 cells and 12 genes and scRNA-seq data containing 4264 cells and 33694 genes^40^. A leave-one-out prediction of the spatially measured genes was performed to confirm the quality of the integrated data (Supplementary Fig. 12). The integrated dataset also correctly identifies *a priori* known locations of the epidermal cell types and agree with known developmental path by epidermal cells from basal to suprabasal layers (Fig. 3a).

Next, we used COMMOT to examine spatial signaling activities among epidermal cells by considering ligand-receptor pairs annotated in the CellChatDB database. Our computational analysis predicts that molecular interactions between ligands GAS6 and PROS1 with their receptor TYRO3 (TYRO3-GAS6 and TYRO3-PROS1) are significant in granular cells and moderately present in basal cells (Fig. 3b). This prediction was confirmed by both immunostaining for proteins (Fig. 3d) and using RNAscope to stain for RNA (Fig. 3e).

At the signaling pathway level, we then examined four specific pathways known to have important roles in epidermal homeostasis, namely the WNT, TGF-β, NOTCH, and JAK/STAT pathways^40^ (Fig. 3f, Supplementary Figs. 13-16). For all four pathways, we observe mainly vertical upward with some downward signaling directions in the basal layers at the bottom of the ridges (Fig. 3f). WNT signaling is known to promote basal stem cell proliferation^41^, whereas TGF-β suppresses it^42,43^. Thus directional signaling generally originating from the suprabasal layer may regulate communication to basal cells on how much or how little to proliferate. We also see strong downward signaling direction at the top of the ridges for JAK/STAT signaling suggesting that there may be strong communication between the suprabasal and basal compartments. JAK/STAT signaling is implicated in several hyperproliferative skin diseases^44^, suggesting that the Basal-III and Basal-IV clusters at the top and bottom of the epidermal ridges may be distinctly implicated in these diseases.

Based on inferred signaling activities, we further identified some of the corresponding differentially expressed genes for each signaling pathway and modeled their expression level changes with increasing received signal without further considering spatial information (Fig. 3g). For the WNT pathway, increasing signal results in higher expression of known basal cell markers *KRT15* and *KRT5*, as well as lower expression of known terminally differentiated GRN markers *LOR* and *FLG*, reinforcing the WNT pathway’s known role in stem cell proliferation^41^. The analysis also predicted that higher WNT signaling would increase the expression of *BCAM, POSTN*, and *STMN1*, the expression localization of which we confirmed by immunostaining on human epidermis (Fig. 3h). Interestingly, computationally we also predicted *IGFBP6, PMAIP1* and *FGF7* to positively correlate with WNT signaling, but we observe their expression mainly in the SPN and GRN layers, possibly due to predicted WNT signaling in both directions in Basal-IV (Fig. 3h). TGF-β signaling showed a similar profile to the WNT pathway, with NOTCH and JAK/STAT signaling showing a more complex response (Fig. 3g). These results suggest how testable hypotheses can be derived from inferred signaling activities.

### Signaling analysis in high spatial resolution ST data

We first explore CCC in ST data with high spatial resolution using the CellChatDB database^6^. We analyzed MERFISH data of the mouse hypothalamic preoptic region with 161 genes and 73655 cells across 12 slices along the anterior-posterior axis^45^ (Fig. 4a-c). Among the signaling pathways available in the data, oxytocin (OXT) signaling, an important pathway modulating social behaviors, was found to be most active. Self-modulation of excitatory neurons and modulation of inhibitory neurons by excitatory neurons through OXT signaling were identified across all the slices (Fig. 4b, Supplementary Fig. 17-18) a result consistent with the known major functions of OXT signaling^46^. Further analysis identifies the local regions of high OXT signaling activity and the spatial direction of OXT signaling (Fig. 4c), which corresponds with protein staining of OXT and its receptor^47^. A gradual change of predicted signaling direction and high-activity regions is observed through the adjacent slices (Fig. 4c, Supplementary Fig. 19).

We then analyzed a STARmap data of mouse placenta with 903 genes and 7203 cells^48^ (Fig. 4d). The midkine (MK) and insulin-like growth factor (IGF) signalings were found to be active in the same regions (Fig. 4e). Interestingly, the inferred signaling directions of MK and IGF signalings are opposite, suggesting a potential feedback loop between the two signaling pathways^49^. In addition, it was found that signaling through the ligand-receptor pair Igf2-Igf2r is active in the labyrinth region and in endothelial cells, both of which are consistent with our predictions^50^. MK signaling was inferred to be active in trophoblast cells, consistent with the previous finding on the role of SDC1 and SDC4 in trophoblast cells^51,52^ (Supplementary Fig. 20). We also found similar active regions and directions for Annexin and Angiopoietin (ANGPT) signalings suggesting that they might function cooperatively (Fig. 4e).

To perform the downstream analysis of CCC, we first studied seqFISH+ data of mouse secondary somatosensory cortex with 10000 genes measured in 523 individual cells^18^ (Fig. 5a-f). The inferred CCC assigns each cell a CCC profile quantifying the amount of signal sent or received through each ligand-receptor pair assembling a (*n*_*s*_ × *nr*_*lr*_) CCC profile matrix where for the *n*_*s*_ cells and *n*_*r*_ ligand-receptor pairs. Using the CCC profiles to cluster the data, a CCC-induced clustering is obtained and cells within the same group are expected to have similar signaling activities (Fig. 5b). As opposed to the spatial separation of cell types, we observe the CCC-induced clusters to be mixed in space suggesting that different cell types might have similar signaling activities through multiple signaling pathways.

We next conducted enrichment analysis for each clustering-profile pair with the gene expression-induced clustering or CCC-induced clustering as the label assignment, and the gene expression or the CCC activity as the profile. For the cell type clustering-CCC activity profile pair, the neuron cells were found to be most active through various ligand-receptor pairs and differential signaling activities were identified for the relatively rare cell types (oligodendrocytes, astrocytes, endothelial, microglia, and mural) (Fig. 5c). Among the two most active signaling pathways, WNT signaling is colocalized in local regions, while noncanonical WNT (ncWNT) signaling is widely distributed across the sample (Fig. 5d). COMMOT predicted significant WNT signaling in neurons (Supplementary Fig. 21), correlating well with critical roles of WNT signaling in neuronal migration and activity in the somatosensory cortex^53^. In addition to cluster 2 and 4 in the CCC-induced clustering that shows hyperactive signaling, cluster 0 and 3 were found to be significant signal senders and receivers, respectively (Fig. 5e). We further identified differentially expressed genes that match the signaling patterns of each of the CCC-induced clusters (Fig. 5f). This analysis reveals known signaling components of the relevant pathways and may represent novel regulators of each pathway. For example, the positive differentially expressed genes associated with cluster 0 (WNT signal senders) include known WNT ligands *Wnt5b, Wnt10a*, and *Wnt2b*, and the differentially expressed genes in cluster 3 (WNT signal receivers) include known target genes of WNT signaling pathway such as *Gja1* and *Acsf2*, and the known corresponding intracellular signaling transductors *Lrp5* and *Lrp6* (Fig. 5f).

We further jointly analyzed mouse cortex datasets generated with three different technologies including Visium, seqFISH+, and STARmap. Across these three datasets, AGT signaling was identified in neurons. Spatially, neurons in the L2-3 region were identified as strong receivers of AGT ligands across the three datasets. Interestingly, a striped signaling pattern was observed, wherein strong signals within individual layers form stripes, while weak signals form inter-stripe regions. Strong AGT signaling activity among oligodendrocytes was also identified in both STARmap and seqFISH+ datasets. (Supplementary Fig. 22). In both Visium and seqFISH+ cortex datasets, we inferred WNT signaling to be active across different cortical layers. In both datasets, we identified WNT signaling to be relatively low in layer 5, compared to other layers. This result is in line with previous study showing that disruption to WNT signaling in *Lrp6* mutant mice leads to cortical hypoplasia and reduced proliferation in neurons in layers 2-4 and 6, with neurons in layer 5 remaining relatively unchanged^54^ (Supplementary Fig. 23). TAC (tachykinin neuropeptide family) signaling activity was consistently found in both Visium and STARmap cortex datasets to be active in non-neuronal cells and in inhibitory neurons, especially in somatostatin-expressing neurons (Sst), because of the high expression of tachykinin receptor-3 in Sst neurons^55^ (Supplementary Fig. 24).

We also applied COMMOT to a large-scale ST dataset, Slide-seqV2 data of mouse hippocampus measuring expression of 23264 genes in 53173 beads (spatial spots) whose size is comparable to individual cells^56^ (Fig. 5g-i). Clustering based on CCC profile separates the spots into six clusters where cluster 1 and 2 mostly consists of DentatePyramid, and the CA1_CA2_CA3_Subiculum, and interneuron cells are generally active in CCC (Fig. 5g-i).

### Signaling analysis in multi-cell resolution ST data

Finally, we applied COMMOT to signaling analysis with Visium^16^ spatial transcriptomics data, where each spatial spot consists of multiple cells. By analyzing the breast cancer data with 3798 spots and 36601 genes, we found clear spatial signaling directions of Midkine (MK) signaling which was identified to be the most active (Fig. 6a), and the regions receiving such signals (Fig. 6b). To identify the genes that may be regulated by or regulate CCC, we used the tradeSeq^57^ to perform a differential expression test for which the amount of MK reception was used as the cofactor, analogous to a temporal differential expression test where pseudotime is used as the cofactor (Fig. 6c,d). COL1A1 was identified as a significantly positive differentially expressed gene with a distinct spatial pattern, whereas S100G was a significantly negative differentially expressed gene with its own unique spatial pattern (Fig. 6c). Furthermore, the level of COL1A1 expression increases while the level of S100G expression decreases as MK signals increase (Fig. 6d). Similar to the temporal DE gene analysis in scRNA-seq data, the signaling DE gene analysis in ST data identifies relation between gene expression and signaling activity, for example, the connection between COL1A1 expression and MK signaling. Therefore, ST data with good coverage of genes and large number of cells or spots is preferred for such analysis.

The differential expression tests typically examine the pairwise correlation between a potential target gene and a cofactor. The higher order interactions among multiple factors (multiple potential upstream genes and the cofactor) are often neglected. To further prioritize the genes that are more likely regulated by CCC, we used a random forest model^58,59^ where the potential target gene is the output. The CCC cofactor and the top intracellular correlated genes are the input features. The feature importance of the cofactor in the trained model then serves as a reference of the unique information provided by the cofactor about the potential target gene, scoring the unique impact of individual ligand-receptor pairs on each of the identified signaling differentially expressed genes. Using this model, we observe that COL1A1 and S100G are distinctly impacted by various MK ligand-receptor pairs (Fig. 6e). Generally, this analysis can be carried out for any ligand-receptor pair that is expressed in the data, for example, the PD1 signaling pathway related to T cell functions (Supplementary Fig. 25).

We also analyzed a Visium^16^ dataset of mouse brain tissue with 3355 spots and 32285 genes (Fig. 6f). We found significant PSAP signaling activity widely across the tissue (Fig. 6g), where broad protective roles of PSAP in the nervous system have been discovered^60^. In addition, FGF signaling activity was identified on the border of cerebellar cortex (Fig. 6g), consistent with its known role in patterning of the cerebellum during development^61^.

### Robust identification of spatial signaling direction and downstream targets

To further evaluate COMMOT, we assess the method robustness, study the correlation between inferred CCC and the expression of known downstream genes, and compare with two existing methods, CellChat^6^ and Giotto^23^ that are designed for scRNA-seq and spatial transcriptomics data respectively.

For the robustness study, we utilized the stage 6 *Drosophila* embryo, an extensively studied system, which contains approximately 6000 cells that have been extensively studied with *in situ* databases available that present systematically annotated spatial gene expression^62,63^. The paired data consists of scRNA-seq of 1297 cells and 8925 genes along with the spatial single-cell resolution data of 3039 cells and 84 genes^64^, with both good depth and spatial organization. We first integrated them using SpaOTsc^31^ to generate two imputed datasets, the imputed scRNA-seq with predicted spatial coordinates of cells and the imputed spatial data with predicted expression of the 8925 genes from the scRNA-seq data. Using subsampling as a test for robustness, we randomly subsampled the data and compared them to the results from the full dataset.

When decreasing the subsample size, the cosine distance (for spatial signaling direction) and Jaccard distance (for cluster level signaling) were found to remain relatively small showing that both signaling directions and cluster level signaling are robustly captured in the subsampled data (Supplementary Fig. 26; See Methods – Evaluation metrics for detailed calculation of the cosine distance and Jaccard distance). The differentially expressed signaling genes were then identified by using the inferred amount of received signal as the co-factor, analogous to the approach for pseudotime^57^. The DE genes identified from subsampled datasets were found to be comparable to the ones identified from the full dataset (Supplementary Fig. 26).

Next, utilizing scSeqComm, a database of known target genes of ligand-receptor pairs^65^, we investigated the correlation between the inferred signaling activities and the expression of the corresponding target genes. scSeqComm is used because it combines multiple major resources including Reactome, TTRUST, and RegNetwork. We used three datasets analyzed in the previous sections with transcriptome or near transcriptome gene coverage: Visium human breast cancer data, Visium mouse brain data, and seqFISH+ mouse somatosensory cortex data. COMMOT was used to quantify all available ligand-receptor pairs in CellChatDB database which were then summarized such that each spot was assigned a score to quantify the level of received signal through each signaling pathway. At the individual-spot level, Spearman’s correlation coefficient was computed for each ligand-receptor pair between the received signal and the average expression of the known downstream genes. The median correlations on the three datasets are 0.237, 0.180, and 0.230, respectively (Supplementary Fig. 27). At cluster-level, we quantified the level of received signal using the averages among the spots of the clusters. We also compare with three widely-used methods that infer cluster-level cell-cell communication network, CellChat^6^, Giotto^23^, and CellPhoneDB v3^25^ where CellChat is designed for scRNA-seq data and Giotto and CellPhoneDB v3 are designed for spatial transcriptomics data. The activity of the downstream genes of a ligand-receptor pair is quantified as the percentage of significant positive DE genes of a cluster. The comparison shows a significant correlation between the inferred CCC and the activity of known downstream genes, and we found COMMOT achieves stronger correlation than the other three methods for most of the datasets and comparable to CellPhondDB v3 in some cases (Supplementary Fig. 28-29). Some examples of CCC inferred by COMMOT and the expression of downstream genes are shown in Supplementary Fig. 30. This evaluation can be further improved if more complete knowledge of gene regulations is available. With such list, one may also formulate the evaluation as a classification problem.

Finally, using Visium human breast cancer and mouse brain datasets, we studied the difference between COMMOT and the three other methods. When assessing the distributions of inter-cluster spatial distance of the identified significant CCCs, the significant CellChat interactions have a relative uniform distribution of the spatial distances. This is expected since CellChat was designed for scRNA-seq data and spatial organization is not considered in the method. Both COMMOT and Giotto prioritize CCC between spatially close clusters while COMMOT identifies more CCC links that are beyond the neighboring cells because Giotto uses cell-contact graph that might limit the range of inferred CCC. CellPhoneDB v3 identifies more CCC links than COMMOT potentially due to its consideration of inter-cluster distances rather than distance among individual cells (Supplementary Figs. 31-36). In the comparison to CellChat, COMMOT uniquely detects CCC that are not differentially expressed in clusters but spatially significant (Supplementary Figs. 31-32). Giotto characterizes the average expression of ligands and receptors of spatially neighboring spots without providing the role for each spot in CCC. COMMOT identifies each spot as a sender, receiver or both in CCC with interpretable results (Supplementary Figs. 33-34). COMMOT identifies CCC that is locally significant which CellPhoneDB v3 may neglect due to its consideration of expression level of entire clusters (Supplementary Figs. 35-36).

To study the efficiency of the CCC inference algorithm, we found COMMOT running time scales linearly to the number of nonzero elements in the CCC (Supplementary Fig. 37). The number of CCC scales linearly to the number of locations in ST data due to the spatial range constraint, and the memory usage also scales linearly to the number of locations since only the finite values of the cost matrix and the non-zero values of the CCC matrix need to be stored. Thus, COMMOT can effectively handle the existing ST datasets since the computing time and the memory usage both scale linearly to the number of spatial locations.

## Discussion

To dissect CCC from the emerging spatial transcriptomics data, we developed COMMOT, a spatial analysis tool, to (1) infer CCC for all ligand and receptor species, simultaneously, (2) visualize spatial CCC at various scales including a novel vector field visualization of spatial signaling directions, and (3) analyze their downstream effects. This tool is based on a novel computational method, collective optimal transport, which incorporates both competing marginal distributions and constrained transport plans, two important properties that cannot be dealt with using the current variants of optimal transport.

We have studied a wide range of data types of different spatial resolutions and gene coverage: *in silico* spatial transcriptomics data obtained by integrating scRNA-seq and spatial staining data, Visium, Slide-seq, and seqFISH+ spatial transcriptomics. COMMOT can consistently capture the CCC activities from the simulation data and the real spatial data. For instance, dorsal-to-ventral Dpp signaling and anterior-to-posterior Wg signaling were observed in the *Drosophila* embryo, consistent with previous *in vivo* data^66, 67^. Our simulations point to a significant TGF-β signaling activity in the hippocampal region and WNT signaling activity in the cerebellar cortex of the mouse brain, coinciding with the known signaling roles of the above pathways in those regions^68,69^. COMMOT is a useful predictive tool to study spatial positions of differentially expressed gene products. In human skin, COMMOT identified that higher WNT signaling increases the expression of *BCAM, POSTN*, and *STMN1*. As WNT signaling has known roles in stem cell proliferation^41^, our analysis suggested that the gene products of WNT signaling would likely be expressed in the basal stem cells, which we confirmed by immunofluorescence staining the three genes above of human epithelia.

Having said that, we acknowledge that false positives in our inferred CCC are inherently possible because spatial transcriptomics data does not directly represent protein abundancy and our method cannot capture protein-specific modifications, such as protein phosphorylation, glycosylation, proteolytic cleavage into fragments, dimerization, that certainly affect their signaling functions and, thus, CCC mechanisms, that COMMOT aims to infer. The reliability of CCC predictions is expected to significantly improve once the emerging spatial proteomics approaches will mature.

The spatial distance constraint in the model is used to capture the effect of ligand diffusivity, which is determined by several factors, including protein weight and tortuosity of extra-cellular space^70^. When screening all ligand-receptor pairs, it is difficult to estimate this parameter for each pair accurately. In our model, the local short-range interactions are emphasized even when the spatial distance range is increased (Supplementary Fig. 38). Thus, when screening many ligand-receptor pairs, a uniform and relatively large spatial distance limit may be used in the model to avoid missing important interactions. Once the important interactions are identified, using an accurate estimation of this parameter would further refine the prediction to remove some false positive CCC links.

Like other existing CCC inference methods using scRNA-seq data, COMMOT outputs static CCC activity networks among cells or clusters using spatial data. With a foreseeable deluge of spatial omics data, the temporal sequences of spatial transcriptomics data will soon further enable the revealing of CCC dynamics^71^. When paired with dynamic formulation of optimal transport, the collective optimal transport will be a powerful tool to incorporate multi-omics data for spatiotemporal dynamics of CCC.

Our PDE model used to generate benchmark simulation data is a flexible and customizable platform for modeling CCC in space. As one possible extension, gene regulatory networks, modeled by a system of ordinary differential equations, could be assigned to each cell, or coupled to each spot of cells to refine the PDE model for more accurate simulation datasets. Building such complex multiscale models is important and tractable as more multi-omics datasets become available, and the collective optimal transport can be integrated with the PDE models to better uncover detailed dynamics of CCC.

While traditional optimal transport is powerful at integrating a pair of datasets and multimarginal optimal transport^72^ integrates multiple datasets, the collective optimal transport is able to effectively deal with competing species. Moreover, collective optimal transport provides a new way to preserve units and control coupled mass. The collective optimal transport in this work will be useful for a broad range of problems beyond the CCC inference.

## Methods

Full details of the theoretical background and implementation of COMMOT can be found in Supplementary Information.

### COMMOT model

COMMOT constructs a collection of CCC networks through various predefined ligand-receptor pairs (user-defined or from aggregated ligand-receptor interaction databases) by solving a global optimization problem that accounts for the higher-order interactions among the multiple ligand and receptor species. To this end, we introduce collective optimal transport that seeks a collection of optimal transport plans for all pairs of species that can be coupled simultaneously. As a result, the coupling between a species pair will affect other couplings and vice versa, which cannot be realized in traditional optimal transport^34^. The collective optimal transport results in a large-scale optimization problem for which new algorithms are needed, and thus we presented one based on the efficient Sinkhorn iteration^73^.

For a ST data of *n*_*s*_ spatial locations and a set of *n*_*l*_ ligand species and *n*_*r*_ receptor species, a collective optimal transport problem is formulated as follows:

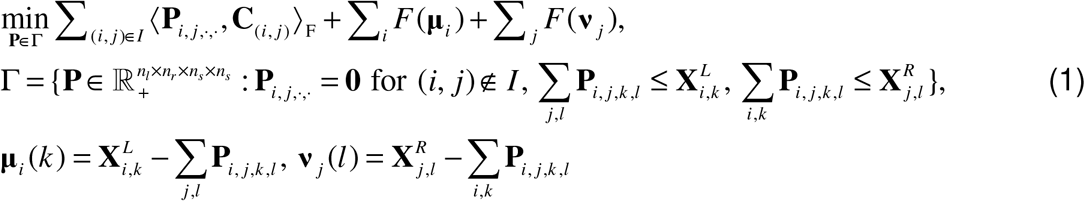

where 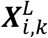 is the expression level of ligand *i* on spot *k*,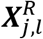 is the expression level of receptor *j* on spot *l*, and *F* penalizes the untransported mass **μ**_*i*_ and **ν**_*j*_. The coupling matrix **P**_*i,j,k*_ scores the signaling strength from spot *k* to spot *l* through the pair of ligands *i* and receptor *j* for (*i, j*) ϵ *I* where *I* is the index set of ligand and receptor species that can bind. The cost matrix ***C***_*(i,j*)_ is based on the thresholded distance matrix such that its *kl*-th entry equals *φ* (***D***_*k, l*_) if **D**_*k,l*_ > *T(*_*i, j*_) and infinity otherwise where **D** is the Euclidean distance matrix among the spots, *T*(_*i,j*_) is the spatial limit of signaling through the pair of ligand *i* and receptor *j*, and *φ* is a scaling function such as square or exponential. When the ligands or receptors contain heteromeric units, the minimum of units is used by default to represent the amount of ligand or receptor.

### Collective optimal transport algorithm

To solve the collective optimal transport problem described above, we rewrite the original problem as

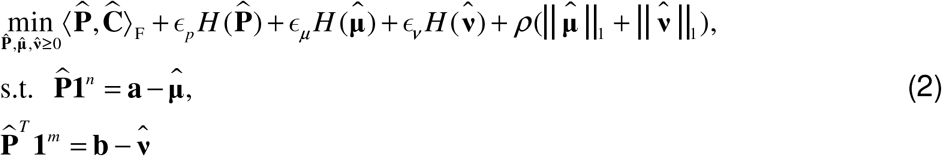

where 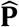 is obtained by reshaping **P** such that 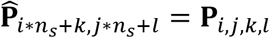. The cost matrix **Ĉ** is obtained similarly and 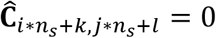 for ligand *i* and receptor *j* that cannot bind. The marginal distributions are constructed such that 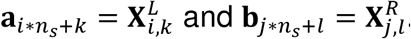. Entropy regularization is added to speed up computation and smooth the result with 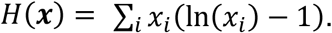.

When the entropy regularization terms have the same coefficient that *ϵ = ϵ*_*p*_ = *ϵ*_*μ*_ = *ϵ*_*ν*_, the problem can be efficiently solved with a stabilized Sinkhorn iteraction^73^

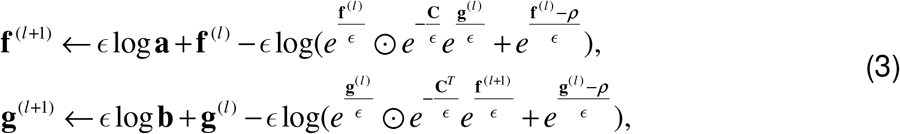

for *l* ≥ 0 with arbitrary initial **f**^(0)^ and **g**^(0)^. The resulting numerical solution to the optimization problem can be constructed by 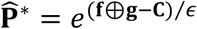. The formulation in Eq. (2) solved by the algorithm in Eq. (3) was used to generate the results in this study. The derivation of the algorithm and algorithms for the general case where the regularization terms have different coefficients are described in Supplementary Information.

### Spatial signaling direction

To visualize the spatial signaling directions, we estimate a spatial vector field 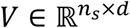 of signaling directions given a CCC matrix 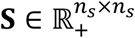 obtained from collective optimal transport algorithm where **S**_*i, j*_ is the strength of signal sent by spot *i* to spot *j*. The ith row of *V* represents the spatial signaling direction. We construct two vector fields, **V**^*s*^ and **S**^*r*^ describing to/from which directions the spots are sending/receiving signals, respectively. Specifically, 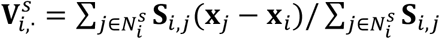, where 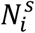 is the index set of top *k* signal receiving spots with the largest value on the *i*th row of **s**. Similarly, 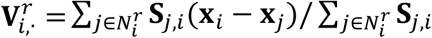, where 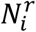 is the index set of top *k* signal sending spots with the largest value on the *i*th column of **S**.

### Cluster level CCC

To elucidate CCC among cell states or local groups of spots, we summarize the spot-by-spot CCC matrix **S** to a cluster-by-cluster matrix **S**^*cl*^. The signaling strength from cluster *i* to cluster *j* is quantified as 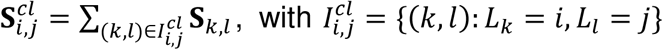 where *L*_*k*_ is the cluster label of spot *k*. The significance (*p*-value) of the cluster level CCC is determined by performing *n* independent permutations of cluster labels and computing the percentile of the original signaling strength among the signaling strengths resulted from these label permutations. Only permutating cluster labels in this spatial context might neglect communications between different clusters. To address this limitation, we provide an option that randomly permutates the locations of all spots or the spots within each cluster.

### Evaluation metrics

The spatial signaling direction is described by a vector field defined on *n* grid points discretizing the tissue space, and is represented by an array **V** ϵ ℝ ^*n* ×2^. Cosine distance is used to compare the vector field **V**_sub_ from subsampled data to the one from the full data **V**_full_ and is defined as *d*_*cos*_ (**V**_full_,**V**_sub_) = Σ_i_ ∥ **V**_full_(*i*) ∥ [1-**V**_full_(*i*). **V**_sub_(*i*)/(∥ **V**_full_(*i*) ∥ ∥ **V**_sub_ (*i*)] Σ_*i*_ ∥ **V**_full_ (*i*) ∥.

To compare two cluster level CCC networks 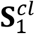 and 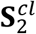, we first binarize them such that the edges with *p*-value smaller than are kept in the edge sets 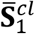 and 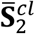. Then, the Jaccard distance is used for quantitative comparison, 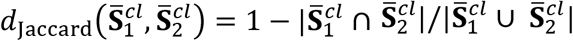.

Spearman’s correlation coefficient was used to quantify the correlation between the inferred signaling activity. The average expression of the known target genes across the cell clusters defined as 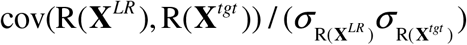, where 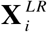 is the average received signal through a ligand-receptor pair in cell cluster *i*, and 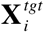 is the average expression of the known target genes of this ligand-receptor pair in cell cluster *i*. The function R converts the vectors into ranks and *σ* is the standard deviation of the rank variables.

### Downstream gene analysis

After computing the CCC matrix **S** of a ligand-receptor pair or a signaling pathway, genes that are potential downstream targets whose expression are regulated by the corresponding CCC can be identified. The amount of signal received by each spot is quantified by 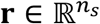 where ***r***_*i*_ = Σ_*j*_**S**_*j,i*_. Then the tradeSeq package^57^ is used to identify the genes that are differentially expressed with respect to **r** which we call differentially expressed CCC genes.

The identified differentially expressed CCC genes may be regulated by other genes within cells through gene regulation. To further prioritize the downstream genes whose expressions are affected by CCC, we train a random forest regression model^58,59^ that takes a potential downstream gene as the output and **r** and a collection of highly correlated genes as input features. The unique impact of CCC on this potential downstream gene is quantified by the feature importance (Gini importance computed as the mean of total impurity decrease within each tree) of **r** in the trained random forest model. The inclusion of highly correlated genes within a cell as input features emphasizes the amount of information of potential target genes explained by inferred CCC which is unlikely explained only by intracellular interactions. If such dilution of importance is not preferred, the users may choose a smaller number of highly correlated genes as input features. The implementation in scikit-learn package^59^ is used.

### CellChat, Giotto, and CellPhoneDB analysis

In the CellChat analysis, the spatial data was treated as a scRNA-seq data, and the count matrix was first normalized using the normalizeData function. The data was then filtered using functions identifyOverExpressedGenes and identifyOverExpressedInteractions with the default parameters. The cluster-level communication scores in CellChat were computed using the computeCommunProb function with default parameters and the results were further filtered using the filterCommunication function with min.cells set to 10. The ligand-receptor pairs categorized as secreted signaling in CellChatDB were examined. In the Giotto analysis, the count data was first normalized using normailizeGiotto function with default parameters. A spatial network was then created using the createSpatialNetwork function with the knn method and k set to 10. The heteromeric ligand-receptor pairs in CellChatDB were converted to pairs of individual subunits. The spatCellCellcom function was then used to generate the cluster-level communication scores with the adjust_method set to fdr. In the CellPhoneDB v3 analysis, the distance between clusters is quantified as the average distance between cells from the pair of clusters. The commend “cellphonedb method statistical_analysis” was used to generate CellPhoneDB results with the threshold parameter set to 0.1.

### Immunostaining and fluorescent *in situ* hybridization

Frozen tissue sections (10μm) were fixed with 4% PFA in PBS for 15 minutes. 10% BSA in PBS was used for blocking. Following blocking, 5% BSA and 0.1% Triton X-100 in PBS was used for permeabilization. The following antibodies were used: mouse anti-KRT5 (1:100; Santa Cruz Biotechnilogy; sc-32721), mouse anti-KRT15 (1:100; Santa Cruz Biotechnology; sc-47697), mouse anti-BCAM (1:100; Santa Cruz Biotechnology; sc-365191), mouse anti-FGF7 (1:100; Santa Cruz Biotechnology; sc-365440), mouse anti-STMN1 (1:100; Santa Cruz Biotechnology; sc-48362); mouse anti-IGFBP6 (1:500; Abgent; AP6764b); mouse anti-PMAIP1 (1:100; Santa Cruz Biotechnology; sc-56169), mouse anti-POSTN (1:100; Santa Cruz Biotechnology; sc-398631); mouse anti-FLG (1:100; Santa Cruz Biotechnology; sc-66192); and rabbit anti-LOR (1:1000; abcam; ab85679). Secondary antibodies include Cy3 AffiniPure (1:500; Jackson ImmunoResearch; 711-165-152, 111-165-003). Slides were mounted with Prolong Diamond Antifade Mountant containing DAPI (Molecular Probes;). Confocal images were acquired at room temperature on a Zeiss LSM700 laser scanning microscope with Plan-Apochromat 20x objective or 40x and 63x oil immersion objectives.

Frozen neonatal human foreskin tissue sections were used for RNA in situ hybridization using RNAscope® kit v2 (323100, Advanced Cell Diagnostics), following the manufacturer’s instructions. The following Homo sapiens probes from Advanced Cell Diagnostics were used: Tyro3 probe (429611), Gas6 (427811-C2), and Pros1 (506991-C2). Confocal images were acquired at room temperature on an Olympus FV3000 confocal microscope with Plan-Apochromat 20x objective or 40x and 60x oil immersion objectives.

## Data Availability

The original public data used in this work can be accessed through the following links: (1) *Drosophila* embryo spatial and scRNA-seq data: Dream Single cell Transcriptomics Challenge through Synapse ID (syn15665609)^64^. (2) Human epidermal scRNA-seq data^40^: GEO assession codes, GSE147482. (3) Mouse hypothalamic preoptic region MERFISH data^45^: original data available at Dryad^74^ through the link https://doi.org/10.5061/dryad.8t8s248. This work used the preprocessed data through the Squidpy package^22^ with the utility squidpy.datasets.merfish. (4) Mouse placenta STARmap data^48^: downloaded from Code Ocean (https://codeocean.com/capsule/9820099/tree/v1). (5) Mouse brain STARmap data^20^: the processed data was downloaded from the same repository as the mouse placenta STARmap data. (6) Mouse somatosensory cortex seqFISH+ data^18^: downloaded through Giotto package^23^. (7) Mouse hippocampus Slide-seqV2 data^56^: downloaded from Broad Institute Single Cell Portal (https://singlecell.broadinstitute.org/single_cell/study/SCP815/sensitive-spatial-genome-wide-expression-profiling-at-cellular-resolution#study-summary). (8) Breast cancer Visium data: downloaded from 10X Genomics website (https://www.10xgenomics.com/resources/datasets/human-breast-cancer-block-a-section-1-1-standard-1-1-0). (9) Mouse brain (sagittal posterior) Visium data: downloaded from 10X Genomics website (https://www.10xgenomics.com/resources/datasets/mouse-brain-serial-section-1-sagittal-anterior-1-standard-1-1-0).

The ligand-receptor pairs with secreted ligand based on CellChatDB database^6^ were used. The downstream target genes were taken from scSeqComm^65^ and the target gene libraries named TF_TG_TRRUSTv2 and TF_TG_TRRUSTv2_RegNetwork_High_mouse were used for human and mouse respectively.

## Code Availability

The open-source software is available at https://github.com/zcang/COMMOT.

## Acknowledgements

This work was supported by two NSF grants DMS1763272 and CBET2134916, a grant from Simons Foundation (594598, QN), and two NIH grants U01AR073159 and R01DE030565. Z. Cang’s work was partially supported by a startup grant from NCSU and NSF grant DMS2151934. Y. Zhao’s work was supported by a grant from the Simons Foundation through Grant No. 357963 and NSF grant DMS2142500. Z. Cang thanks Wei Zhao for helpful discussion.

## Author Contributions

Z.C., Y.Z., and Q.N. conceived the method. Z.C. implemented the method. Z.C. and A.A.A. generated the numerical results. R.R., A.S., and S.X.A. generated the experimental results. Z.C., R.R., A.S., M.P., S.X.A., and Q.N. interpreted the results, generated the visualizations, and wrote the paper. All authors reviewed the manuscript.

## Competing Interests

The authors declare no competing interests.

## Notes

### Competing Interest Statement

The authors have declared no competing interest.

