## Supplementary material for "Screening cell-cell communication in spatial transcriptomics via collective optimal transport"

### Supplementary Figures

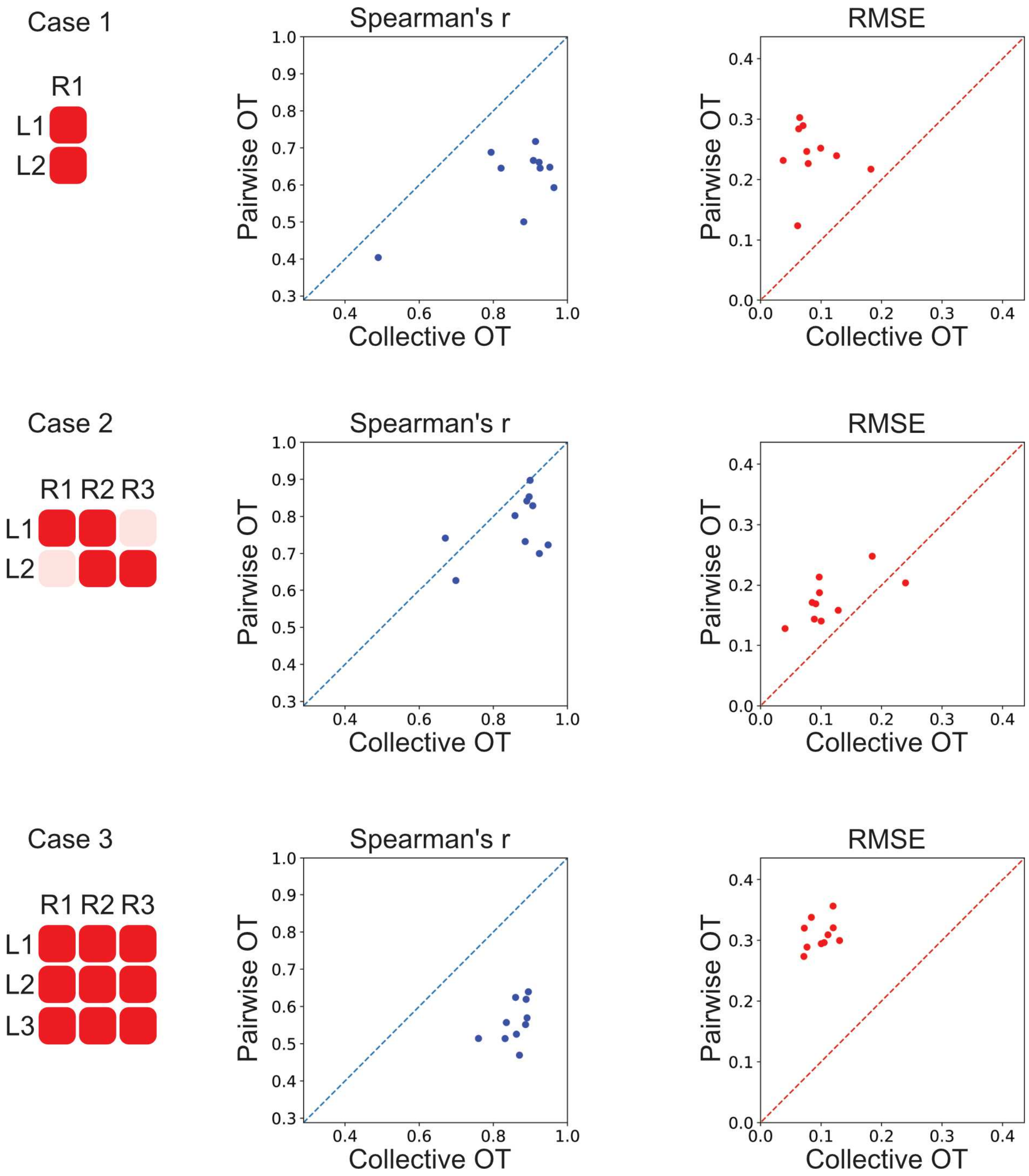

**Supplementary Figure 1**

Performance of collective optimal transport (benchmark cases 1-3)

The performance of collective optimal transport and traditional pairwise optimal transport obtained by comparison with benchmarks (cases 1-3) from PDE simulations. Each benchmark case has a specified number of ligand species and receptor species and a specific binding pattern among the species. The dark red color annotates the ligand-receptor pairs that can bind. The performance is measured by root-mean-squared error (RMSE) and Spearman's correlation coefficient among the grid points.

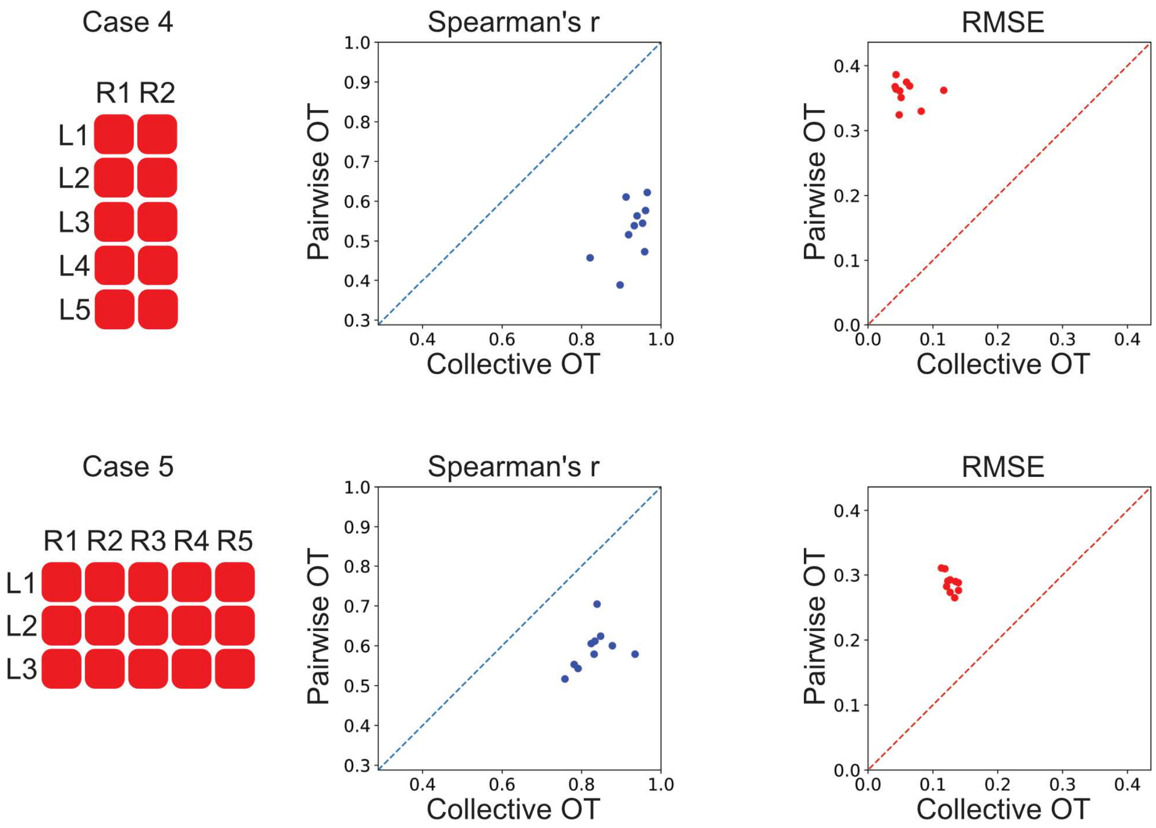

**Supplementary Figure 2**

Performance of collective optimal transport (benchmark cases 4-5)

The performance of collective optimal transport and traditional pairwise optimal transport obtained by comparison with benchmarks (cases 4-5) from PDE simulations. Each benchmark case has a specified number of ligand species and receptor species and a specific binding pattern among the species. The dark red color annotates the ligand-receptor pairs that can bind. The performance is measured by root-mean-squared error (RMSE) and Spearman's correlation coefficient among the grid points.

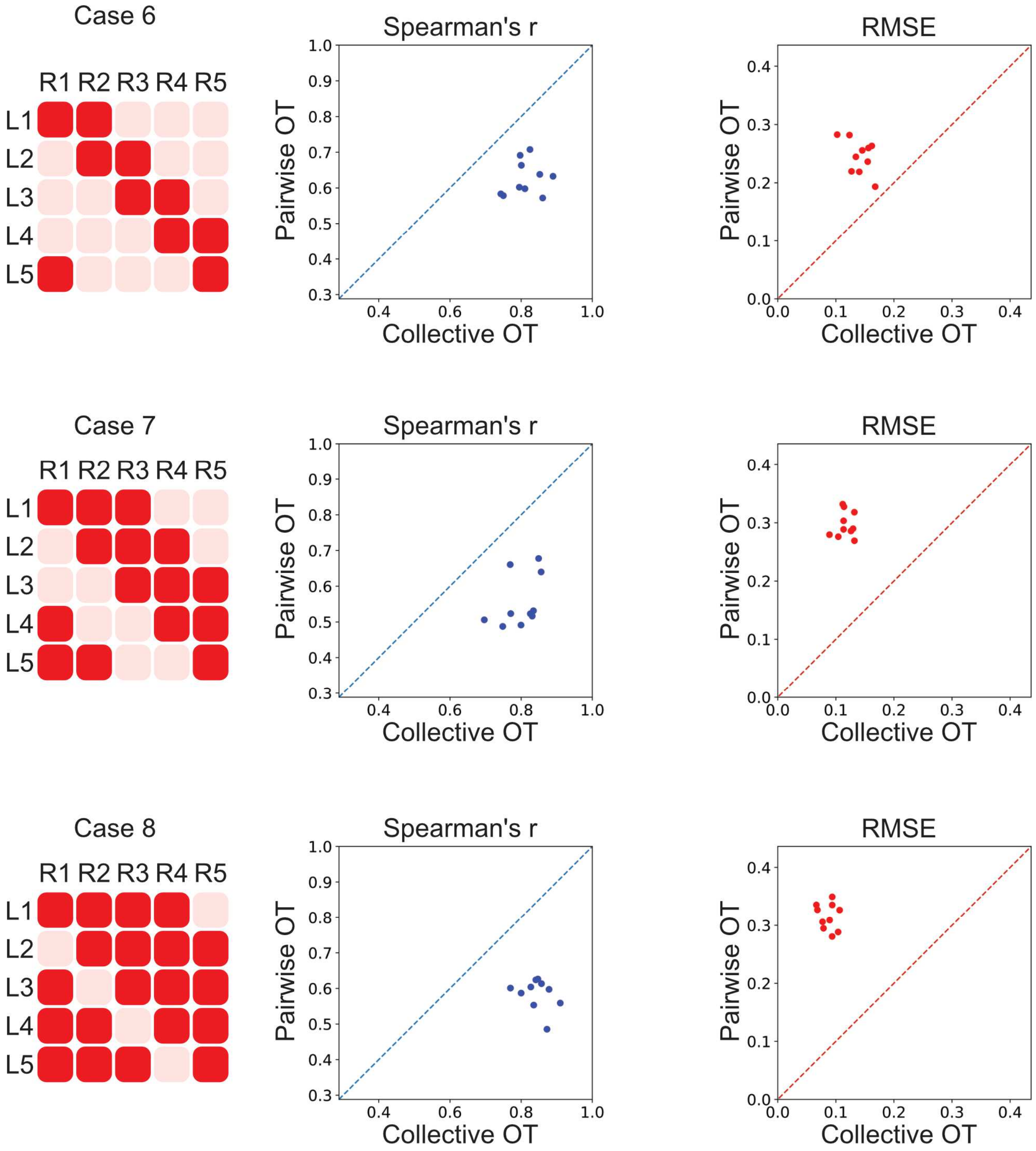

**Supplementary Figure 3**

Performance of collective optimal transport (benchmark cases 6-8)

The performance of collective optimal transport and traditional pairwise optimal transport obtained by comparison with benchmarks (cases 6-8) from PDE simulations. Each benchmark case has a specified number of ligand species and receptor species and a specific binding pattern among the species. The dark red color annotates the ligand-receptor pairs that can bind. The performance is measured by root-mean-squared error (RMSE) and Spearman's correlation coefficient among the grid points.

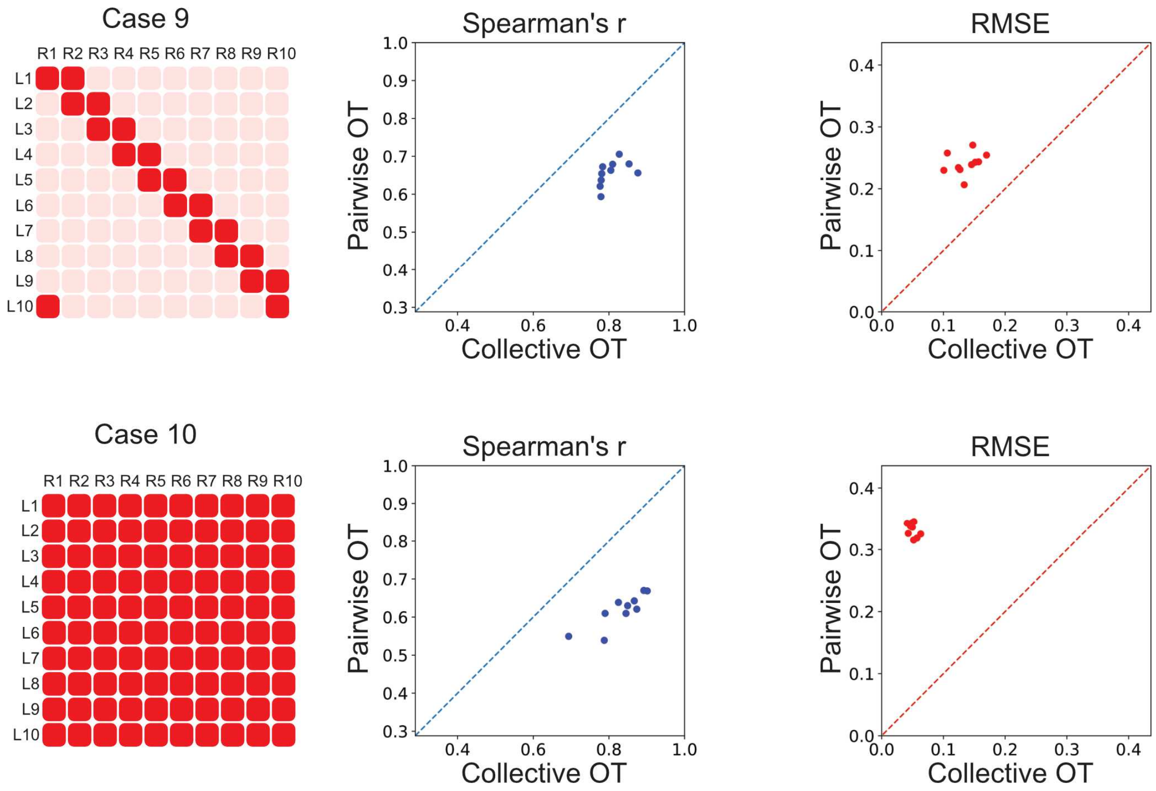

#### Supplementary Figure 4

Performance of collective optimal transport (benchmark cases 9-10)

The performance of collective optimal transport and traditional pairwise optimal transport obtained by comparison with benchmarks (cases 9-10) from PDE simulations. Each benchmark case has a specified number of ligand species and receptor species and a specific binding pattern among the species. The dark red color annotates the ligand-receptor pairs that can bind. The performance is measured by root-mean-squared error (RMSE) and Spearman's correlation coefficient among the grid points.

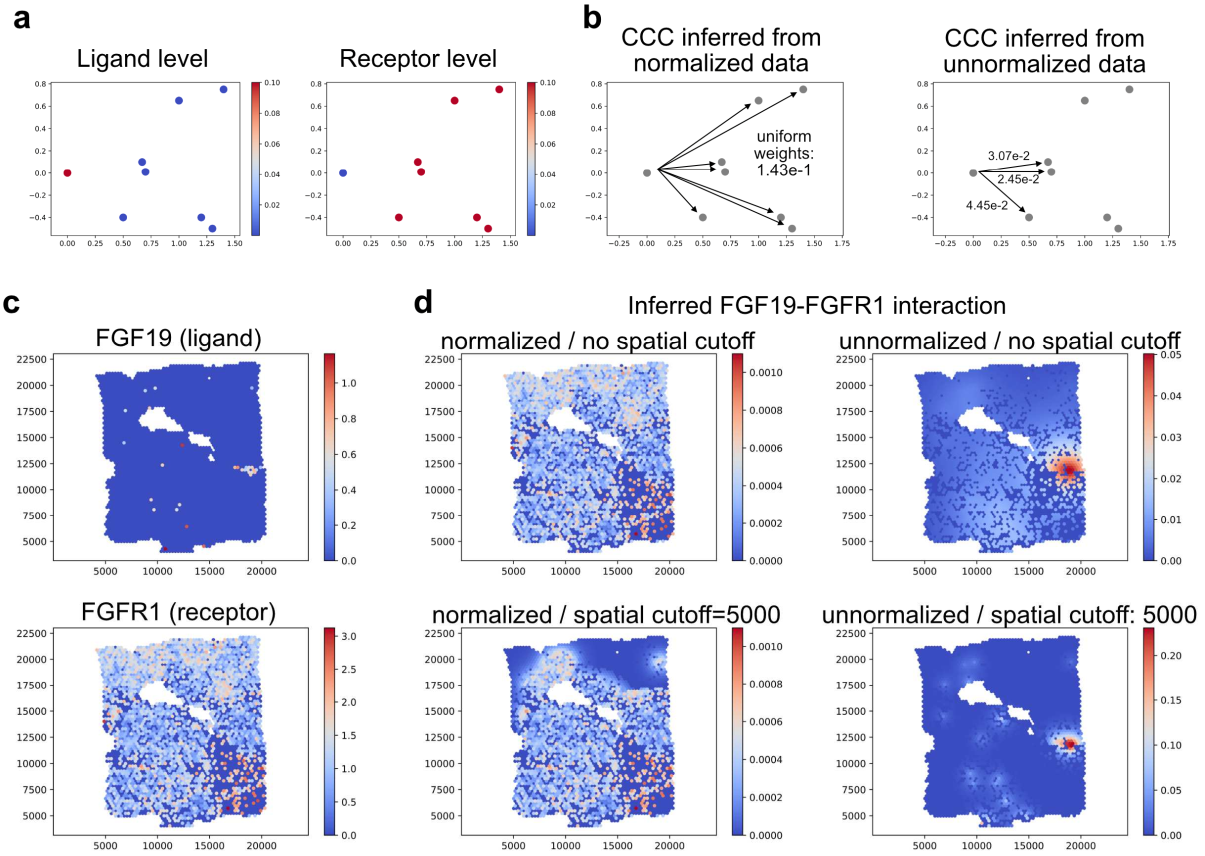

#### Supplementary Figure 5

Comparison of whether to normalize expression to probability distributions.

**a** An example with 8 cells where 1 cell expresses the ligand gene and the other 7 cells express the receptor gene. **b** The inferred cell-cell communication (CCC) with or without normalizing the expression levels to probability distributions. **c** An example ligand-receptor pair in a Visium human breast cancer data. **d** The inferred spatial distribution of received signals through the ligand-receptor pair FGF19-FGFR1 with or without normalization to probability distribution and with or without a spatial distance cutoff.

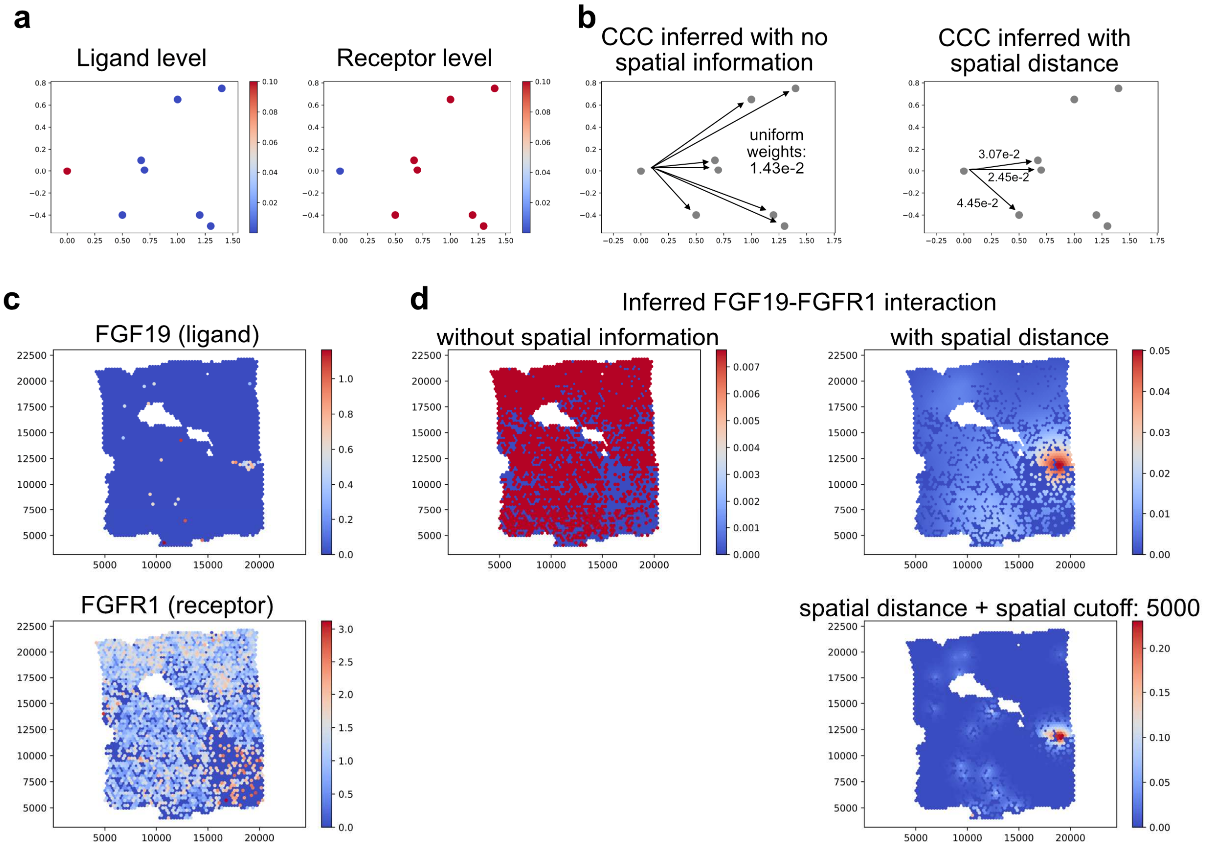

#### Supplementary Figure 6

Comparison of whether to incorporate spatial distance information.

**a** An example with 8 cells where 1 cell expresses the ligand gene and the other 7 cells express the receptor gene. **b** The inferred cell-cell communication (CCC) with or without normalizing the expression levels to probability distributions. Both results are from the spatial expression levels without normalizing to probability distributions. **c** An example ligand-receptor pair in a Visium human breast cancer data. **d** The inferred spatial distribution of received signals through the ligand-receptor pair FGF19-FGFR1 with or without incorporating spatial information and if spatial information is used, with or without a spatial distance cutoff.

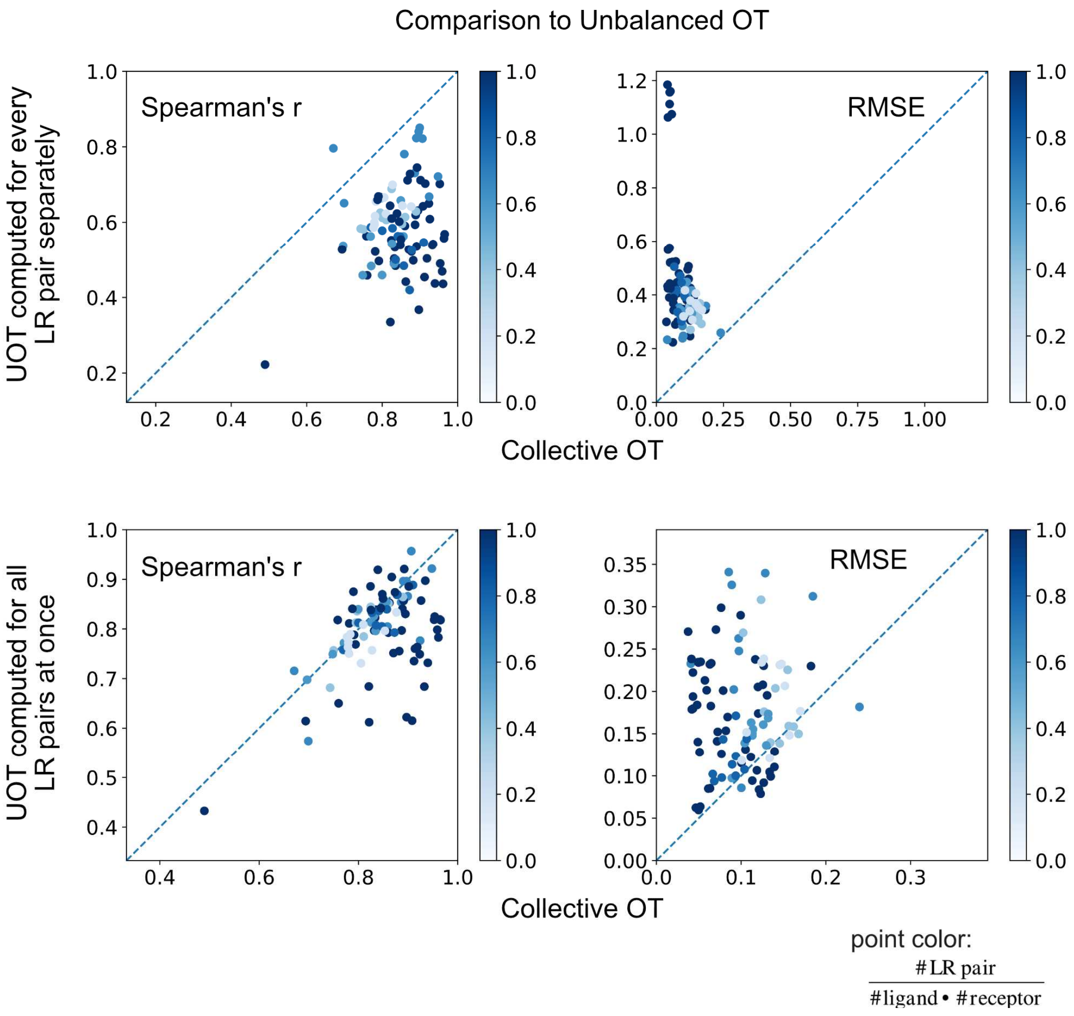

#### Supplementary Figure 7

##### Comparison with unbalanced optimal transport

The comparisons with unbalanced optimal transport that examines each ligand-receptor pair separately (top) or all pairs together (bottom). All analyses use the same spatial cutoff such that the transport cost is set to infinity if the distance between a location pair exceeds the distance cutoff. The optimal transport analysis results are evaluated by comparing the inferred amount of received signal to those obtained by simulation using Spearman’s correlation coefficient and root-mean-squared error.

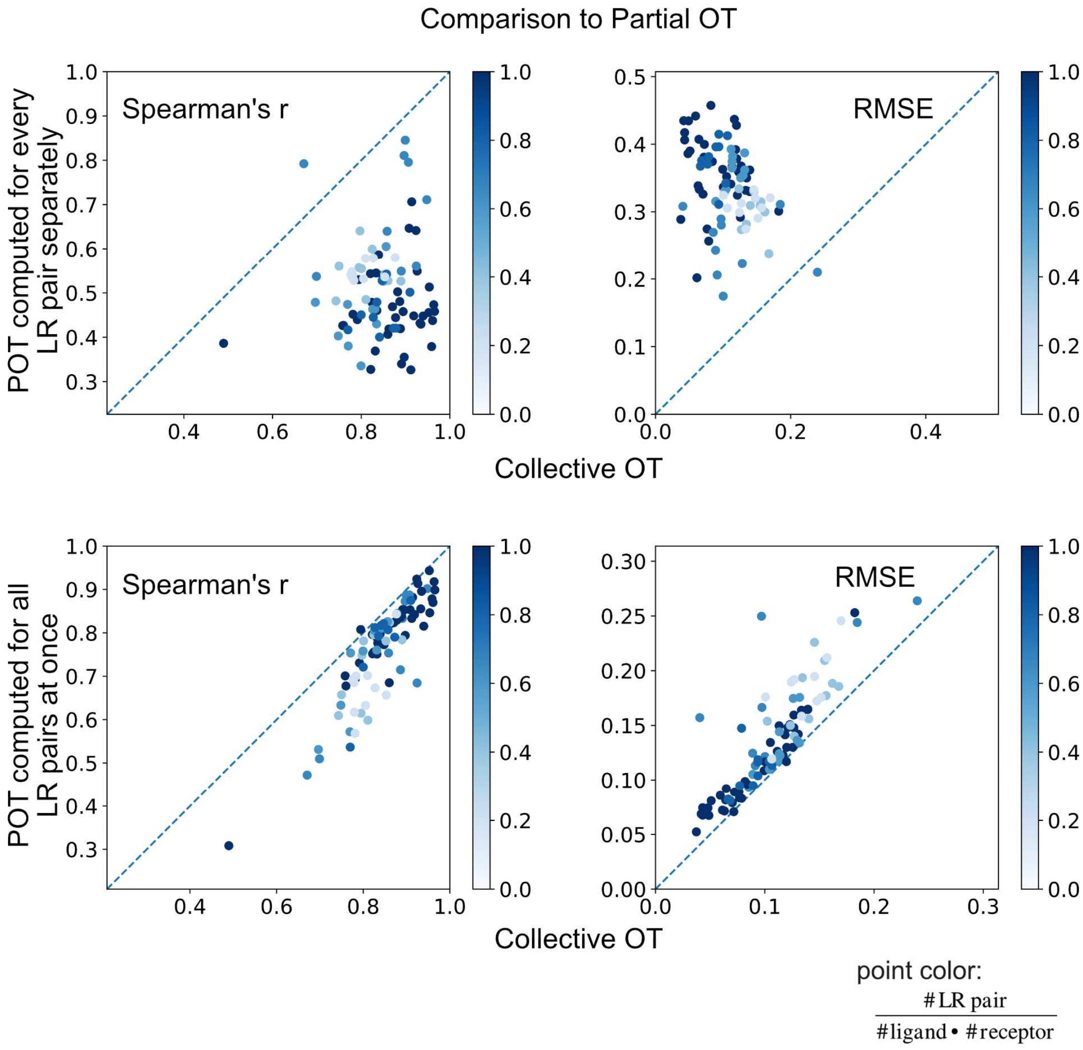

#### Supplementary Figure 8

##### Comparison with partial optimal transport

The comparisons with partial optimal transport that examines each ligand-receptor pair separately (top) or all pairs together (bottom). All analyses use the same spatial cutoff such that the transport cost is set to infinity if the distance between a location pair exceeds the distance cutoff. The optimal transport analysis results are evaluated by comparing the inferred amount of received signal to those obtained by simulation using Spearman’s correlation coefficient and root-mean-squared error.

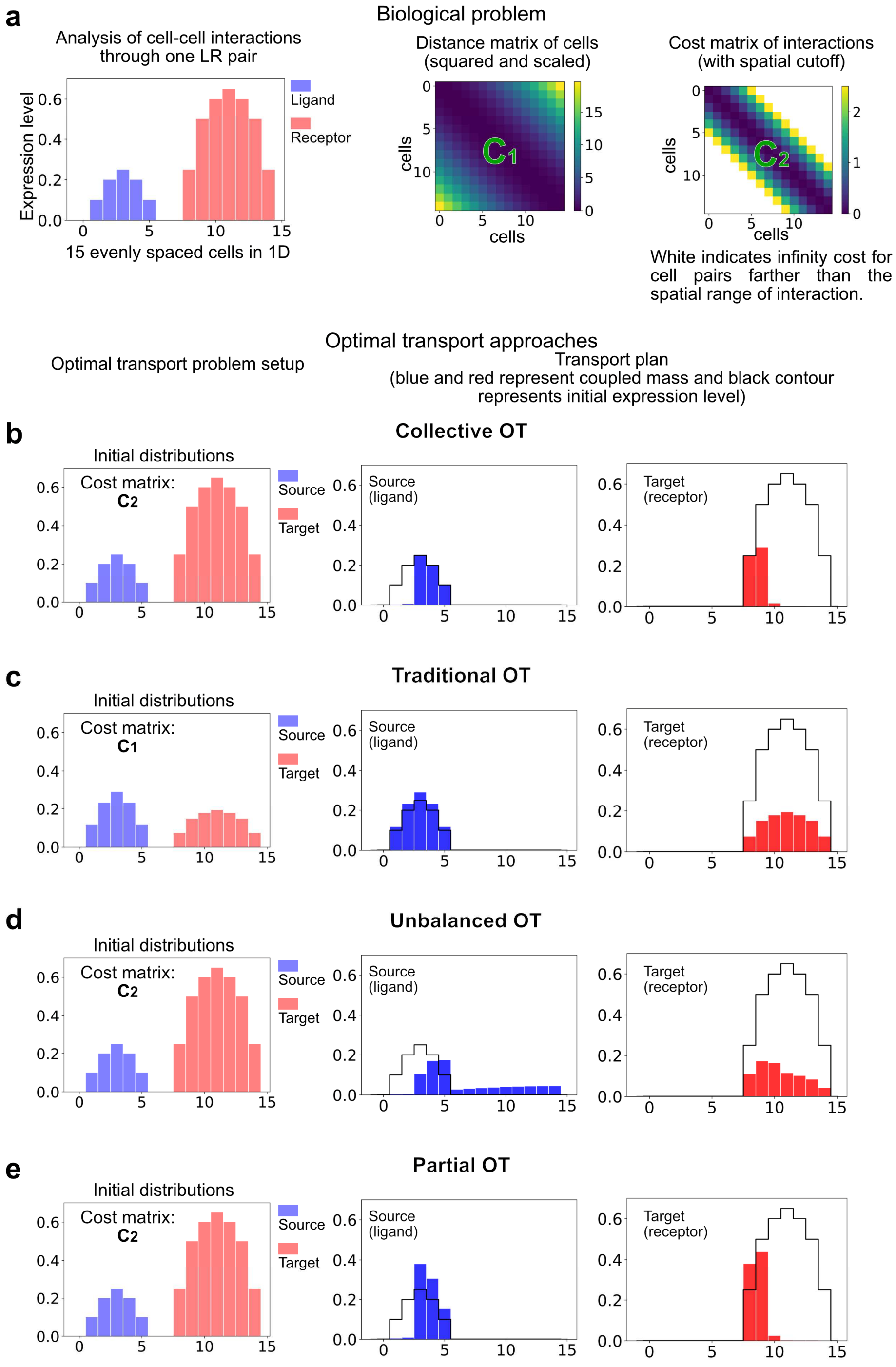

#### Supplementary Figure 9

##### Illustration of differences among several optimal transport variants

**a** The problem of fifteen cells placed one unit apart in 1D through one ligand-receptor pair with a spatial range cutoff of five units. **b-e** The input distributions and cost matrices of each optimal transport approach and the marginal distributions of the optimal transport plans. All three variants (collective OT, unbalanced OT, and partial OT) can take the initial unnormalized distributions and the cost matrix with infinity entries. However, unbalanced OT and partial OT result in artificial extra amount of ligand and receptor exceeding their expression levels. Traditional OT also induces artificial expressions due to normalization and cannot enforce the spatial range of signaling. Collective OT controls the coupled mass by the initial distributions and enforces the spatial range of signaling.

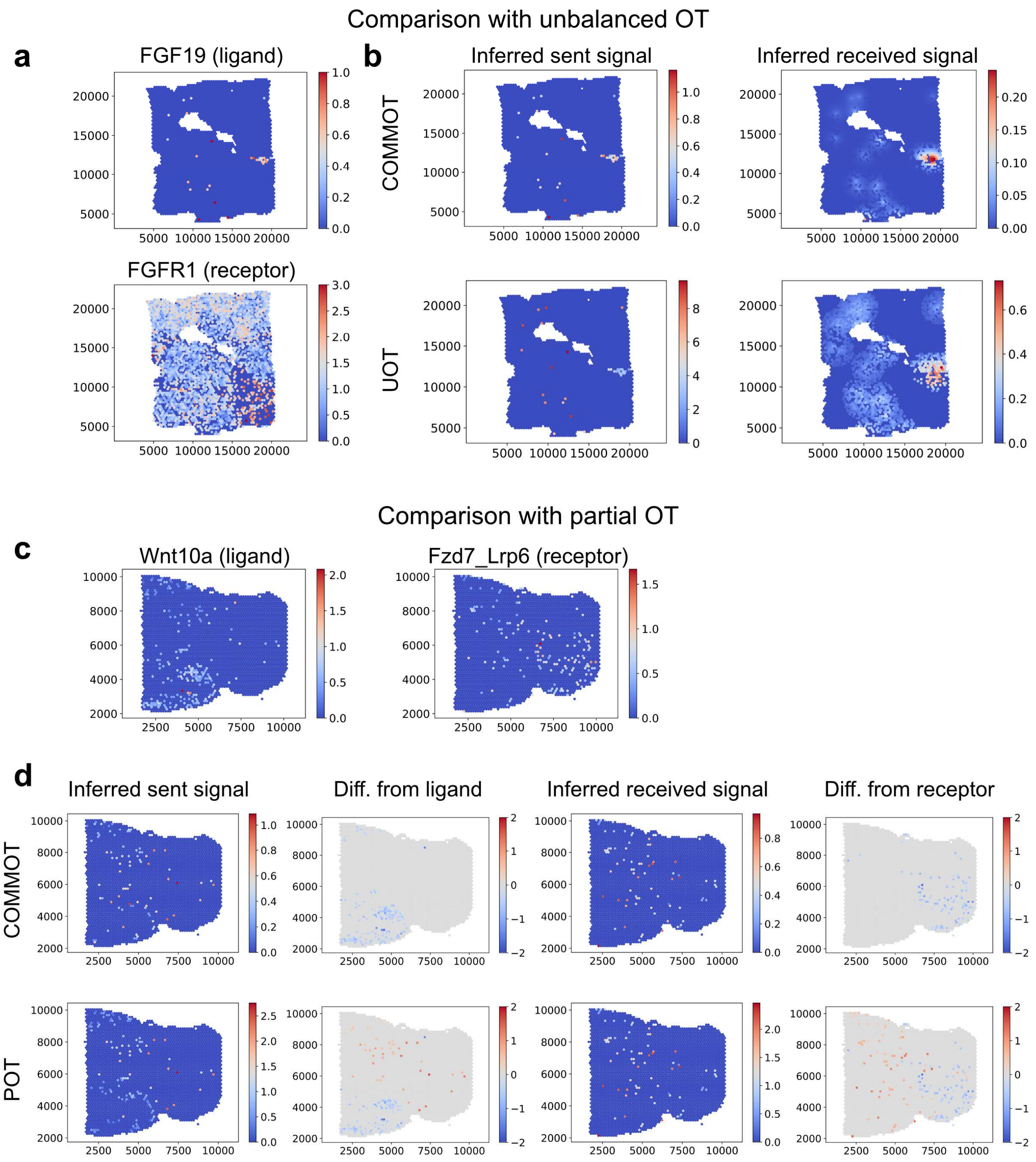

#### Supplementary Figure 10

Differences between OT variants on example Visium datasets

**a** The difference between COMMOT and unbalanced OT is demonstrated with a Visium dataset of human breast cancer focusing on the ligand-receptor pair FGF19-FGFR1 whose total expression levels are significantly different. **b** The amount of sent and received signal is plotted. Compared to COMMOT, unbalanced OT overestimates the signaling activity. Specifically, the amount of sent signal inferred using unbalanced OT significantly exceeds the expression level of ligand. **c** The difference between COMMOT and partial OT is demonstrated with a Visium dataset of mouse brain focusing on the ligand-receptor pair Wnt10a-Fzd7_Lrp6 whose total expression levels are comparable but are expressed in spatially distinct regions. **d** The amounts of sent, received signal and their difference to the expression levels of ligand and receptor are plotted. Compared to partial OT, COMMOT further prioritizes potential signal senders and receivers. The difference plots show that the sent and received signal inferred using partial OT exceed ligand and receptor expressions in some regions leading to overestimation of signaling activity.

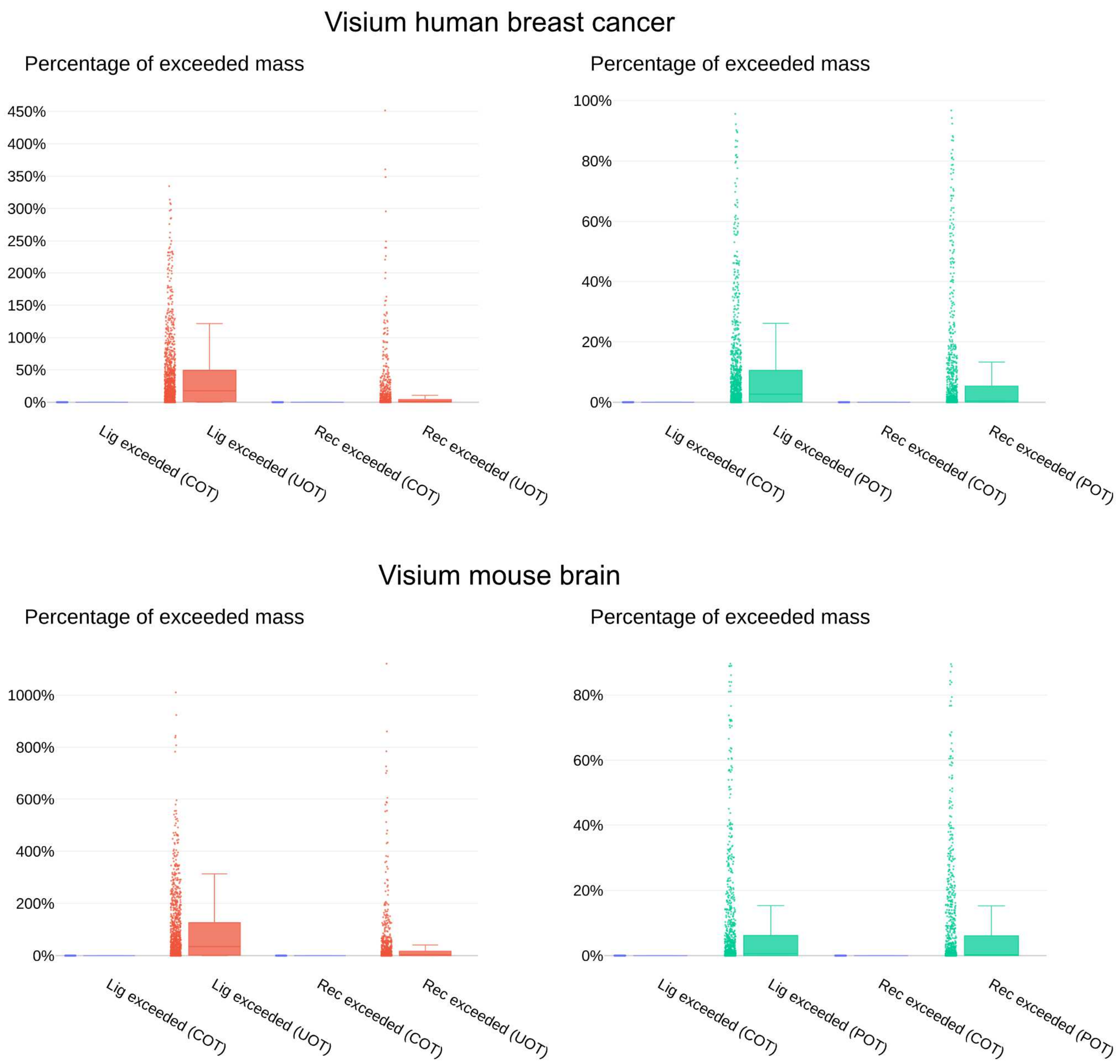

**Supplementary Figure 11**

Inferred sent or received signal exceeding the amount of ligand or receptors

Each dot represents a ligand-receptor pair. The commonly used KL divergence-based unbalanced optimal transport is used.

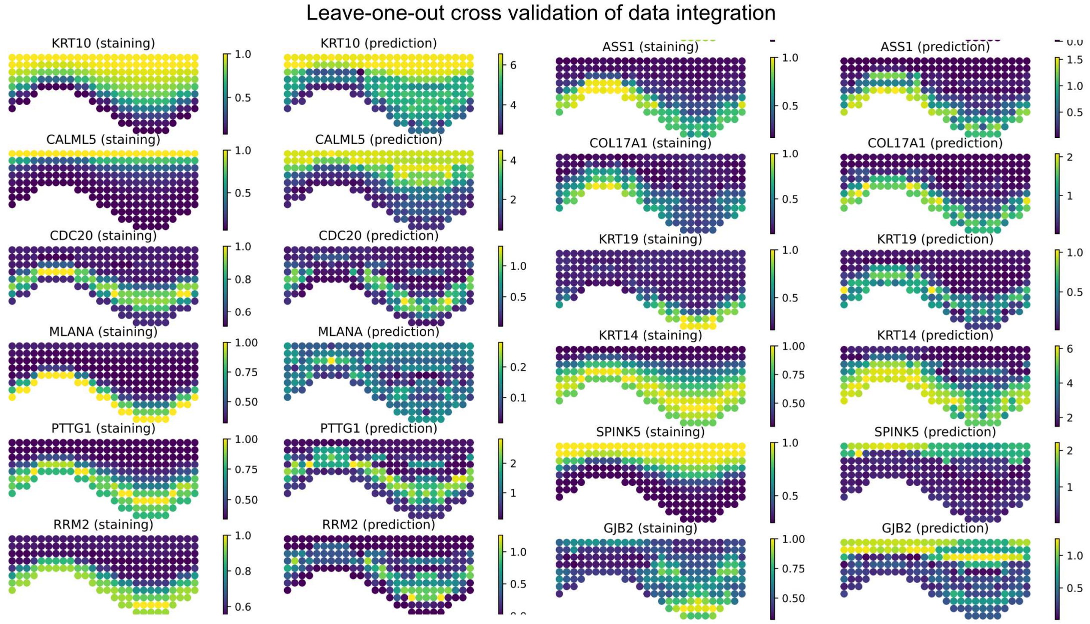

#### Supplementary Figure 12

Leave-one-out cross validation of scRNA-seq and spatial data integration for the human epidermis data

For each gene in the spatial data, the integration of scRNA-seq and spatial data is performed using the rest of the genes and the predicted spatial expression of this gene is compared to the ground-truth.

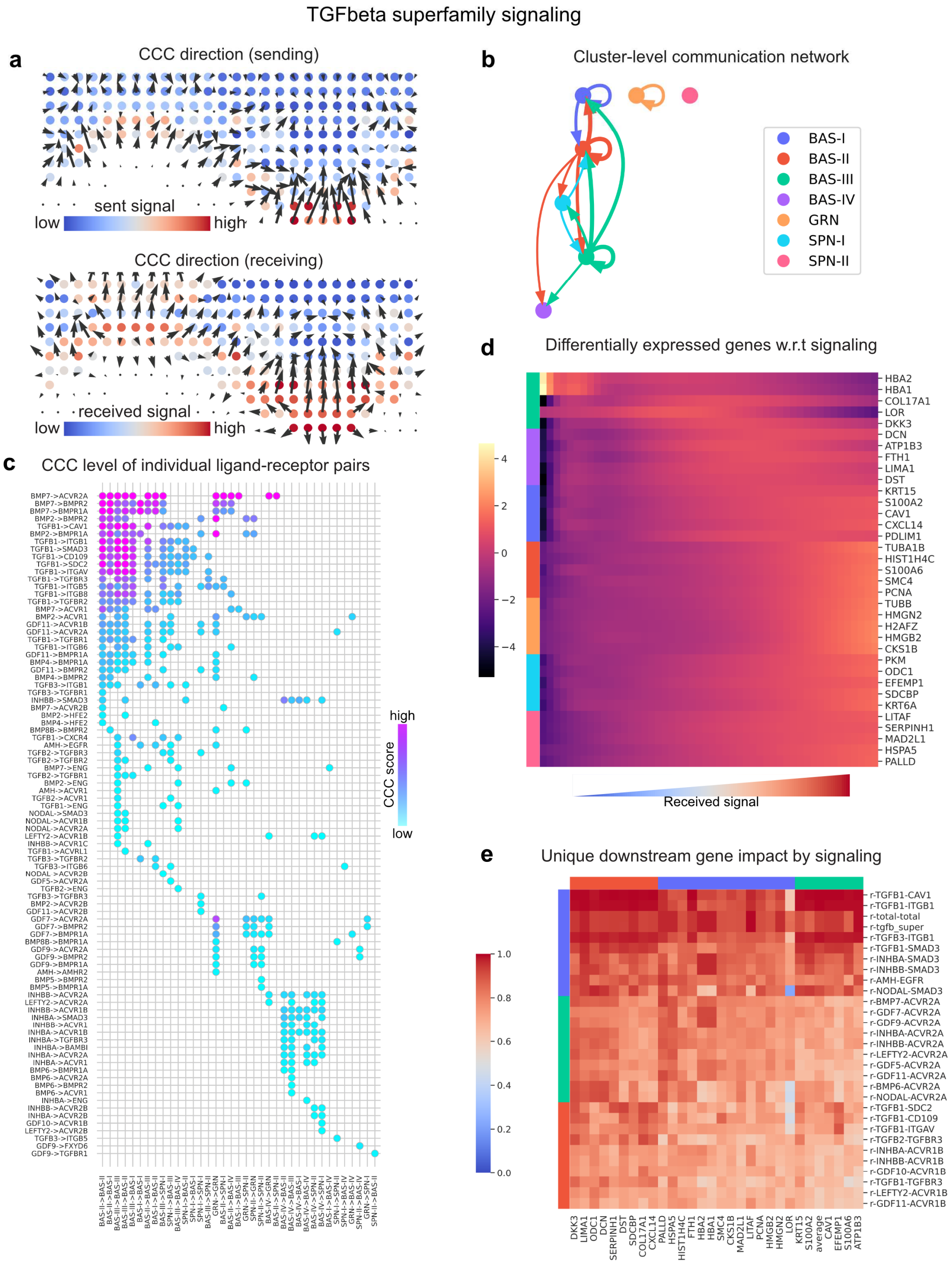

### Supplementary Figure 13

#### TGFβ superfamily signaling in human skin

**a** Cell-cell communication directions. Sender: the direction shows to where the signal sending cells are sending the signal and color depicts the amount of signal sent by each cell. Receiver: the direction shows from where the signaling receiving cells are receiving the signal and color depicts the amount of signal received by each cell. **b** The cell-cell communication of all ligand-receptor pairs summarized to clusters (clustering identical to the recent NC paper), a thicker arrow means stronger signaling activity. **c** Dotplot showing the detailed signaling strength among the clusters through the individual ligand-receptor pairs. **d** The identified differentially expressed genes due to cell-cell communication activity. This is analogous to pseudotime DE genes but with the horizontal axis describing the amount of received signal. **e** Unique impact of CCC on downstream genes. A high score means the ligand-receptor pair is likely affecting the expression of the downstream gene considering the effects from other genes in the same cell.

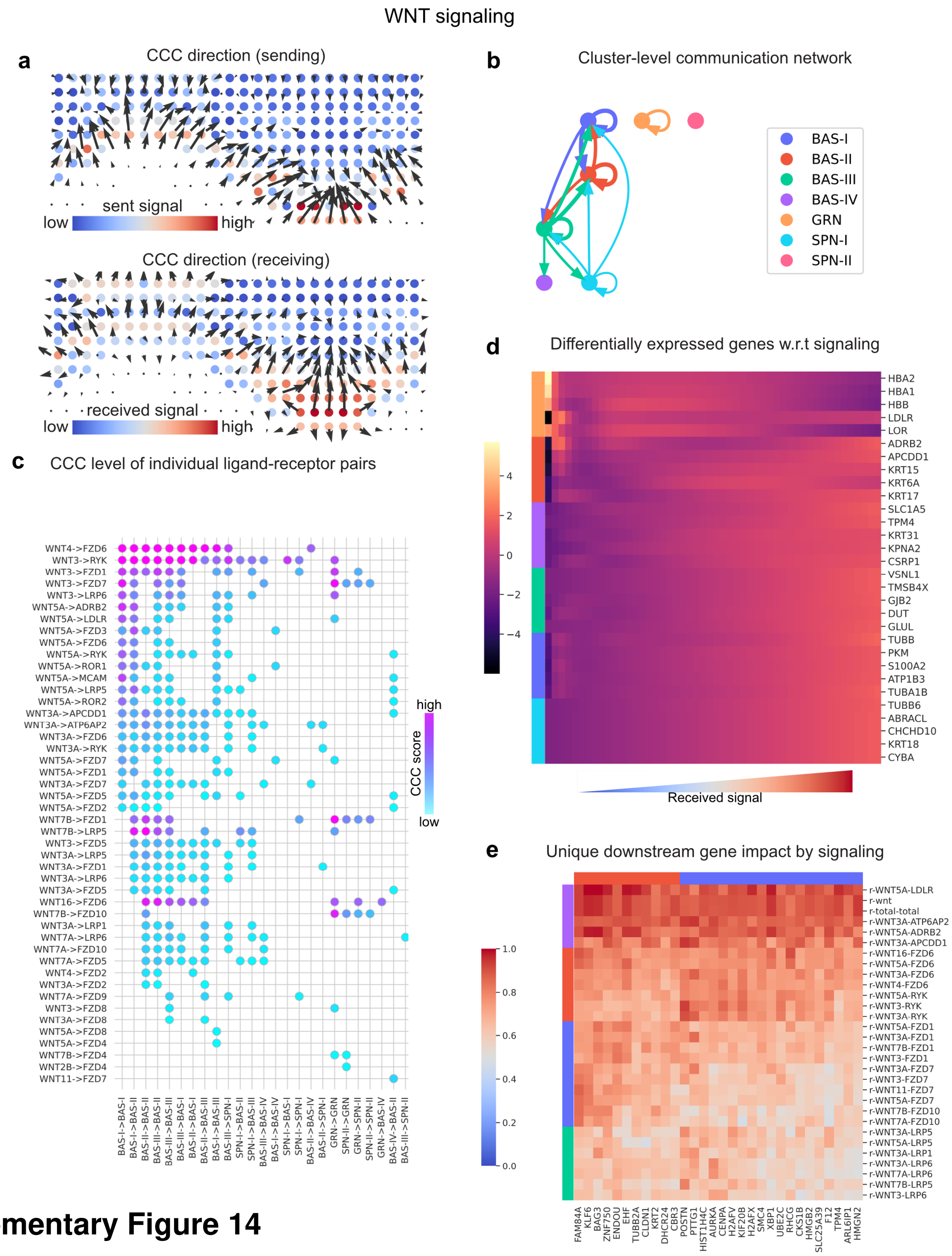

**Supplementary Figure 14**

WNT signaling in human skin

**a** Cell-cell communication directions. Sender: the direction shows to where the signal sending cells are sending the signal and color depicts the amount of signal sent by each cell. Receiver: the direction shows from where the signaling receiving cells are receiving the signal and color depicts the amount of signal received by each cell. **b** The cell-cell communication of all ligand-receptor pairs summarized to clusters (clustering identical to the recent NC paper), a thicker arrow means stronger signaling activity. **c** Dotplot showing the detailed signaling strength among the clusters through the individual ligand-receptor pairs. **d** The identified differentially expressed genes due to cell-cell communication activity. This is analogous to pseudotime DE genes but with the horizontal axis describing the amount of received signal. **e** Unique impact of CCC on downstream genes. A high score means the ligand-receptor pair is likely affecting the expression of the downstream gene considering the effects from other genes in the same cell.

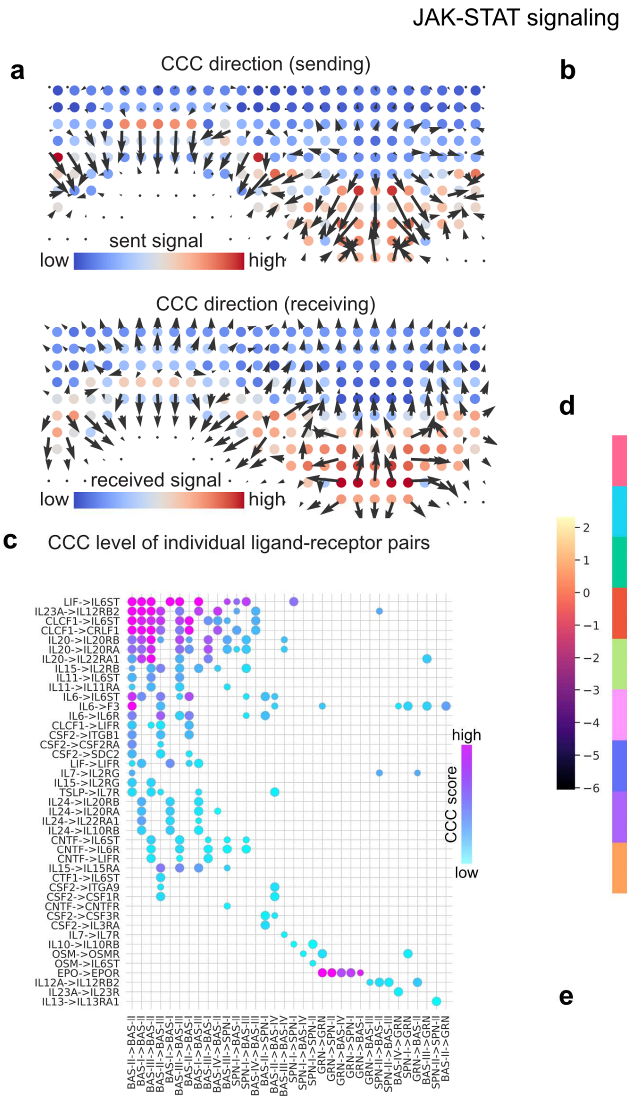

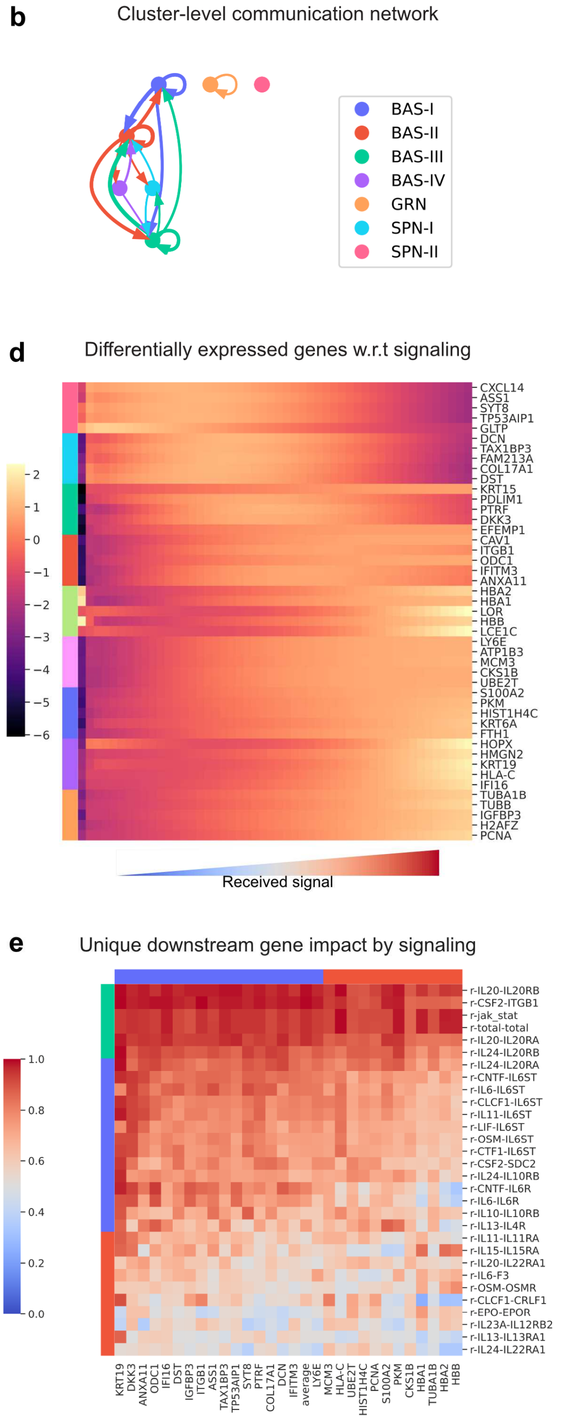

#### Supplementary Figure 15

#### JAK-STAT signaling in human skin

**a** Cell-cell communication directions. Sender: the direction shows to where the signal sending cells are sending the signal and color depicts the amount of signal sent by each cell. Receiver: the direction shows from where the signaling receiving cells are receiving the signal and color depicts the amount of signal received by each cell. **b** The cell-cell communication of all ligand-receptor pairs summarized to clusters (clustering identical to the recent NC paper), a thicker arrow means stronger signaling activity. **c** Dotplot showing the detailed signaling strength among the clusters through the individual ligand-receptor pairs. **d** The identified differentially expressed genes due to cell-cell communication activity. This is analogous to pseudotime DE genes but with the horizontal axis describing the amount of received signal. **e** Unique impact of CCC on downstream genes. A high score means the ligand-receptor pair is likely affecting the expression of the downstream gene considering the effects from other genes in the same cell.

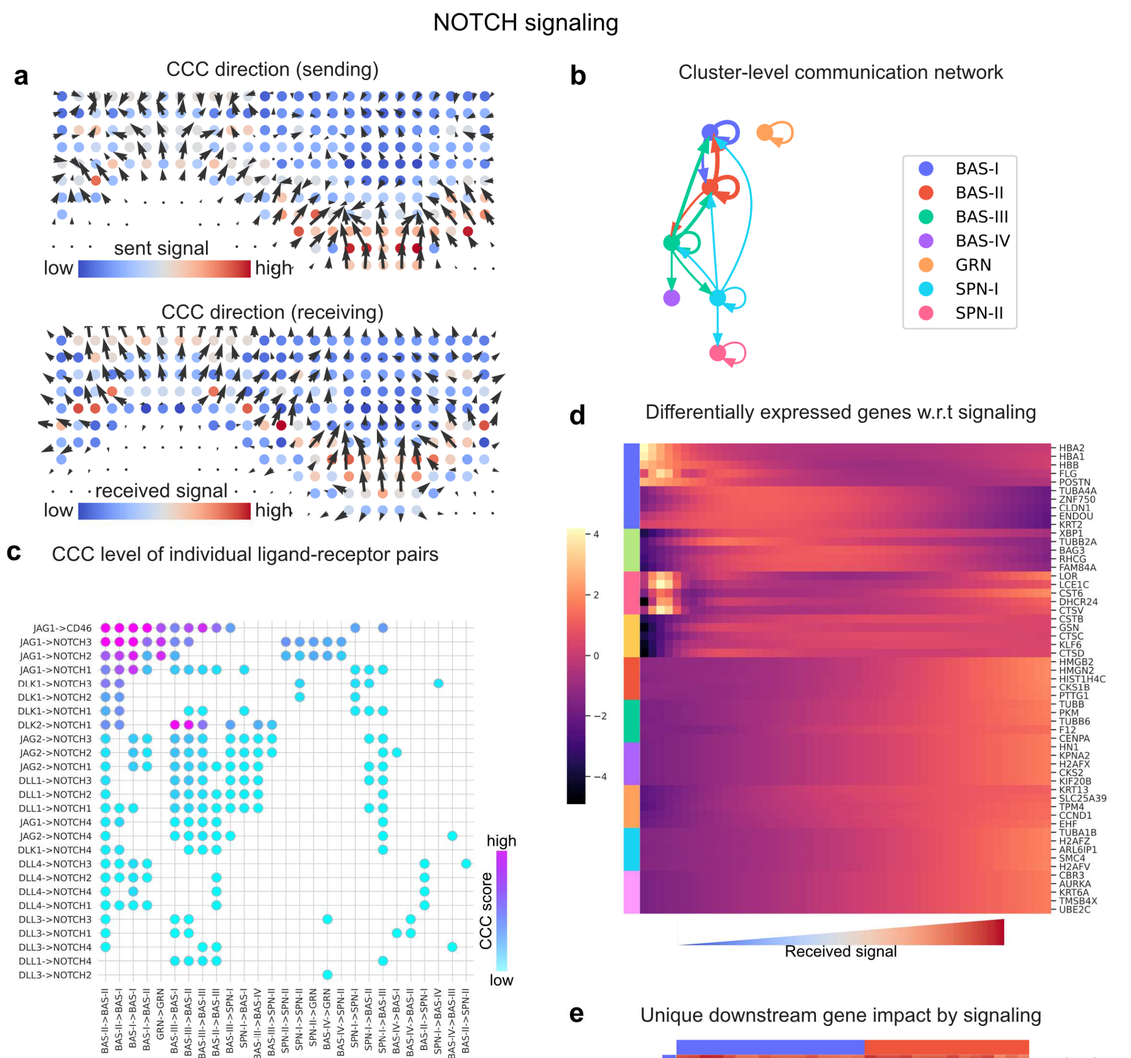

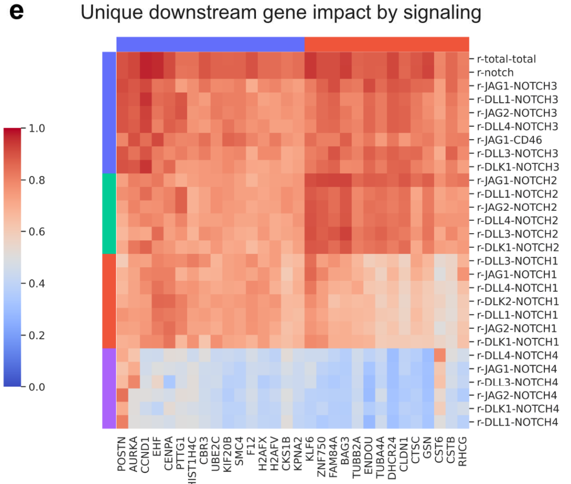

#### Supplementary Figure 16

#### NOTCH signaling in human skin

**a** Cell-cell communication directions. Sender: the direction shows to where the signal sending cells are sending the signal and color depicts the amount of signal sent by each cell. Receiver: the direction shows from where the signaling receiving cells are receiving the signal and color depicts the amount of signal received by each cell. **b** The cell-cell communication of all ligand-receptor pairs summarized to clusters (clustering identical to the recent NC paper), a thicker arrow means stronger signaling activity. **c** Dotplot showing the detailed signaling strength among the clusters through the individual ligand-receptor pairs. **d** The identified differentially expressed genes due to cell-cell communication activity. This is analogous to pseudotime DE genes but with the horizontal axis describing the amount of received signal. **e** Unique impact of CCC on downstream genes. A high score means the ligand-receptor pair is likely affecting the expression of the downstream gene considering the effects from other genes in the same cell.

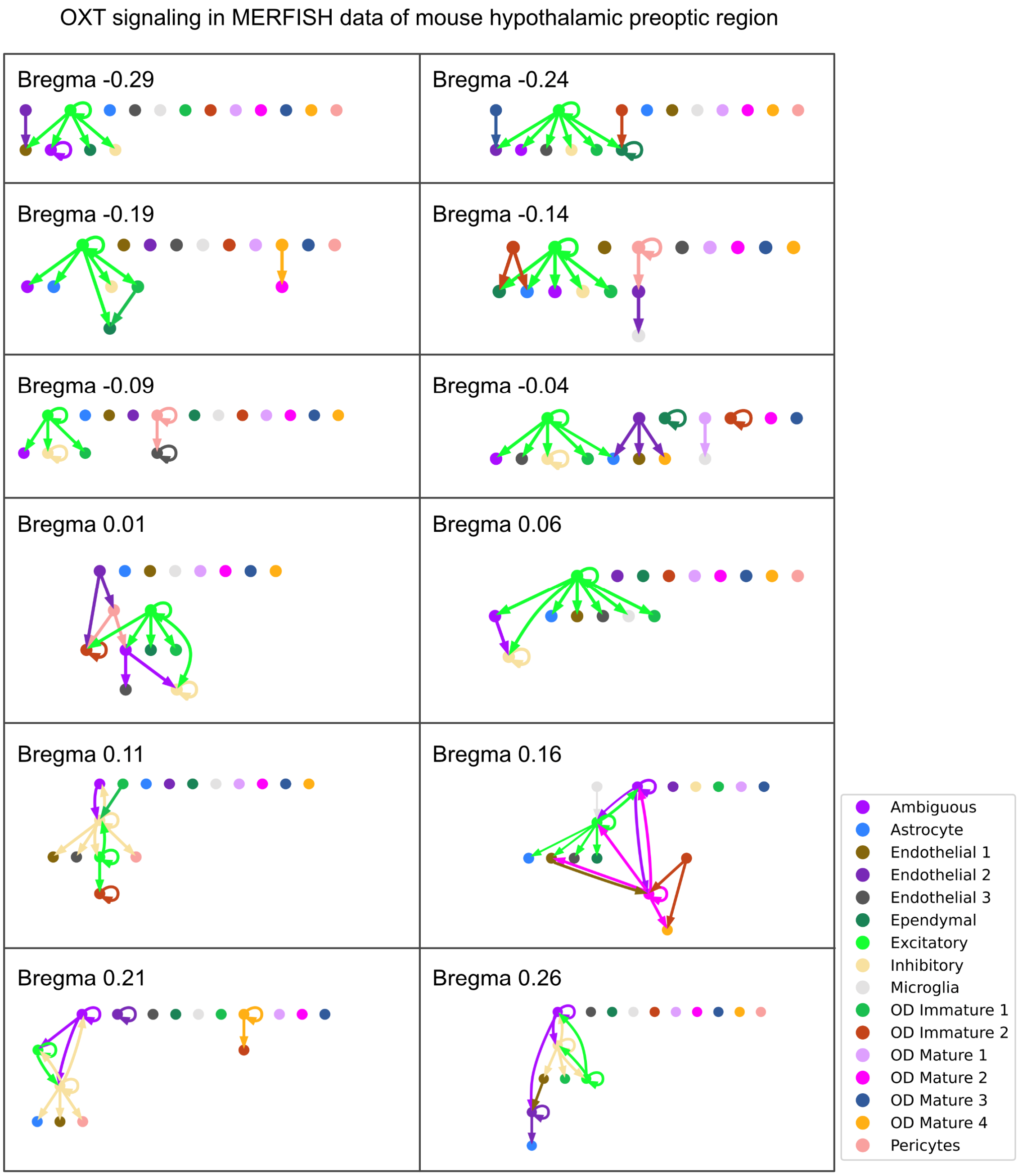

#### Supplementary Figure 17

Cluster-level OXT CCC in MERFISH mouse hypothalamic preoptic region

The inferred OXT CCC activity summarized to clusters in each of the slice of the MERFISH data.

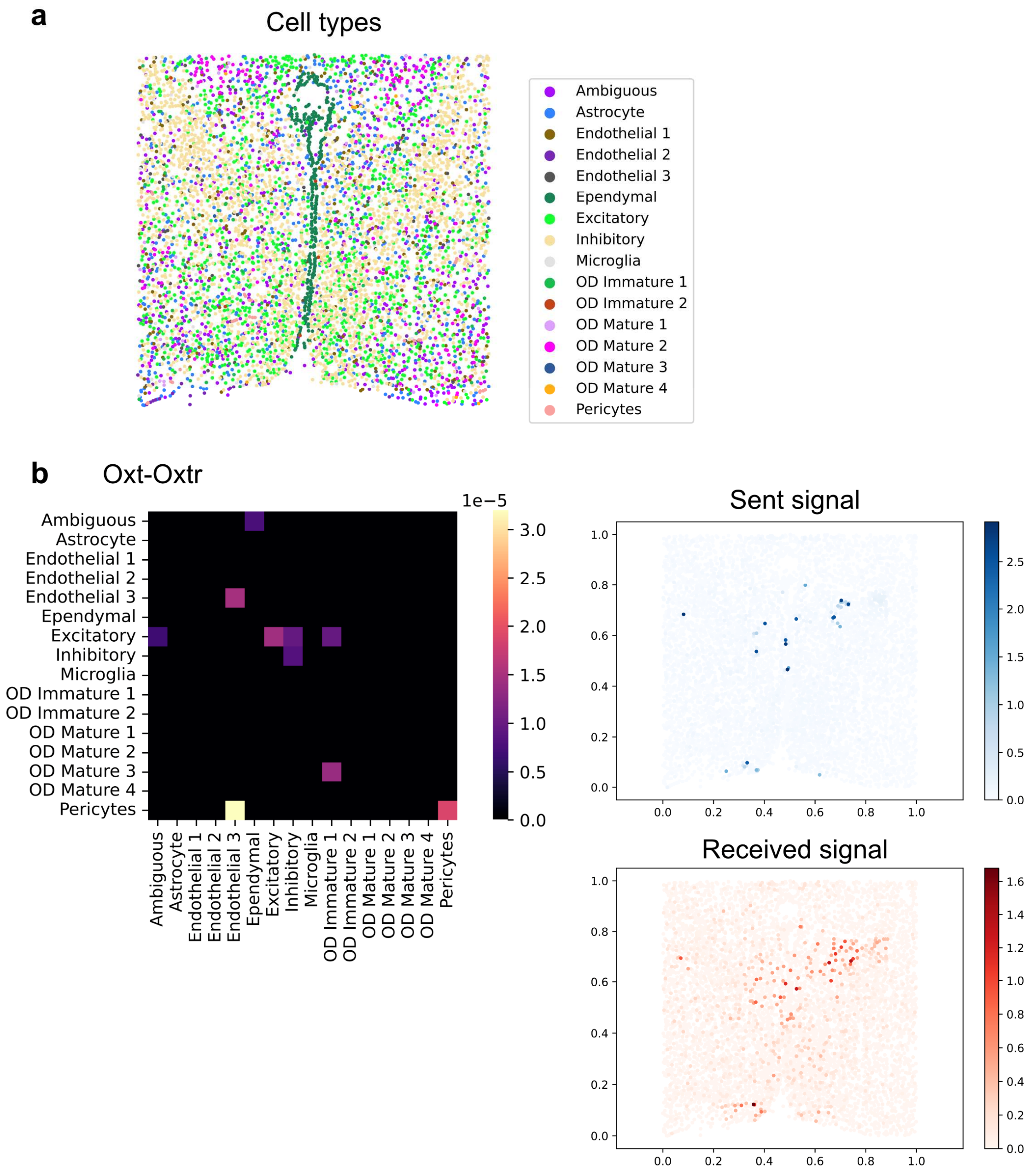

#### Supplementary Figure 18

OXT CCC in MERFISH mouse hypothalamic preoptic region

**a** Cell type plot of MERFISH mouse hypothalamic preoptic region data. **b** Heat map of cluster-level CCC where rows and columns correspond to senders and receivers, and the scatter plot of inferred sent and received signal through the ligand-receptor pair Oxt-Oxtr.

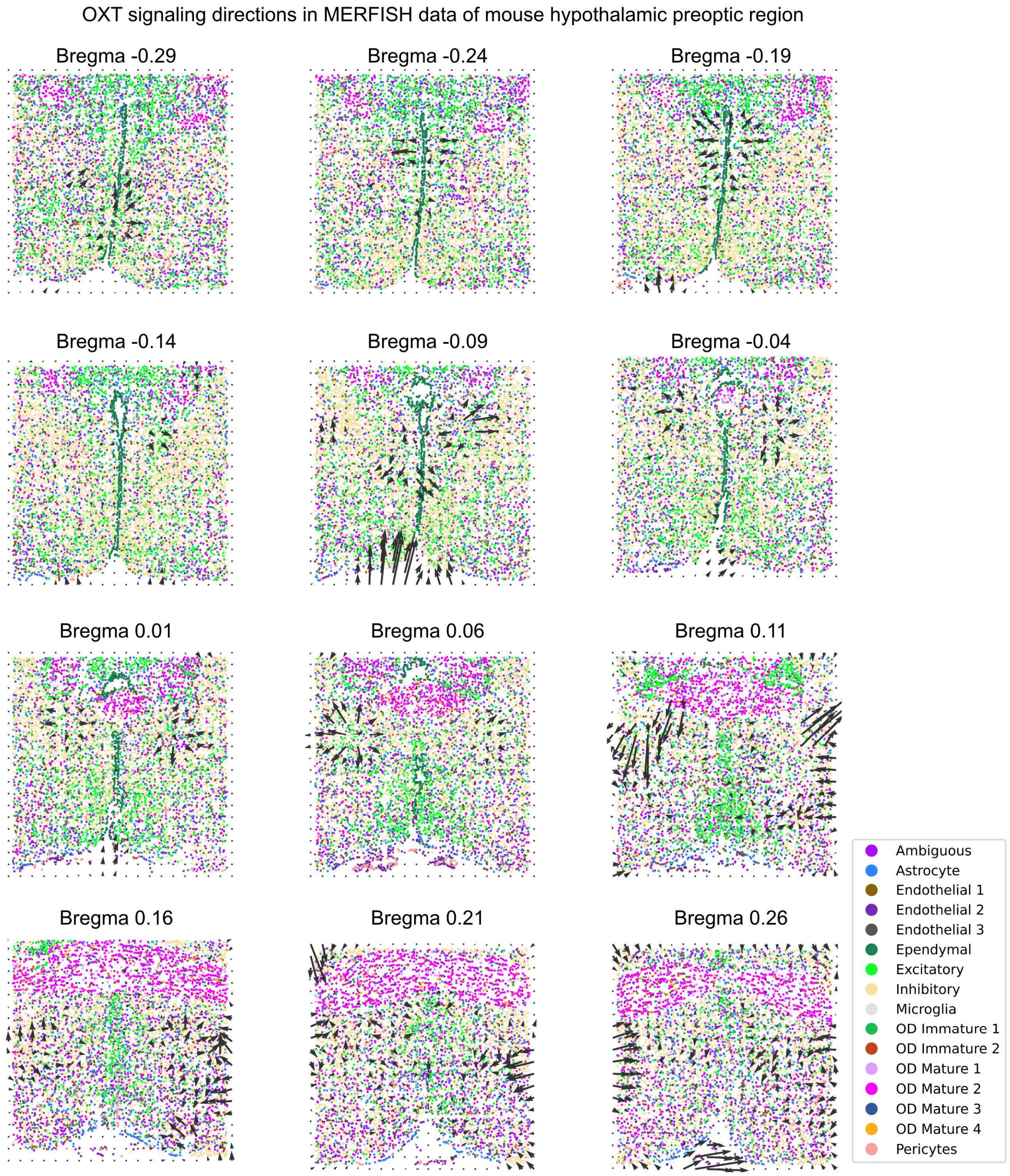

#### Supplementary Figure 19

OXT signaling directions in MERFISH mouse hypothalamic preoptic region

The inferred OXT signaling directions in each of the slice of the MERFISH data.

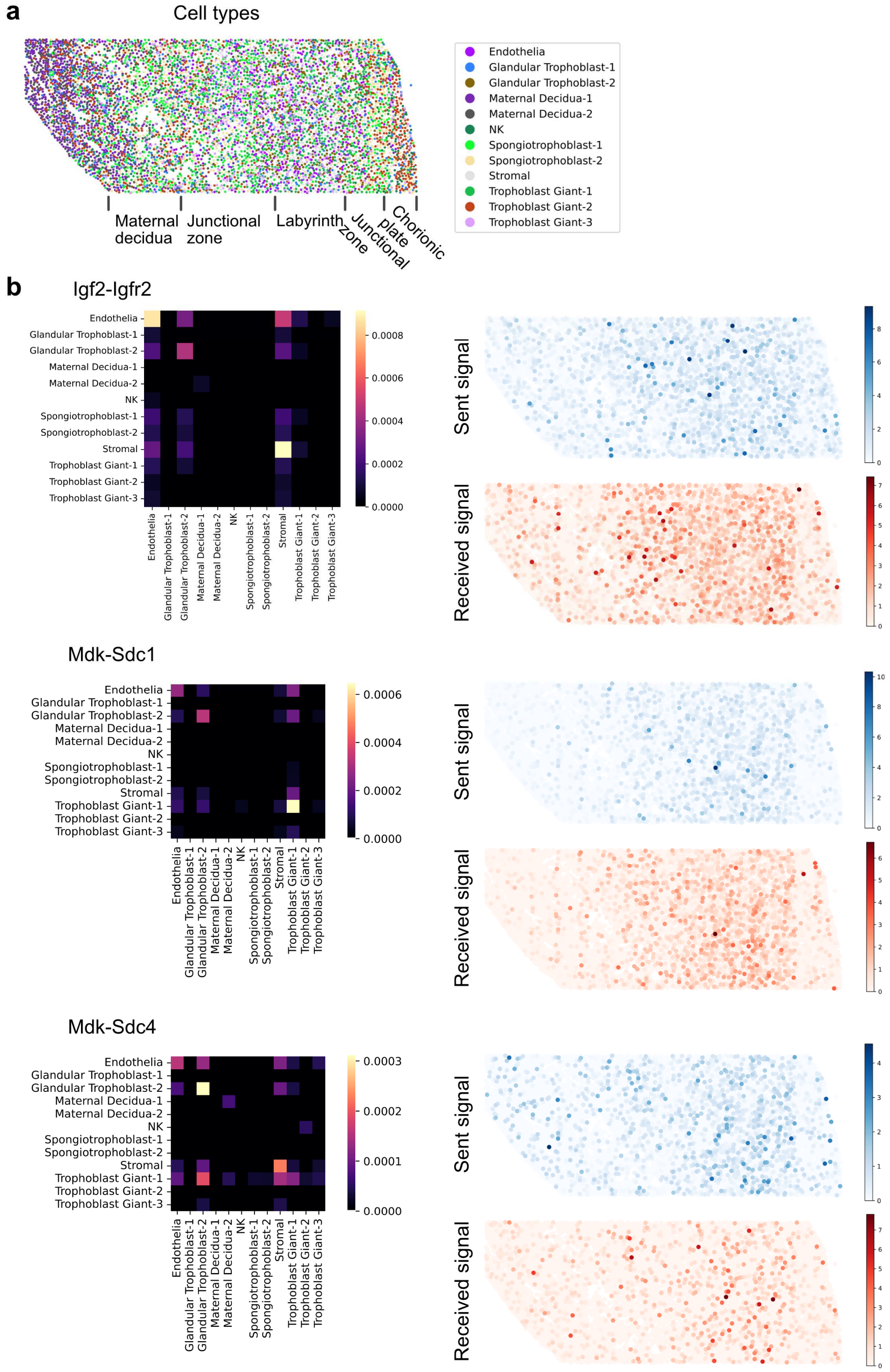

### Supplementary Figure 20

#### CCC in STARmap mouse placenta data

a Cell type plot of the STARmap mouse placenta data. **b** Cluster-level CCC heatmaps where rows and columns correspond to signal senders and receivers respectively and the spatial distribution of sent and received signal through the ligand-receptor pairs Igf2-Igfr2, Mdk-Sdc1, and Mdk-Sdc4.

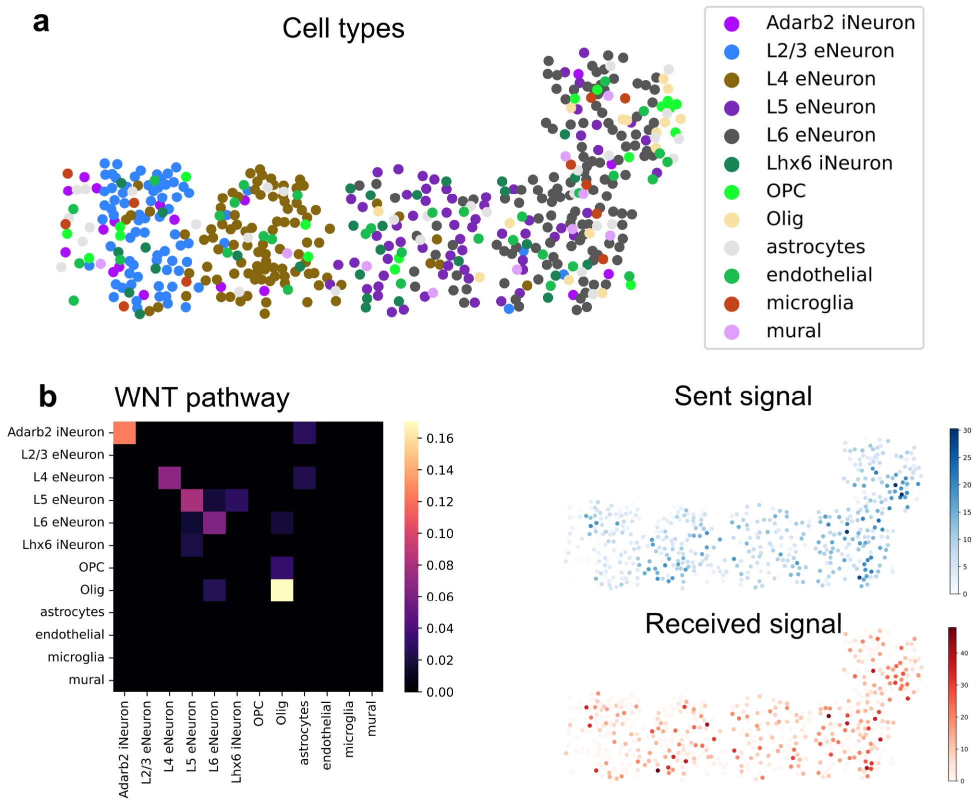

#### Supplementary Figure 21

WNT signaling in seqFISH+ mouse secondary somatosensory cortex

a Cell type plot of the seqFISH+ sscortex data. **b** Cluster-level CCC heatmaps where rows and columns correspond to signal senders and receivers respectively and the spatial distribution of sent and received signal through the ligand-receptor pairs from the WNT signaling pathway.

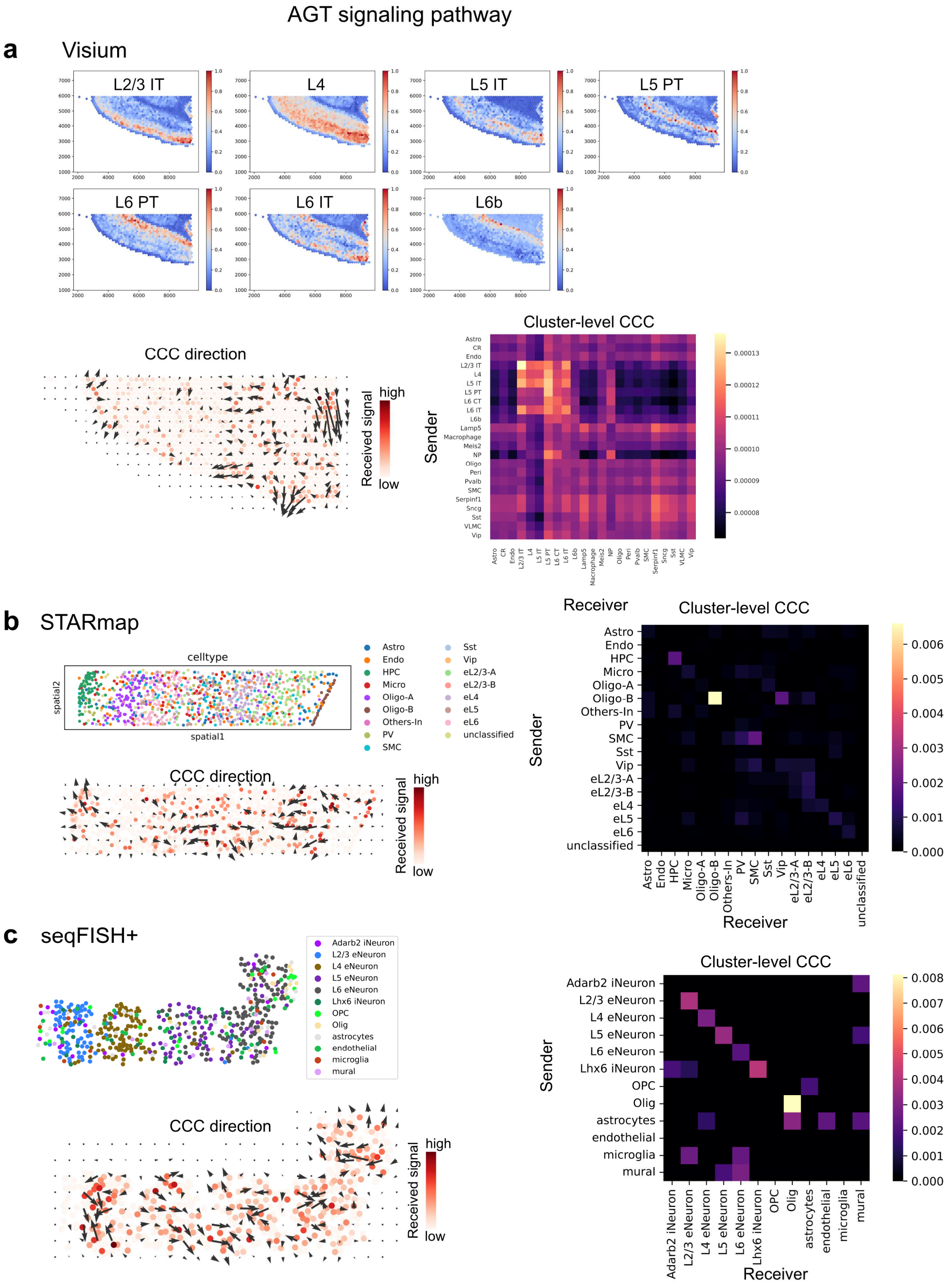

### Supplementary Figure 22

#### AGT signaling pathway in mouse cortex

The 1) cell type plots, 2) spatial directions of CCC, and 3) heatmaps of cluster-level CCC of the AGT signaling pathway in **a** Visium, **b** STARmap, and **c** seqFISH+ mouse cortex data.

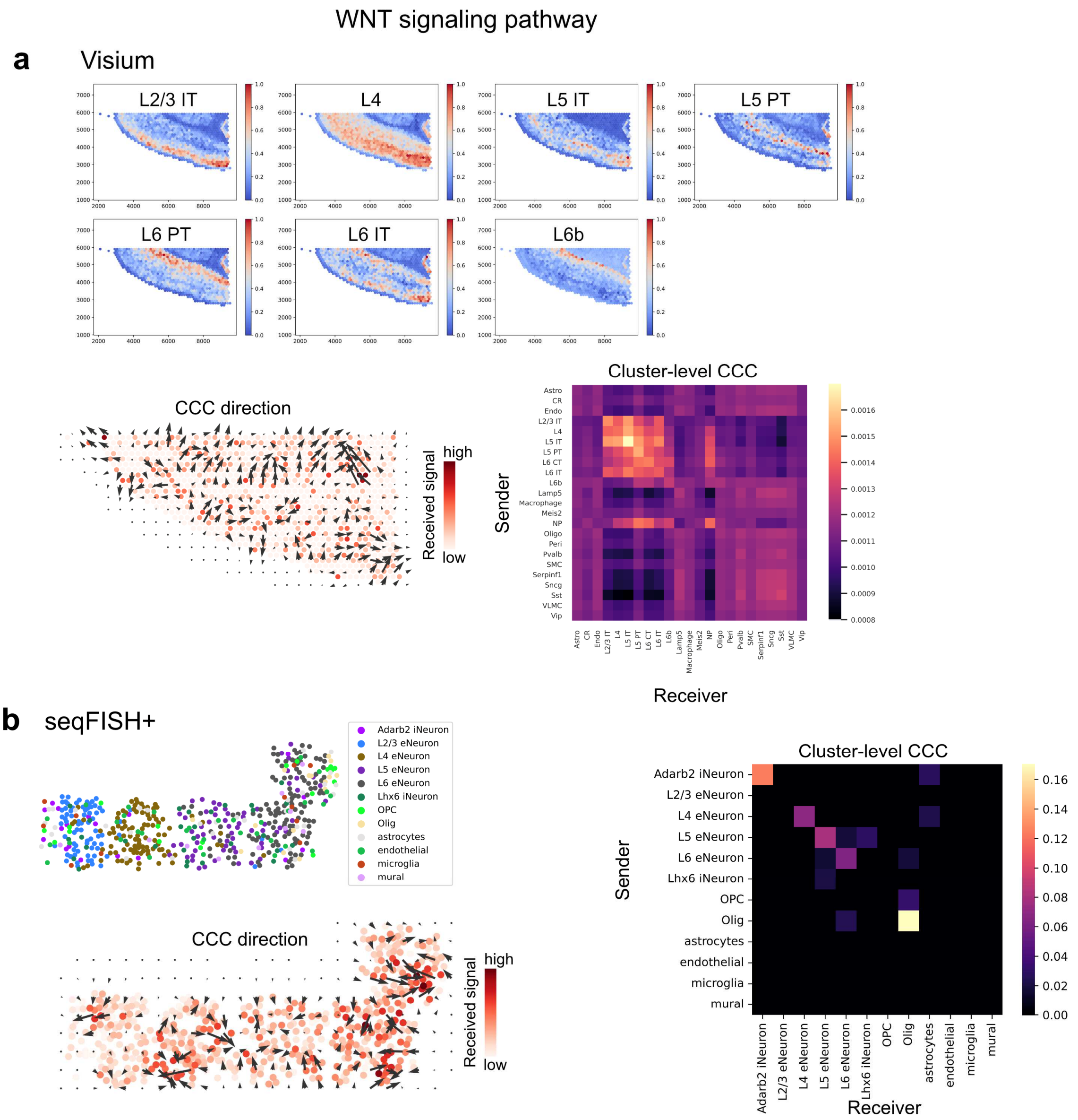

#### Supplementary Figure 23

WNT signaling pathway in mouse cortex

The 1) cell type plots, 2) spatial directions of CCC, and 3) heatmaps of cluster-level CCC of the WNT signaling pathway in **a** Visium, and **b** seqFISH+ mouse cortex data.

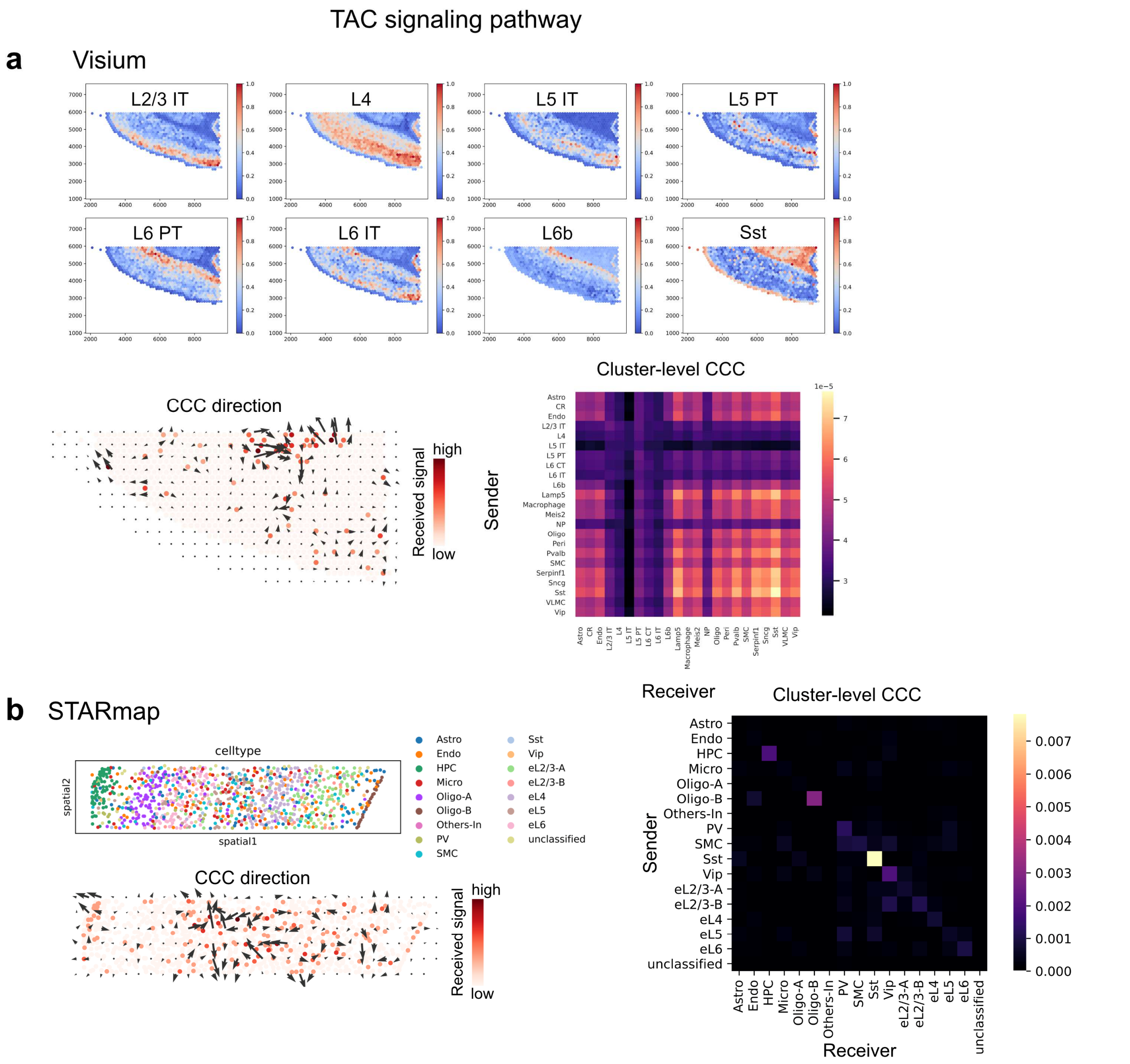

#### Supplementary Figure 24

TAC signaling pathway in mouse cortex

The 1) cell type plots, 2) spatial directions of CCC, and 3) heatmaps of cluster-level CCC of the TAC signaling pathway in **a** Visium, **b** STARmap mouse cortex data.

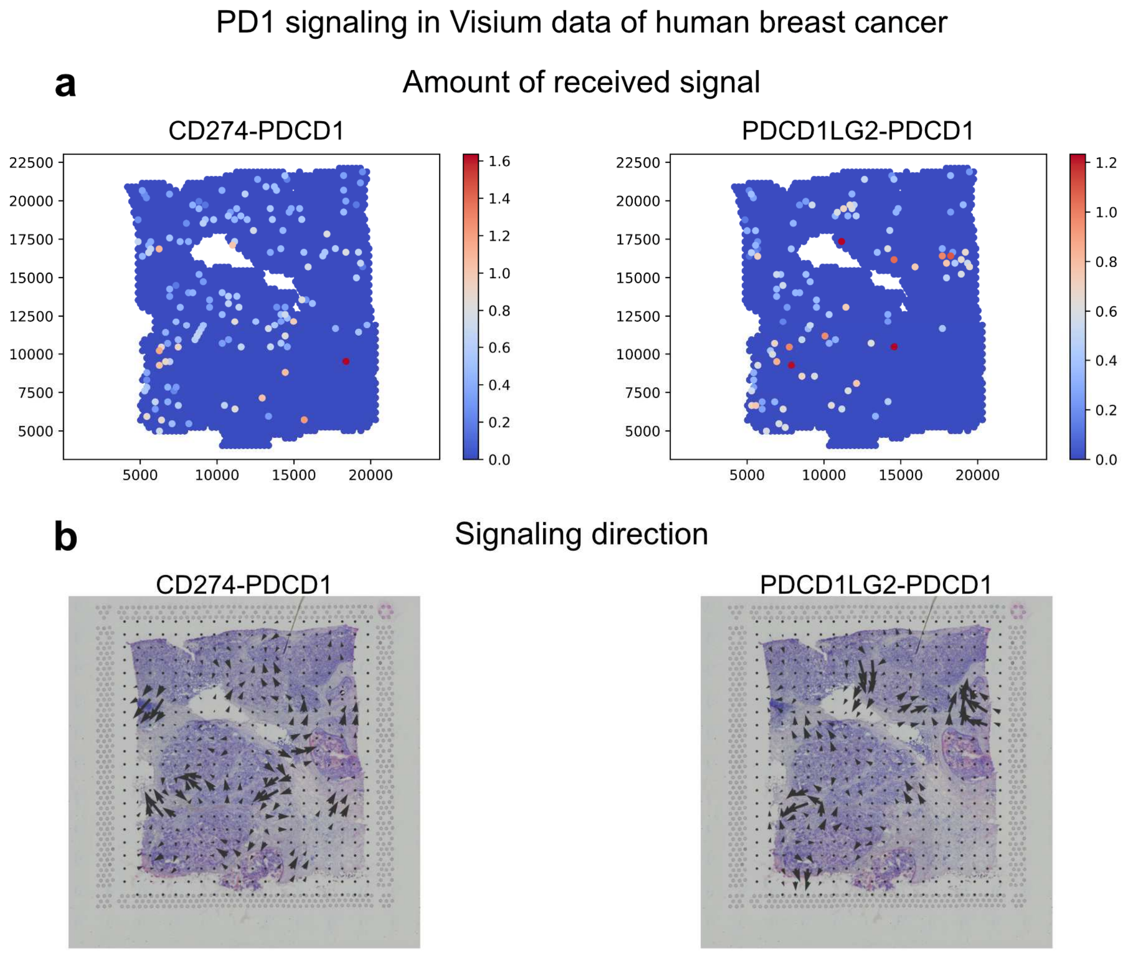

#### Supplementary Figure 25

PD1 signaling in the Visium data of human breast cancer.

**a** The amount of received signaling through two ligand-receptor pairs of PD1 signaling. **b** The signaling directions of the two ligand-receptor pairs.

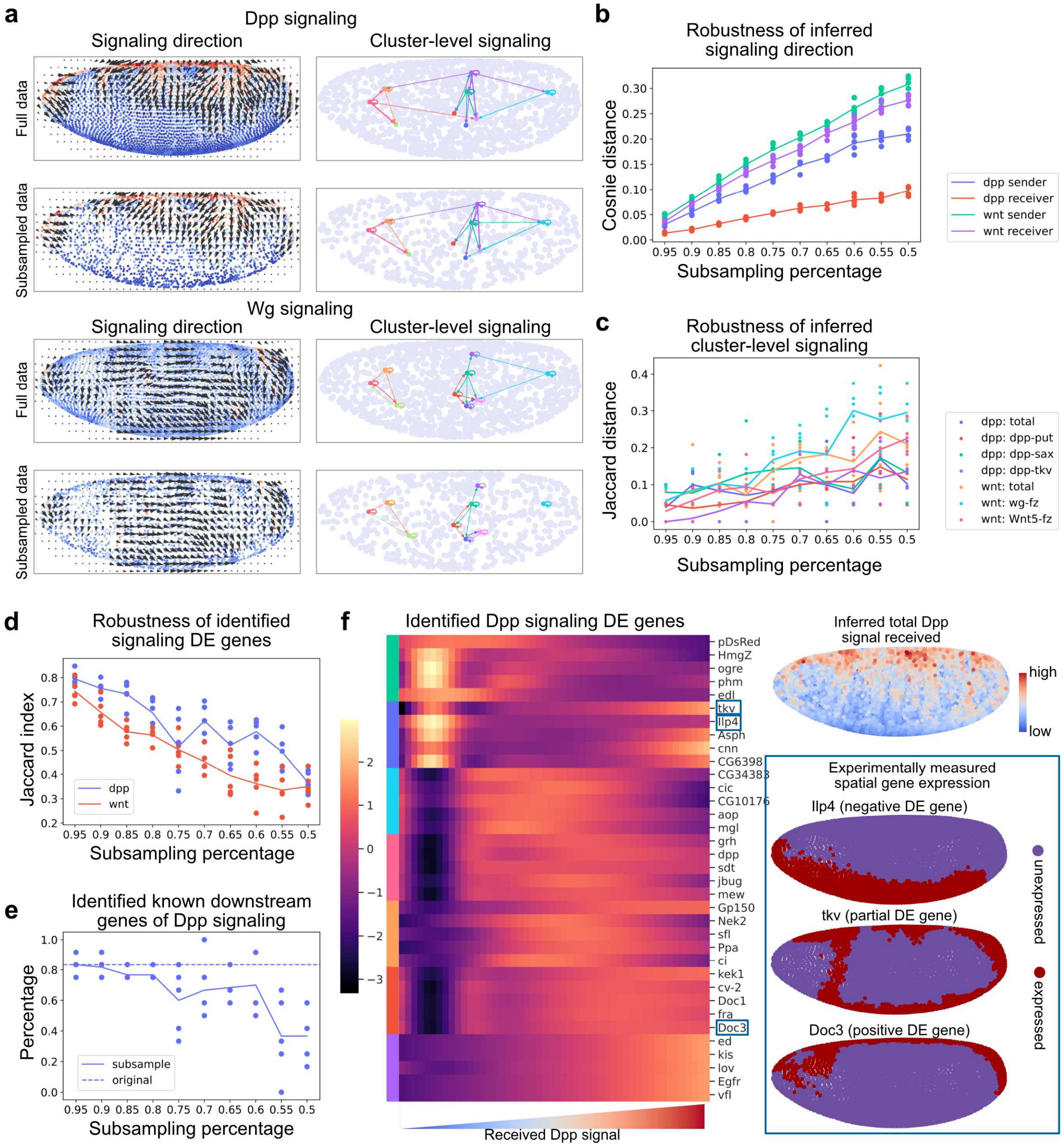

#### Supplementary Figure 26

Robustness of CCC analysis on a well-studied dataset

**a** The spatial signaling direction and signaling among cell clusters for Dpp and Wg signaling pathways. **b** Robustness of inferred signaling direction evaluated by comparing the direction obtained from subsampled dataset to the one from the full dataset using cosine distance. Each point is an independent test and the line shows the average of the tests. **c** Robustness of inferred cluster-level communication evaluated by comparing random subsamples to the full dataset using the Jaccard distance. **d** Robustness of downstream gene identification. **e** Percentage of known downstream genes that are identified as differentially expressed gene due to signaling activity. **f** Examples of the identified positively, negatively, and partially differentially expressed genes associated to Dpp signaling.

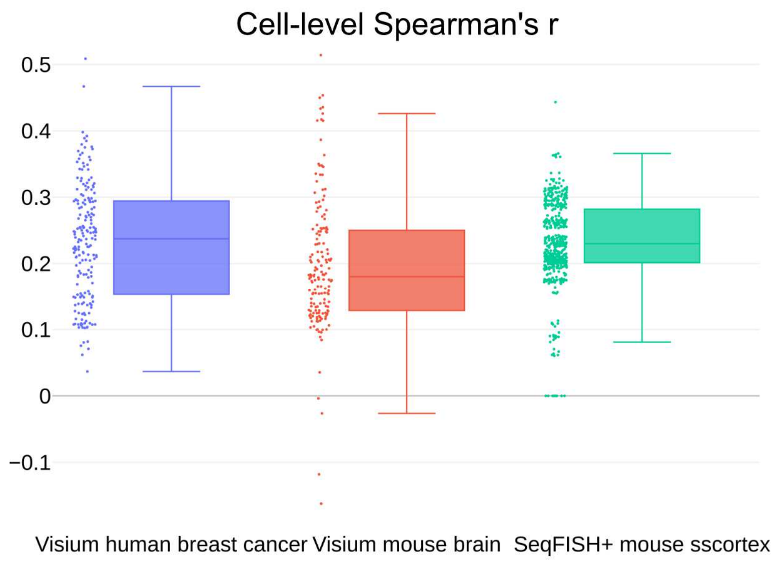

#### Supplementary Figure 27

Cell-level correlation between inferred signaling and expression of known downstream genes

Each dot represents a ligand-receptor pair. The Spearman’s correlation coefficient is computed between the amount of received signal and the average expression of known downstream genes.

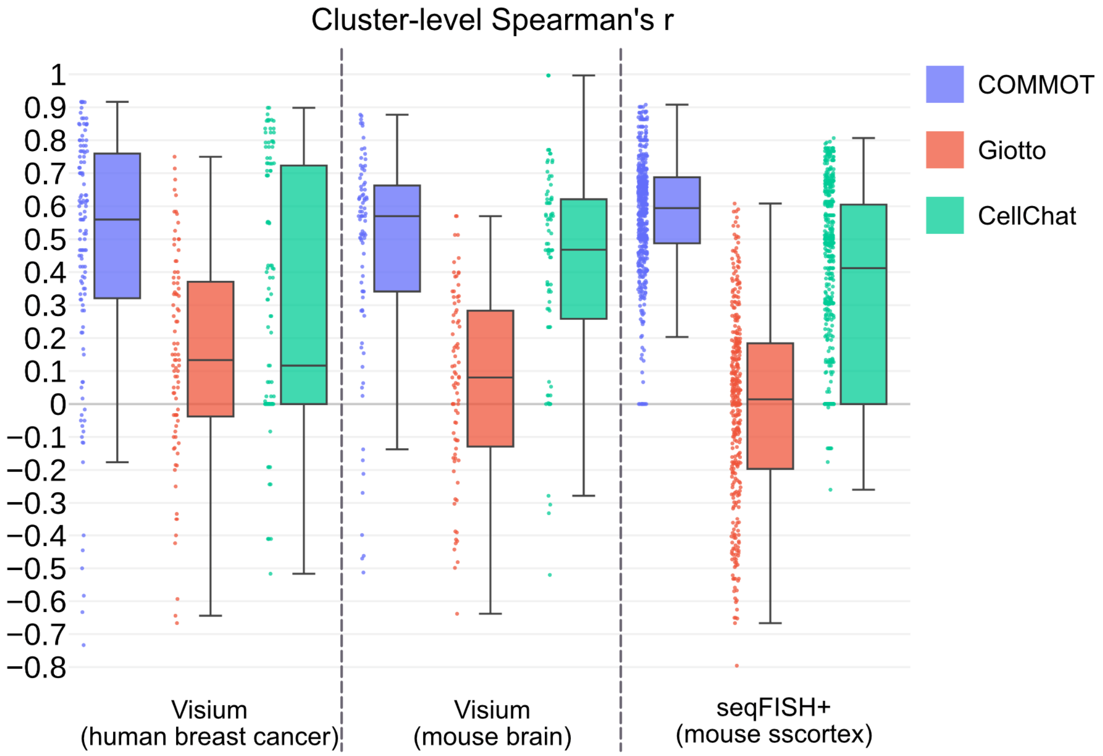

#### Supplementary Figure 28

##### Cluster-level correlation between inferred signaling and activity of known downstream genes (comparison with CellChat and Giotto)

Each dot represents a ligand-receptor pair. The activity of known downstream genes of a ligand-receptor pair is quantified for each cell cluster as the percentage of positive significant differentially expressed ones (p-value < 0.05) in that cluster. Spearman's correlation coefficients are computed between the amount of received signal and the downstream gene activity. For COMMOT, the amount of received signal is quantified by the average received signal of the cells in that cluster. For Giotto and CellChat which output a cluster-level signaling table, the maximum CCC score with each cluster as the receiver is used to quantify the amount of received signal.

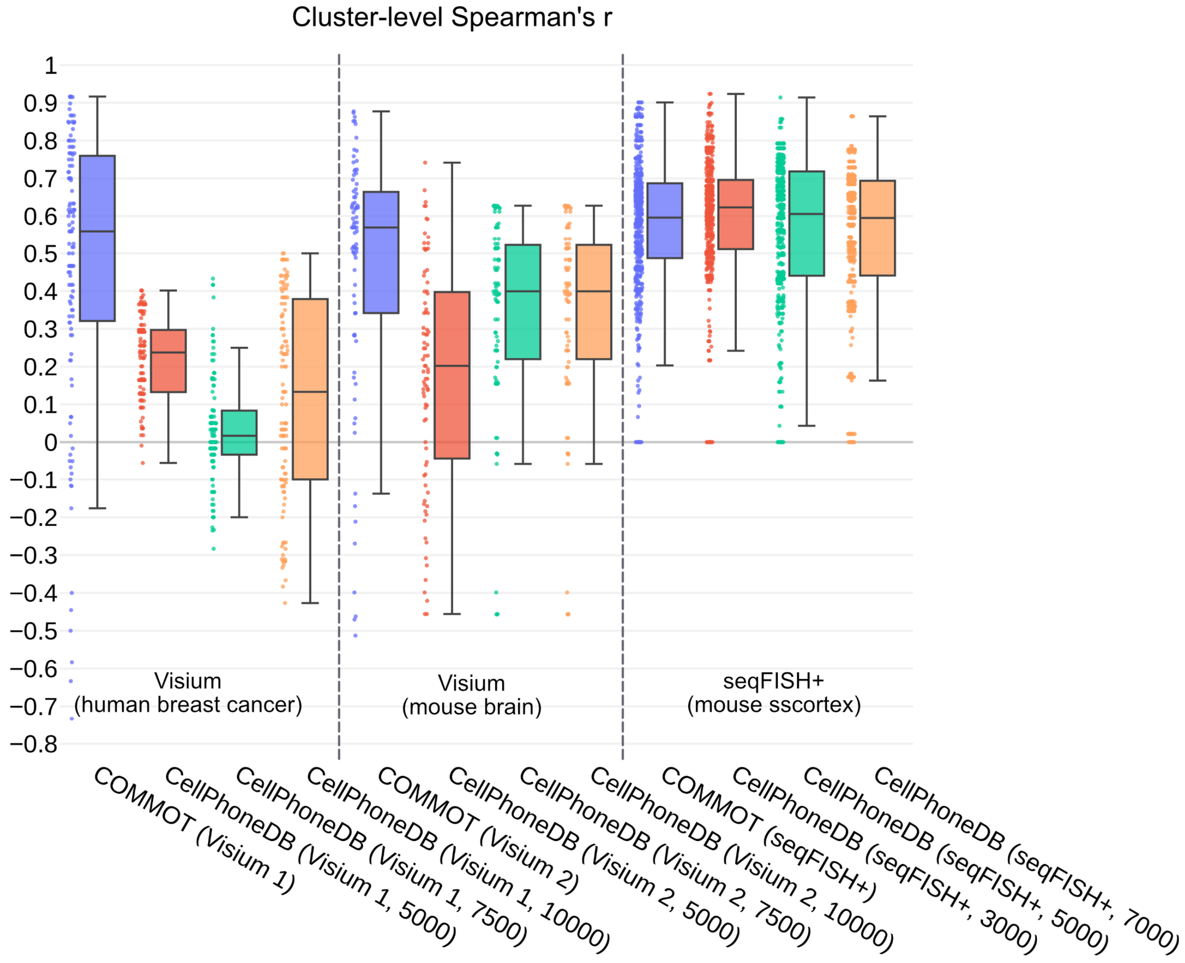

#### Supplementary Figure 29

Cluster-level correlation between inferred signaling and activity of known downstream genes (comparison with CellPhoneDB v3)

Each dot represents a ligand-receptor pair. The activity of known downstream genes of a ligand-receptor pair is quantified for each cell cluster as the percentage of positive significant differentially expressed ones (p-value < 0.05) in that cluster. Spearman's correlation coefficients are computed between the amount of received signal and the downstream gene activity. For COMMOT, the amount of received signal is quantified by the average received signal of the cells in that cluster. For CellPhoneDB, the total CCC score with each cluster as the receiver is used to quantify the amount of received signal. For COMMOT, a uniform 1000 μm (spot/cell level) distance cutoff is applied. For CellPhoneDB, the distance between clusters is first quantified as the average distance between spots/cells of the two clusters, and different cutoffs are tried (5000 μm, 7500 μm, and 10000 μm for Visium datasets and 3000 μm, 5000 μm, and 7000 μm for the seqFISH+ data).

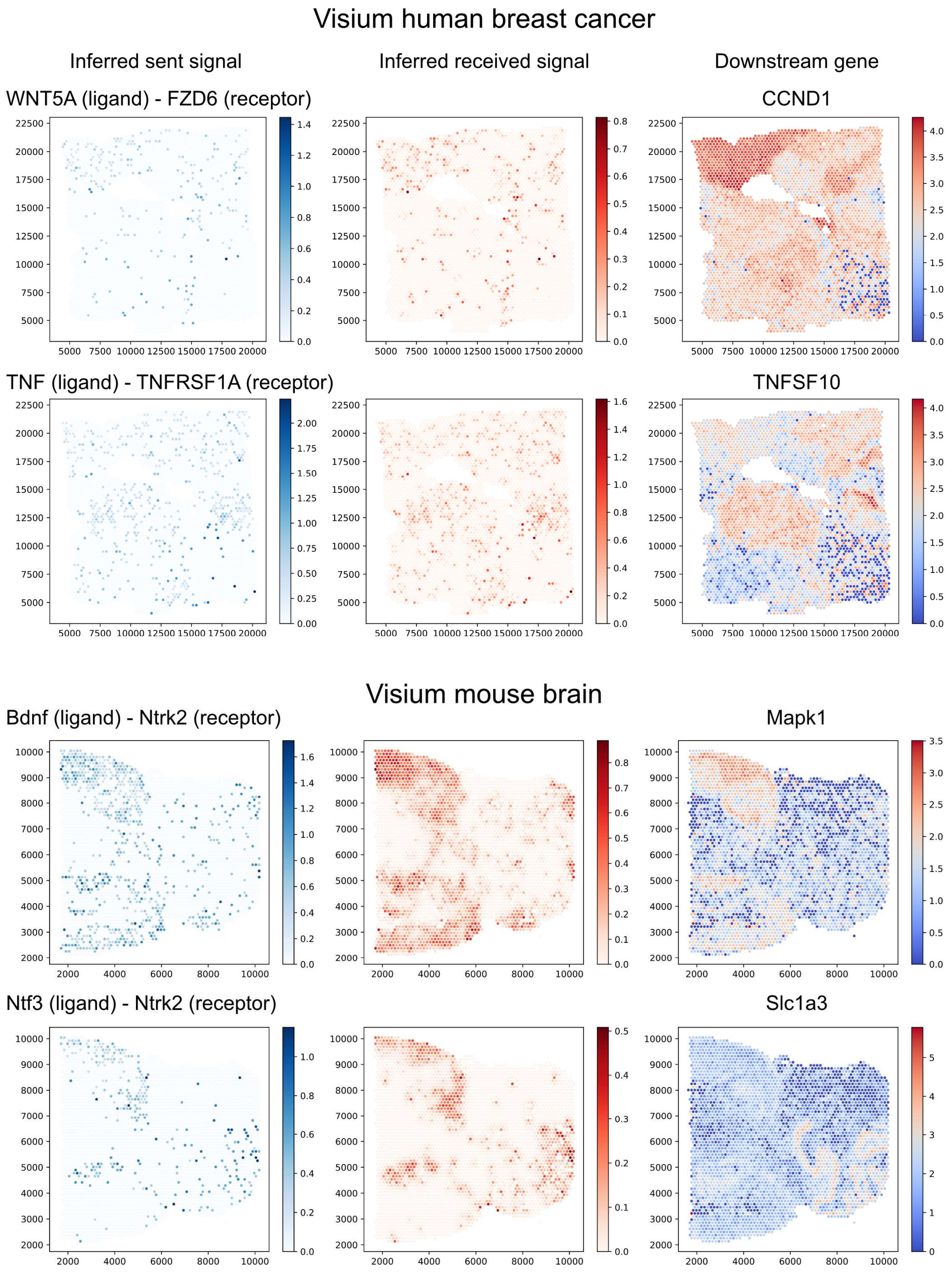

#### Supplementary Figure 30

Examples of inferred signaling and expression of known downstream genes.

The left column shows the ligands contributing to the inferred ligand-receptor complex. The middle column shows the inferred ligand-receptor complex. The right column shows the expression levels of known downstream target genes of the ligand-receptor pairs according to scSeqComm database.

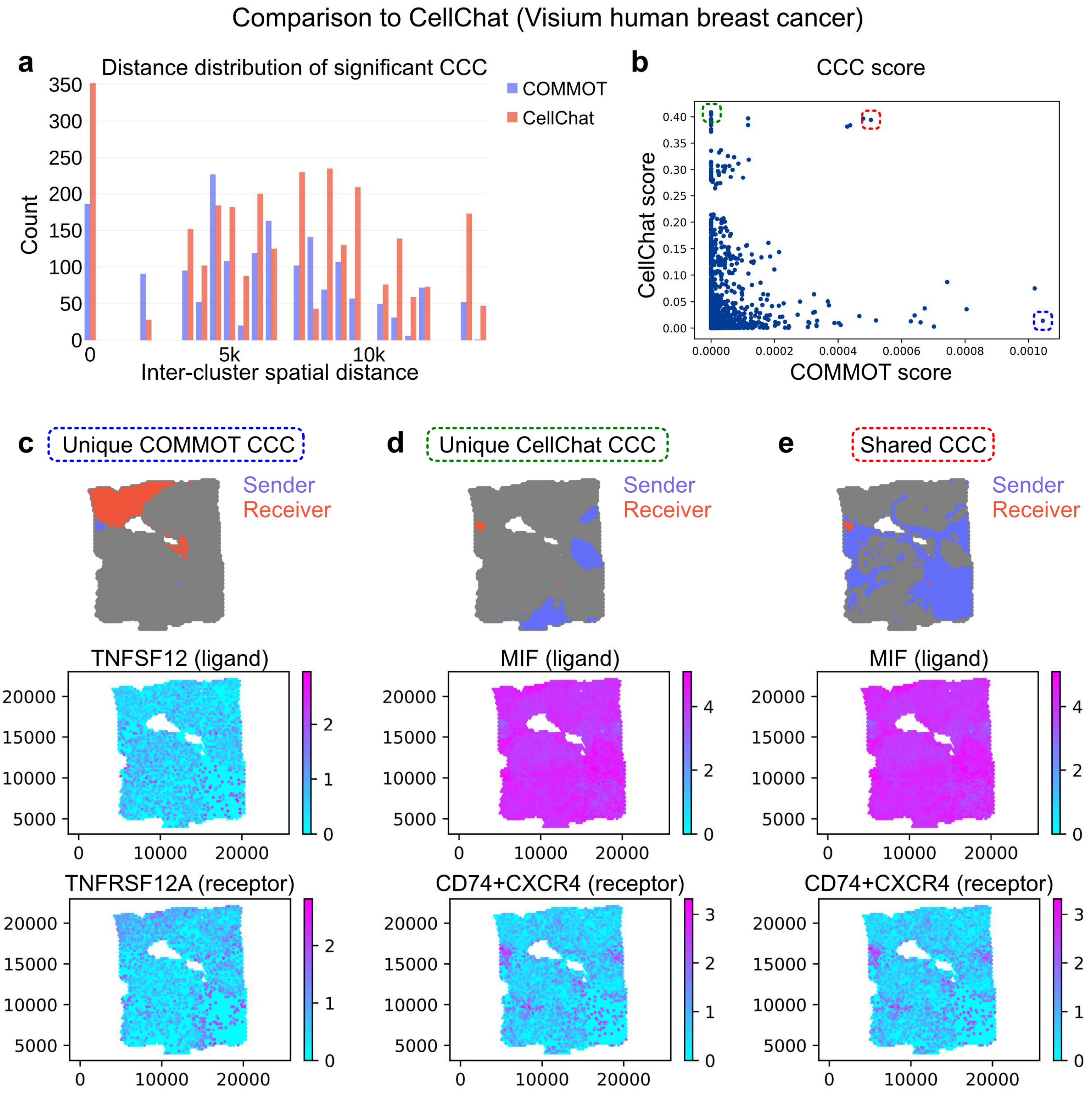

#### Supplementary Figure 31

Comparison with CellChat on a Visium human breast cancer dataset

**a** The distribution of inter-cluster distances (distance between geometric center of the clusters) of the significant CCC cluster pairs. **b** Each point represent a ligand-receptor pair and a cluster pair. **c-e** Example significant cluster-level CCC that are uniquely identified by COMMOT and CellChat, and by both methods.

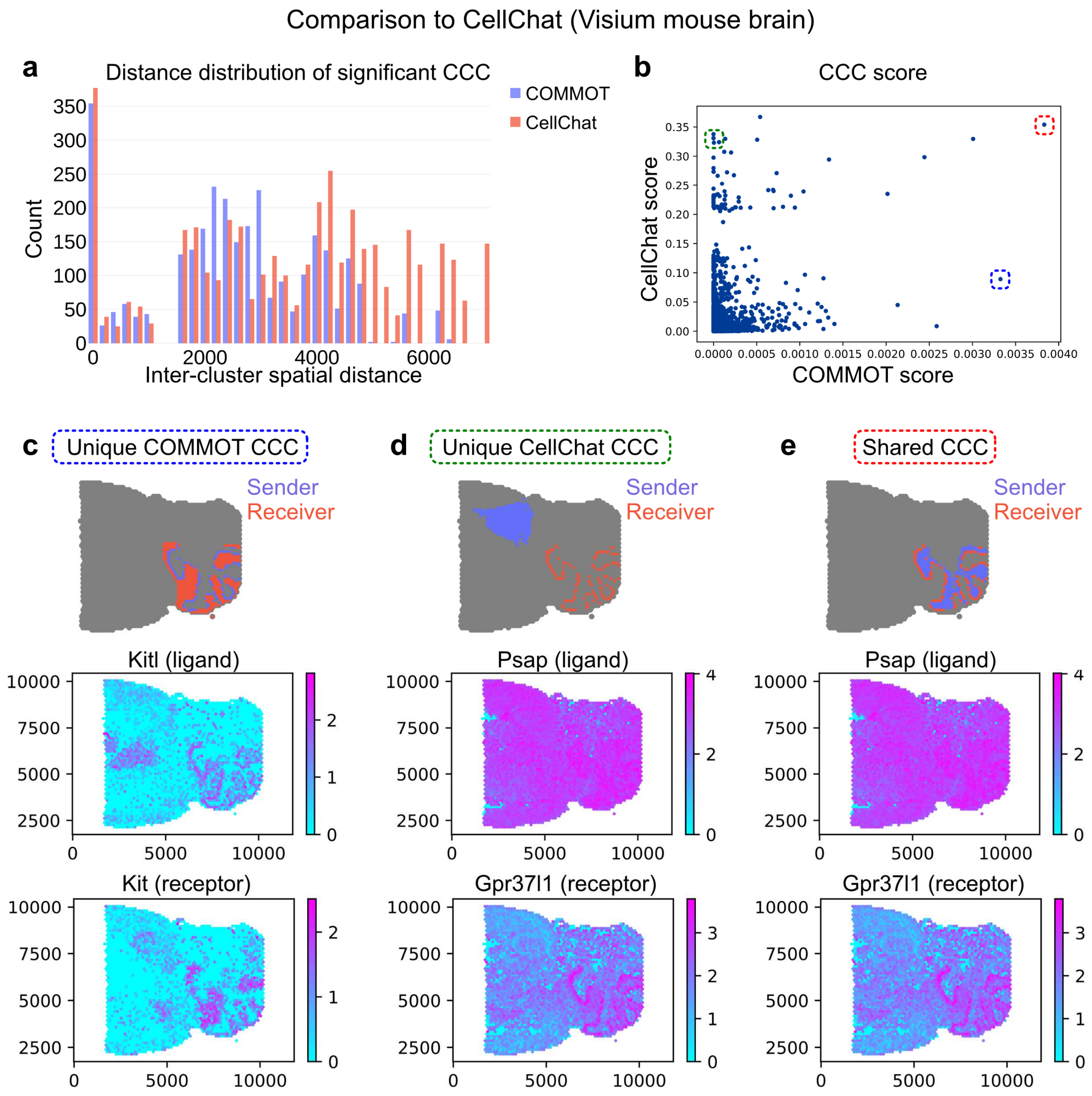

#### Supplementary Figure 32

Comparison with CellChat on a Visium mouse brain dataset

**a** The distribution of inter-cluster distances (distance between geometric center of the clusters) of the significant CCC cluster pairs. **b** Each point represent a ligand-receptor pair and a cluster pair. **c-e** Example significant cluster-level CCC that are uniquely identified by COMMOT and CellChat, and by both methods.

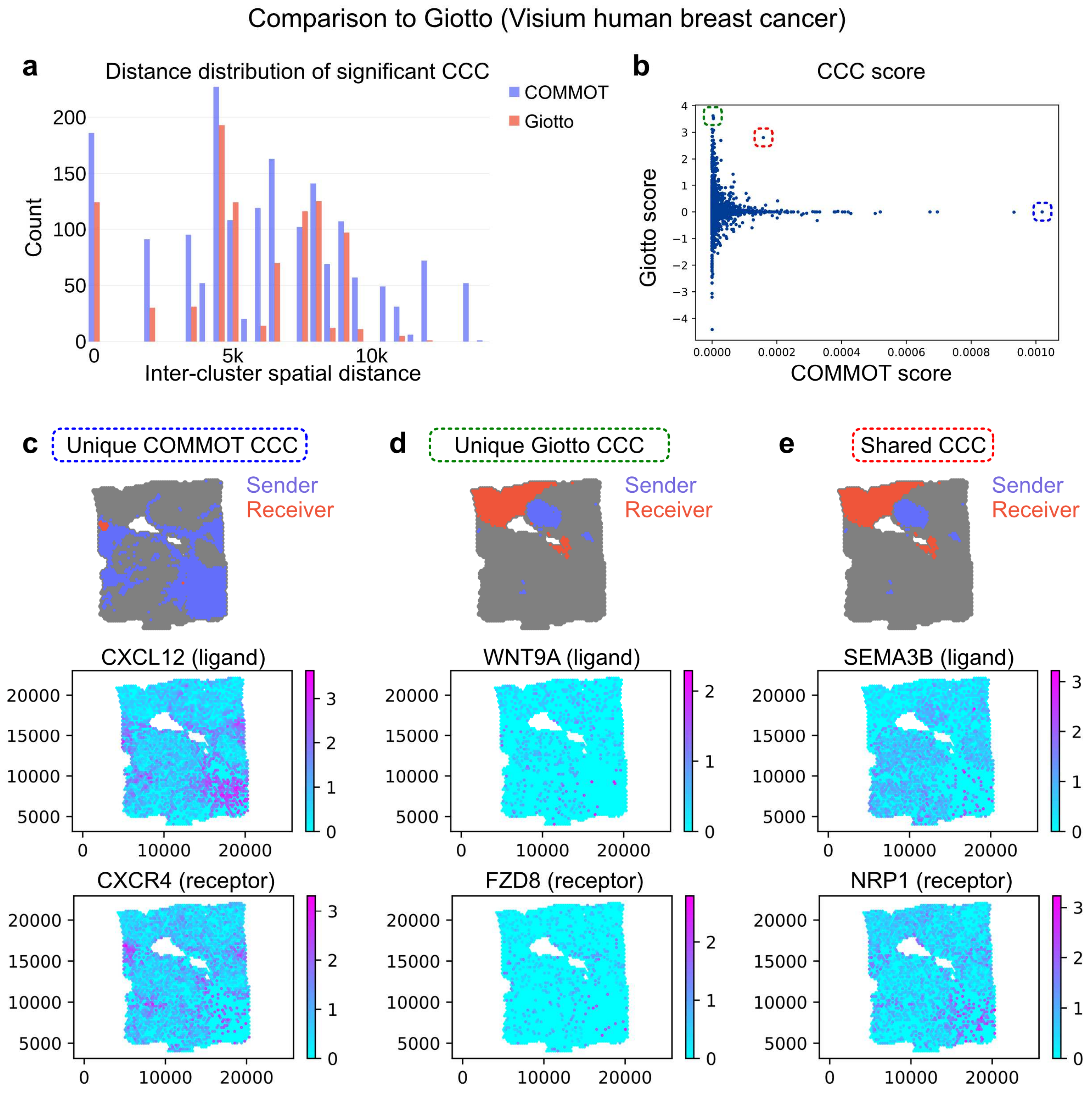

**Supplementary Figure 33**

Comparison with Giotto on a Visium human breast cancer dataset

**a** The distribution of inter-cluster distances (distance between geometric center of the clusters) of the significant CCC cluster pairs. **b** Each point represent a ligand-receptor pair and a cluster pair. **c-e** Example significant cluster-level CCC that are uniquely identified by COMMOT and Giotto, and by both methods.

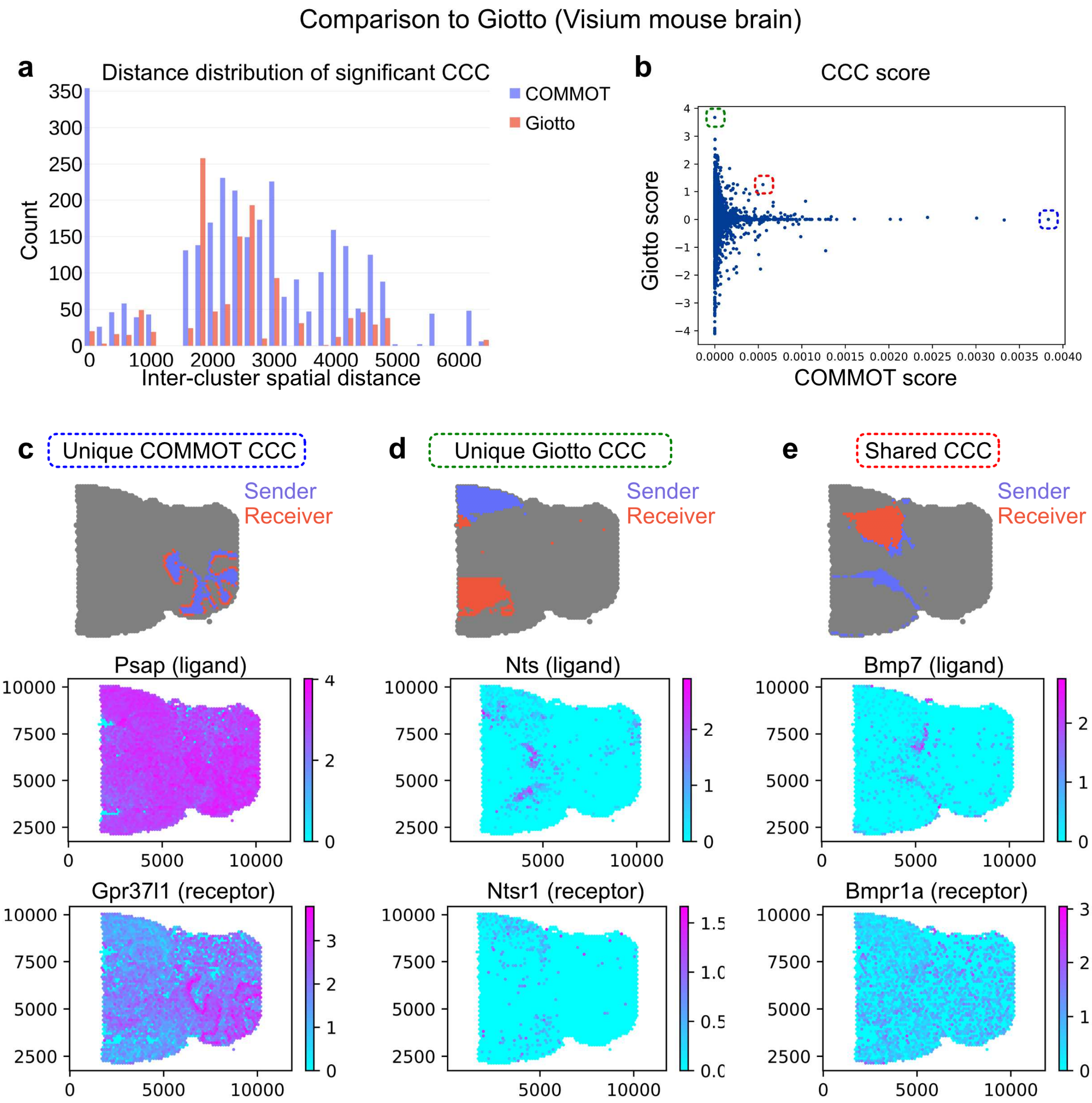

**Supplementary Figure 34**

Comparison with Giotto on a Visium mouse brain dataset

**a** The distribution of inter-cluster distances (distance between geometric center of the clusters) of the significant CCC cluster pairs. **b** Each point represent a ligand-receptor pair and a cluster pair. **c-e** Example significant cluster-level CCC that are uniquely identified by COMMOT and Giotto, and by both methods.

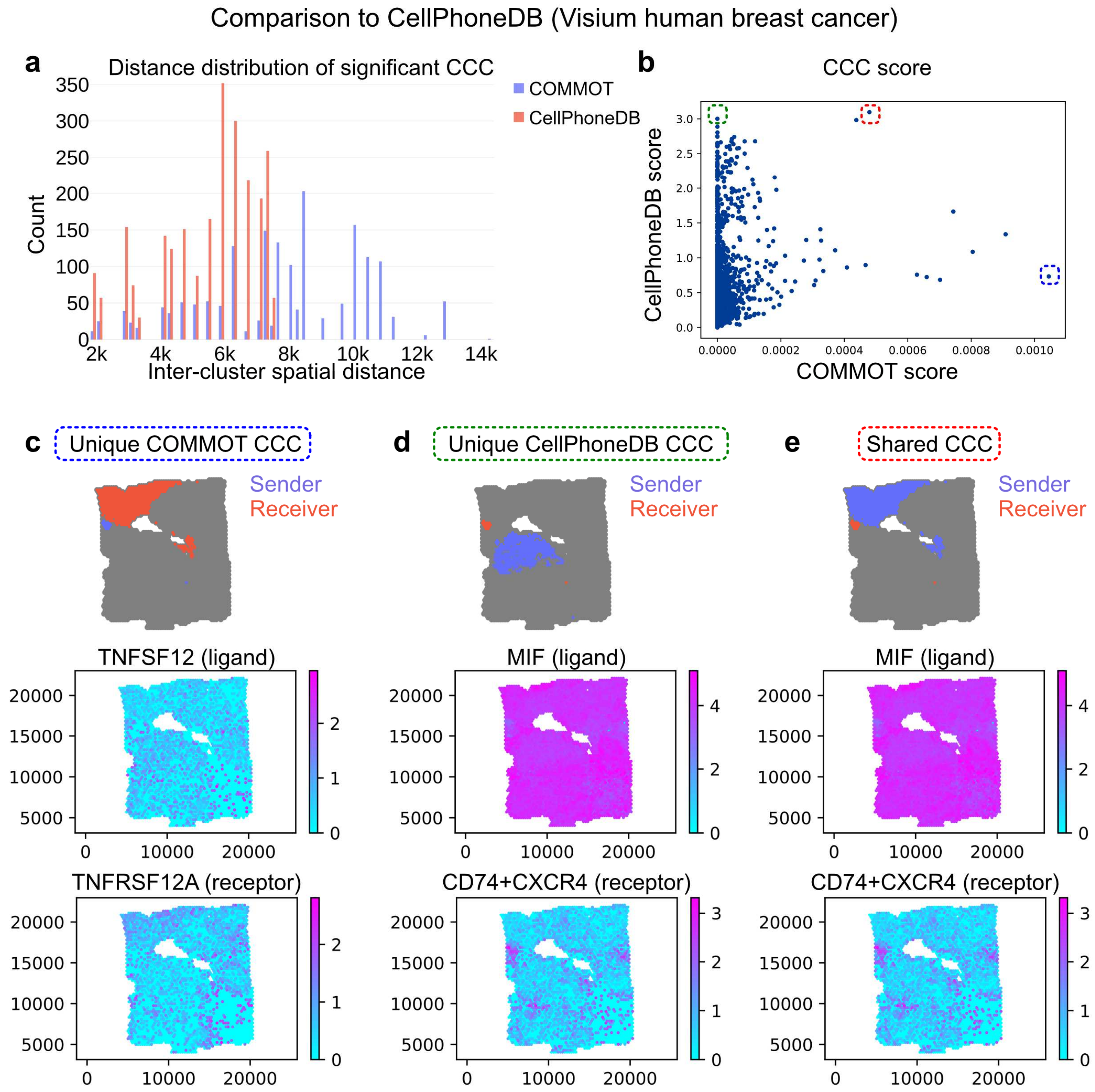

**Supplementary Figure 35**

Comparison with CellPhoneDB v3 on a Visium human breast cancer dataset

**a** The distribution of inter-cluster distances (distance between geometric center of the clusters) of the significant CCC cluster pairs. **b** Each point represent a ligand-receptor pair and a cluster pair. **c-e** Example significant cluster-level CCC that are uniquely identified by COMMOT and CellPhoneDB, and by both methods.

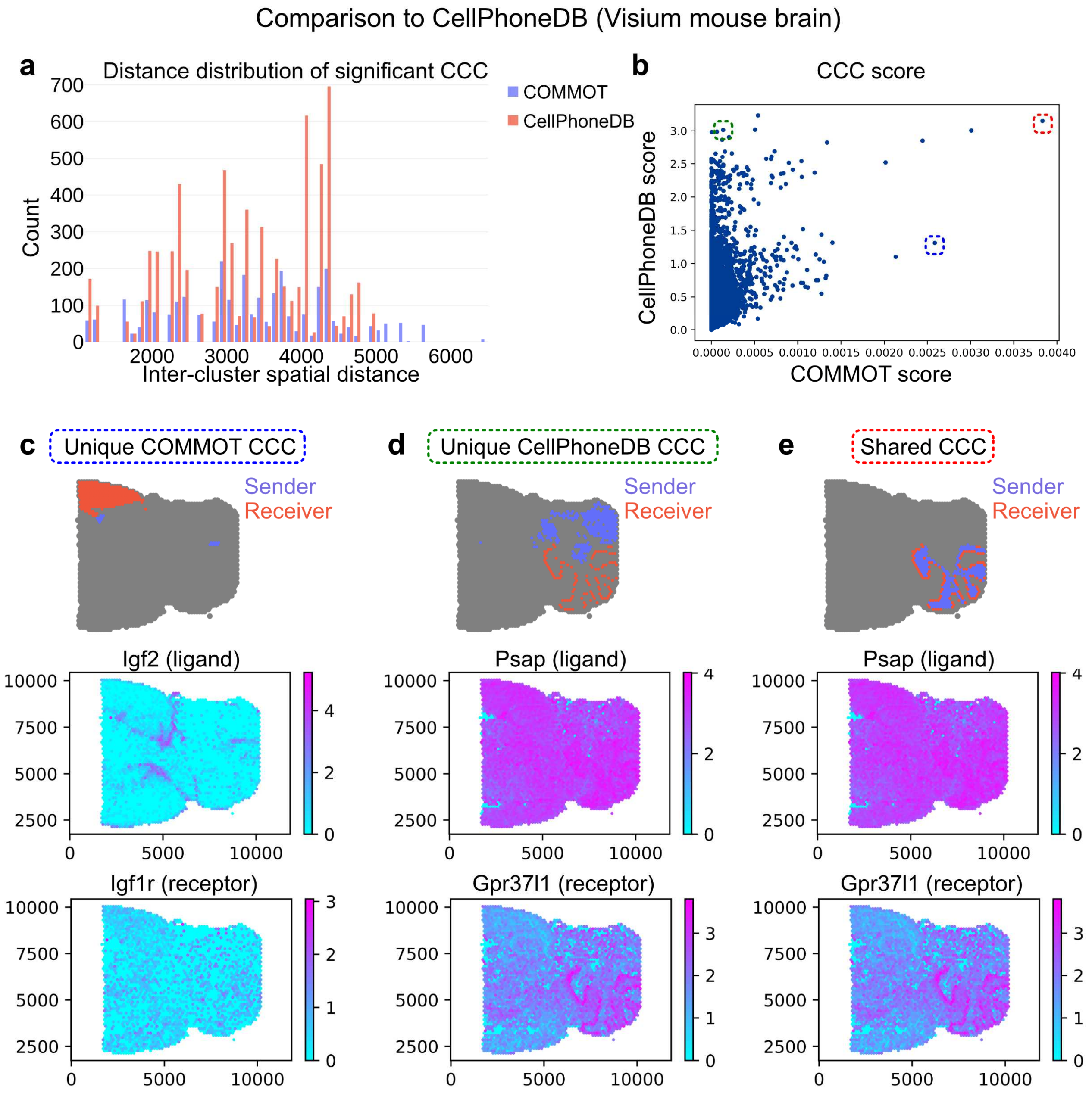

#### Supplementary Figure 36

Comparison with CellPhoneDB v3 on a Visium mouse brain dataset

**a** The distribution of inter-cluster distances (distance between geometric center of the clusters) of the significant CCC cluster pairs. **b** Each point represent a ligand-receptor pair and a cluster pair. **c-e** Example significant cluster-level CCC that are uniquely identified by COMMOT and CellPhoneDB, and by both methods.

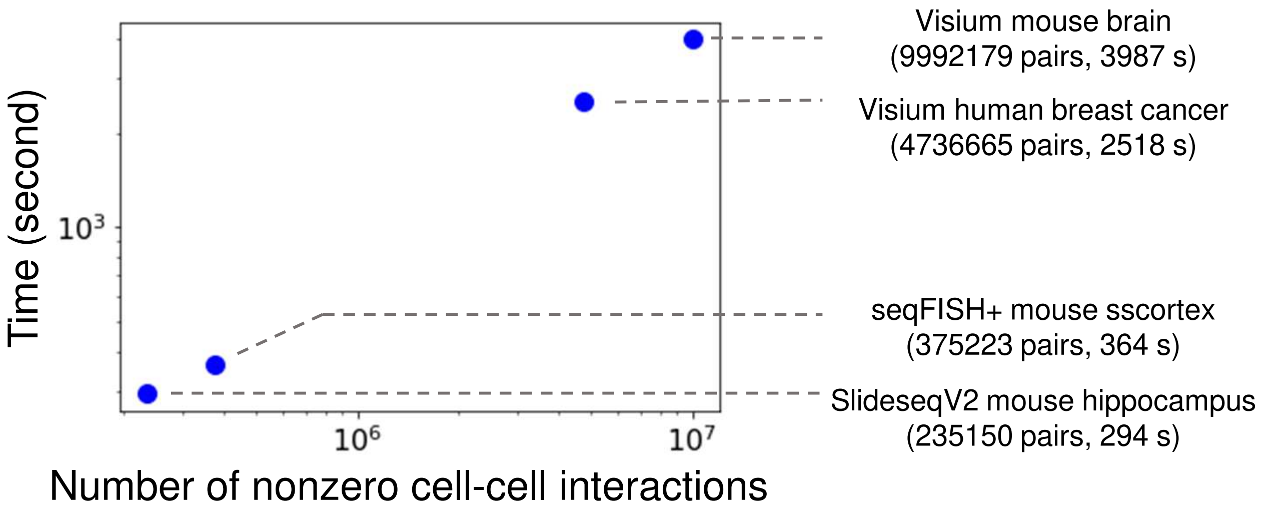

**Supplementary Figure 37**

CCC inference running time of COMMOT

The running time (tested on a desktop with i7-8700K running Ubuntu 20.04) of the collective optimal transport algorithm of COMMOT (the core CCC inference function) on several datasets. The wall time of computation is plotted against the number of nonzero elements in the interaction networks. Both axes are log transformed.

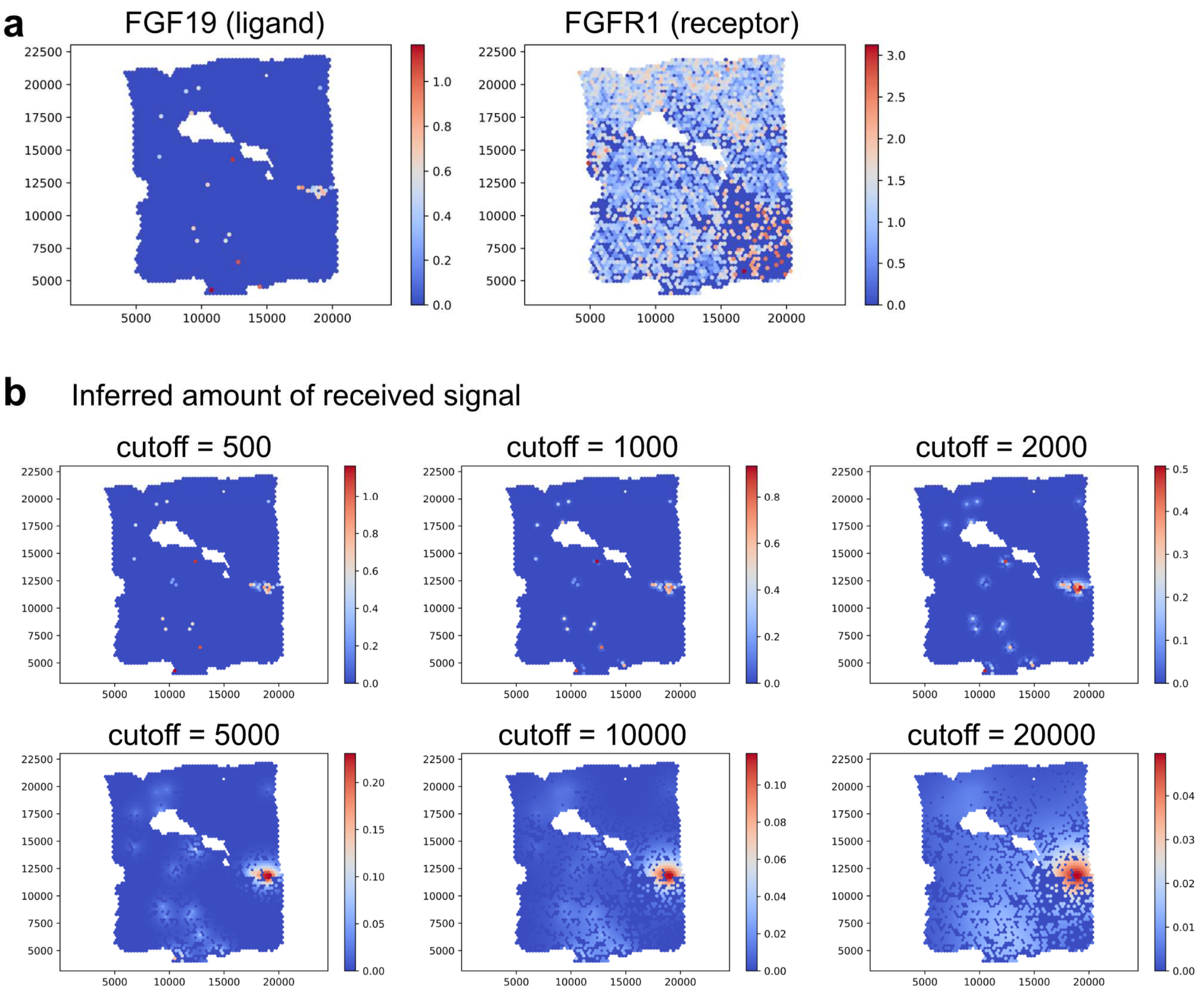

#### Supplementary Figure 38

Illustration of the effect of different distance cutoffs

An example ligand-receptor pair (FGF19-FGFR1) and the Visium data of human breast cancer is used to demonstrate the effect of using different distance cutoffs in COMMOT. **a** The spatial expression of FGF19 and FGFR1. **b** The inferred amount of received signal (spatial distribution of FGF19-FGFR1 complexes with different distance cutoffs in the unit of microns.

### Supplementary Methods

### 1 Collective optimal transport in COMMOT

#### 1.1 Collective optimal transport

Motivated by the biological application of analyzing interactions among multiple species of ligand and receptors, we introduce collective optimal transport (COT) which considers multiple source and target species, unlike traditional optimal transport [1, 2] that only couples two single species. Consider $n_s$ points (cells of spots of cells) on which the discrete distributions are supported. Collective optimal transport couples $n_l$ source species (ligands in the COMMOT application), $\boldsymbol{\alpha}_i \in \mathbb{R}_+^{n_s}, i = 1, \cdots, n_l$ and $n_r$ target species (receptors in the COMMOT application), $\boldsymbol{\beta}_j \in \mathbb{R}_+^{n_s}, j = 1, \cdots, n_r$. Among these species, only part of the species pairs can interact and we use $I$ to denote the indexes of iteracting species such that $(i, j) \in I$ indicates that source species $i$ interacts with target species $j$. For each species pair $(i, j) \in I$, a precomputed cost matrix $\mathbf{C}_{(i,j)} \in \mathbb{R}_+^{n_s \times n_s}$ is given. An example of the cost matrix is the Euclidean distance between the $n_s$ points where a distance entry is replaced by $\infty$ if it exceeds the spatial range of signaling. Collective optimal transport finds a collection of coupling plans $\mathbf{P}_{(i,j)} \in \mathbb{R}_+^{n_s \times n_s}, (i, j) \in I$ for each species pair in $I$ that minimizes the total coupling cost over all pair.

The following optimization problem is formulated:

$$
\begin{aligned}
&\min_{\{\mathbf{P}_{(i,j)}\}\in\mathbf{\Gamma}} \sum_{(i,j)\in I} \langle \mathbf{P}_{(i,j)}, \mathbf{C}_{(i,j)}\rangle_{\mathsf{F}} + \sum_i F(\boldsymbol{\mu}_i) + \sum_j F(\boldsymbol{\nu}_j), \\
&\mathbf{\Gamma} = \left\{ \{\mathbf{P}_{(i,j)}\} : \sum_{j \text{ if } (i,j)\in I} \sum_l \mathbf{P}_{(i,j)}(k,l) \le \boldsymbol{\alpha}_i(k), \sum_{i \text{ if } (i,j)\in I} \sum_k \mathbf{P}_{(i,j)}(k,l) \le \boldsymbol{\beta}_j(l), \mathbf{P}_{(i,j)}(k,l) \ge 0 \right\} \\
&\boldsymbol{\mu}_i(k) = \boldsymbol{\alpha}_i(k) - \sum_{j \text{ if } (i,j)\in I} \sum_l \mathbf{P}_{(i,j)}(k,l), \boldsymbol{\nu}_j(l) = \boldsymbol{\beta}_j(l) - \sum_{i \text{ if } (i,j)\in I} \sum_k \mathbf{P}_{(i,j)}(k,l),
\end{aligned}
\tag{1}
$$

where $F$ penalizes the uncoupled masses $\boldsymbol{\mu}_i$ and $\boldsymbol{\nu}_j$.

In practice, there could be infinity entries in the cost matrices to block certain coupling according to specific interpretations in the applied domain. Also, $\boldsymbol{\alpha}_i$ and $\boldsymbol{\beta}_j$ are not required to be probability distributions. This property enables the application of collective optimal transport to problems where the mass distribution should not be normalized, for example, when the units need to be preserved. Here, the units are the expression levels of various ligand and receptor species and they need not to be normalized individually so that they stay comparable.

#### 1.2 Alternative formulation of COT

To solve the collective optimal transport problem described above, we reformulate the problem. The multiple source and target species are combined to formulate an optimal transport problem of size $(n_s{*}n_l)\times(n_s{*}n_r)$. A new pair of distributions $\boldsymbol{\alpha} \in \mathbb{R}^{n_s * n_l}$ and $\boldsymbol{\beta} \in \mathbb{R}^{n_s * n_r}$ to be coupled is generated such that $\boldsymbol{\alpha}(i * n_s + k) = \boldsymbol{\alpha}_i(k)$ and $\boldsymbol{\beta}(j * n_s + l) = \boldsymbol{\beta}_j(l)$ The corresponding new cost matrix is then defined as

$$
\mathbf{C}(i * n_l + 1 : (i+1) * n_l, j * n_l + 1 : (j+1) * n_l) = \begin{cases} \mathbf{C}_{(i,j)}, \text{ if } (i,j) \in I \\ \infty, \text{ otherwise} \end{cases} \tag{2}
$$

In this setup, the discrete OT problem is formulated as the following,

$$
\begin{aligned}
\min_{\mathbf{P},\boldsymbol{\mu},\boldsymbol{\nu}} \quad & \langle \mathbf{P}, \mathbf{C}\rangle_{\mathrm{F}} + F(\boldsymbol{\mu}) + G(\boldsymbol{\nu}) \\
\text{s.t.} \quad & \mathbf{P} \in \mathbb{R}_+^{m\times n}, \boldsymbol{\mu} \in \mathbb{R}_+^m, \boldsymbol{\nu} \in \mathbb{R}_+^n \\
& \mathbf{P}\mathbf{1}^n = \boldsymbol{\alpha} - \boldsymbol{\mu}, \\
& \mathbf{P}^T\mathbf{1}^m = \boldsymbol{\beta} - \boldsymbol{\nu},
\end{aligned}
\tag{3}
$$

where $m = n_p * n_s$, $n = n_p * n_t$, $\boldsymbol{\mu}$ and $\boldsymbol{\nu}$ are the unpaired masses, and $F$ and $G$ are penalty functions. In this work, we use two specific constructions with $l_1$ or $l_2$ penalty for the untransported mass.

*COT with $l_1$ penalty*

$$
\begin{aligned}
\min_{\mathbf{P},\boldsymbol{\mu},\boldsymbol{\nu}} \quad & \langle \mathbf{P}, \mathbf{C}\rangle_{\mathrm{F}} + \epsilon_p H(\mathbf{P}) + \epsilon_\mu H(\boldsymbol{\mu}) + \epsilon_\nu H(\boldsymbol{\nu}) + \rho(\|\boldsymbol{\mu}\|_1 + \|\boldsymbol{\nu}\|_1) \\
\text{s.t.} \quad & \mathbf{P}\mathbf{1}^n = \boldsymbol{\alpha} - \boldsymbol{\mu}, \\
& \mathbf{P}^T\mathbf{1}^m = \boldsymbol{\beta} - \boldsymbol{\nu}.
\end{aligned}
\tag{4}
$$

*COT with $l_2$ penalty*

$$\begin{aligned} \min_{\mathbf{P},\boldsymbol{\mu},\boldsymbol{\nu}} \quad & \langle \mathbf{P}, \mathbf{C}\rangle_{\mathrm{F}} + \epsilon_p H(\mathbf{P}) + \epsilon_\mu H(\boldsymbol{\mu}) + \epsilon_\nu H(\boldsymbol{\nu}) + \frac{\rho}{2}(\|\boldsymbol{\mu}\|_2^2 + \|\boldsymbol{\nu}\|_2^2) \\ \text{s.t.} \quad & \mathbf{P}\mathbf{1}^n = \boldsymbol{\alpha} - \boldsymbol{\mu}, \\ & \mathbf{P}^T\mathbf{1}^m = \boldsymbol{\beta} - \boldsymbol{\nu}. \end{aligned} \tag{5}$$

In the equations above, $H(\mathbf{x}) = \sum_i \mathbf{x}_i(\ln(\mathbf{x}_i) - 1)$ is an entropy regularization function which is widely used to increase the computational efficiency [3]. The nonnegativity constraints in Eq. (3) are enforced here by the entropy barriers in Eqs. (4) and (5). The entropy terms also enforce a strong convexity which enables efficient numerical solutions.

#### 1.3 Numerical solutions

In this section, we derive two kinds of numerical solutions to the optimization problems in Eqs. (4) and (5).

**COT with $l_1$ penalty** We begin with the Lagrangian associated with Eq. (4),

$$\begin{aligned} \mathcal{E}(\mathbf{P}, \boldsymbol{\mu}, \boldsymbol{\nu}; \mathbf{f}, \mathbf{g}) =& \langle \mathbf{P}, \mathbf{C}\rangle_{\mathrm{F}} + \epsilon_p H(\mathbf{P}) + \epsilon_\mu H(\boldsymbol{\mu}) + \epsilon_\nu H(\boldsymbol{\nu}) + \rho(\|\boldsymbol{\mu}\|_1 + \|\boldsymbol{\nu}\|_1) \\ & - \langle \mathbf{f}, \mathbf{P}\mathbf{1}^n - (\boldsymbol{\alpha} - \boldsymbol{\mu})\rangle - \langle \mathbf{g}, \mathbf{P}^T\mathbf{1}^m - (\boldsymbol{\beta} - \boldsymbol{\nu})\rangle \end{aligned} \tag{6}$$

The first order condition relates $\mathbf{P}$ with the multipliers as

$$\frac{\partial \mathcal{E}}{\partial \mathbf{P}_{ij}} = \mathbf{C}_{ij} + \epsilon_p \log \mathbf{P}_{ij} - \mathbf{f}_i - \mathbf{g}_j = 0 \quad \Rightarrow \quad \mathbf{P} = e^{\frac{\mathbf{f}\oplus\mathbf{g}-\mathbf{C}}{\epsilon_p}}. \tag{7}$$

With $(\epsilon H(\cdot) + \rho\|\cdot\|_1)^*(y) = \epsilon e^{\frac{y-\rho}{\epsilon}}$ and condition in Eq. (7), we have the following Lagrange dual function,

$$\begin{aligned} L(\mathbf{f}, \mathbf{g}) =& \langle \mathbf{f}, \boldsymbol{\alpha}\rangle + \langle \mathbf{g}, \boldsymbol{\beta}\rangle - \epsilon_p \langle e^{\frac{\mathbf{f}\oplus\mathbf{g}-\mathbf{C}}{\epsilon_p}}, \mathbf{1}^{m\times n}\rangle_{\mathrm{F}} \\ & - \epsilon_\mu \langle e^{\frac{\mathbf{f}-\rho}{\epsilon_\mu}}, \mathbf{1}^{m\times n}\rangle_{\mathrm{F}} - \epsilon_\nu \langle e^{\frac{\mathbf{g}-\rho}{\epsilon_\nu}}, \mathbf{1}^{m\times n}\rangle_{\mathrm{F}}, \end{aligned} \tag{8}$$

whose first order derivatives are

$$\begin{aligned} \frac{\partial L}{\partial \mathbf{f}} &= \boldsymbol{\alpha} - e^{\frac{\mathbf{f}}{\epsilon_p}} \odot (e^{-\frac{\mathbf{C}}{\epsilon_p}} e^{\frac{\mathbf{g}}{\epsilon_p}}) - e^{\frac{\mathbf{f}-\rho}{\epsilon_\mu}}, \\ \frac{\partial L}{\partial \mathbf{g}} &= \boldsymbol{\beta} - e^{\frac{\mathbf{g}}{\epsilon_p}} \odot (e^{-\frac{\mathbf{C}^T}{\epsilon_p}} e^{\frac{\mathbf{f}}{\epsilon_p}}) - e^{\frac{\mathbf{g}-\rho}{\epsilon_\nu}}. \end{aligned} \tag{9}$$

These immediately leads to gradient based methods, such as momentum gradient ascent and Nesterov's method [4].

If we set $\epsilon = \epsilon_p = \epsilon_\mu = \epsilon_\nu$, we can easily obtain the explicit first order conditions,

$$\begin{aligned} \mathbf{f} &= \epsilon \log \boldsymbol{\alpha} - \epsilon \log(e^{-\frac{\mathbf{C}}{\epsilon}} e^{\frac{\mathbf{g}}{\epsilon}} + e^{-\frac{\rho}{\epsilon}}), \\ \mathbf{g} &= \epsilon \log \boldsymbol{\beta} - \epsilon \log(e^{-\frac{\mathbf{C}^T}{\epsilon}} e^{\frac{\mathbf{f}}{\epsilon}} + e^{-\frac{\rho}{\epsilon}}). \end{aligned} \tag{10}$$

We now apply the log-sum-exp trick to stabilize computation for small value of $\epsilon$. Note that

$$\log(e^{-\frac{\mathbf{C}}{\epsilon}}e^{\frac{\mathbf{g}}{\epsilon}} + e^{-\frac{\rho}{\epsilon}}) = -\frac{\mathbf{a}}{\epsilon} + \log(e^{\frac{\mathbf{a}}{\epsilon}} \odot e^{-\frac{\mathbf{C}}{\epsilon}}e^{\frac{\mathbf{g}}{\epsilon}} + e^{\frac{\mathbf{a}-\rho}{\epsilon}}) \tag{11}$$

for any $\mathbf{a} \in \mathbb{R}^m$. We now set $\mathbf{a}$ to $\mathbf{f}$ and $\mathbf{g}$ respectively and obtain a stabilized Sinkhorn iteration,

$$\begin{aligned} \mathbf{f}^{(l+1)} &\leftarrow \epsilon \log \boldsymbol{\alpha} + \mathbf{f}^{(l)} - \epsilon \log(e^{\frac{\mathbf{f}^{(l)}}{\epsilon}} \odot e^{-\frac{\mathbf{C}}{\epsilon}} e^{\frac{\mathbf{g}^{(l)}}{\epsilon}} + e^{\frac{\mathbf{f}^{(l)}-\rho}{\epsilon}}), \\ \mathbf{g}^{(l+1)} &\leftarrow \epsilon \log \boldsymbol{\beta} + \mathbf{g}^{(l)} - \epsilon \log(e^{\frac{\mathbf{g}^{(l)}}{\epsilon}} \odot e^{-\frac{\mathbf{C}^T}{\epsilon}} e^{\frac{\mathbf{f}^{(l+1)}}{\epsilon}} + e^{\frac{\mathbf{g}^{(l)}-\rho}{\epsilon}}), \end{aligned} \tag{12}$$

for $l \geq 0$ with arbitary $\mathbf{f}^{(0)}$ and $\mathbf{g}^{(0)}$.

**COT with $l_2$ penalty** The algorithms for the COT with $l_2$ penalty can be derived similarly. To avoid solving the highly nonlinear systems, we first introduce two primal variables $\tilde{\boldsymbol{\mu}}$ and $\tilde{\boldsymbol{\nu}}$ leading to the following minimization problem,

$$\begin{aligned} \min_{\mathbf{P},\boldsymbol{\mu},\boldsymbol{\nu},\tilde{\boldsymbol{\mu}},\tilde{\boldsymbol{\nu}}} \quad & \langle \mathbf{P}, \mathbf{C} \rangle_{\mathrm{F}} + \epsilon_p H(\mathbf{P}) + \epsilon_\mu H(\boldsymbol{\mu}) + \epsilon_\nu H(\boldsymbol{\nu}) + \frac{\rho}{2}(\|\tilde{\boldsymbol{\mu}}\|^2 + \|\tilde{\boldsymbol{\nu}}\|^2) \\ \text{s.t.} \quad & \mathbf{P}\mathbf{1}^n = \boldsymbol{\alpha} - \boldsymbol{\mu}, \\ & \mathbf{P}^T\mathbf{1}^m = \boldsymbol{\beta} - \boldsymbol{\nu}, \\ & \boldsymbol{\mu} = \tilde{\boldsymbol{\mu}}, \quad \boldsymbol{\nu} = \tilde{\boldsymbol{\nu}}. \end{aligned} \tag{13}$$

The associated Lagrangian then reads

$$\begin{aligned} \mathcal{E}(\mathbf{P}, \boldsymbol{\mu}, \boldsymbol{\nu}, \tilde{\boldsymbol{\mu}}, \tilde{\boldsymbol{\nu}}; \mathbf{f}, \mathbf{g}, \mathbf{r}, \mathbf{s}) =& \langle \mathbf{P}, \mathbf{C} \rangle_{\mathrm{F}} + \epsilon_p H(\mathbf{P}) + \epsilon_\mu H(\boldsymbol{\mu}) + \epsilon_\nu H(\boldsymbol{\nu}) + \frac{\rho}{2}(\|\tilde{\boldsymbol{\mu}}\|^2 + \|\tilde{\boldsymbol{\nu}}\|^2) \\ & - \langle \mathbf{f}, \mathbf{P}\mathbf{1}^n - (\boldsymbol{\alpha} - \boldsymbol{\mu}) \rangle - \langle \mathbf{g}, \mathbf{P}^T\mathbf{1}^m - (\boldsymbol{\beta} - \boldsymbol{\nu}) \rangle \\ & - \langle \mathbf{r}, \boldsymbol{\mu} - \tilde{\boldsymbol{\mu}} \rangle - \langle \mathbf{s}, \boldsymbol{\nu} - \tilde{\boldsymbol{\nu}} \rangle. \end{aligned} \tag{14}$$

Taking the first order derivatives leads to the first order conditions,

$$\begin{aligned} \frac{\partial \mathcal{E}}{\partial \mathbf{P}_{ij}} &= \mathbf{C}_{ij} + \epsilon_p \log \mathbf{P}_{ij} - \mathbf{f}_i - \mathbf{g}_j = 0 && \Rightarrow \quad \mathbf{P} = e^{\frac{\mathbf{f} \oplus \mathbf{g} - \mathbf{C}}{\epsilon_p}}, \\ \frac{\partial \mathcal{E}}{\partial \boldsymbol{\mu}_i} &= \epsilon_\mu \log \boldsymbol{\mu}_i - \mathbf{f}_i - \mathbf{r}_i = 0 && \Rightarrow \quad \boldsymbol{\mu} = e^{\frac{\mathbf{f}+\mathbf{r}}{\epsilon_\mu}}, \\ \frac{\partial \mathcal{E}}{\partial \boldsymbol{\nu}_i} &= \epsilon_\nu \log \boldsymbol{\nu}_i - \mathbf{g}_i - \mathbf{s}_i = 0 && \Rightarrow \quad \boldsymbol{\mu} = e^{\frac{\mathbf{g}+\mathbf{s}}{\epsilon_\nu}}, \\ \frac{\partial \mathcal{E}}{\partial \tilde{\boldsymbol{\mu}}_i} &= \rho \tilde{\boldsymbol{\mu}}_i + \mathbf{r}_i && \Rightarrow \quad \tilde{\boldsymbol{\mu}} = -\frac{\mathbf{r}}{\rho}, \\ \frac{\partial \mathcal{E}}{\partial \tilde{\boldsymbol{\nu}}_i} &= \rho \tilde{\boldsymbol{\nu}}_i + \mathbf{s}_i && \Rightarrow \quad \tilde{\boldsymbol{\nu}} = -\frac{\mathbf{s}}{\rho}. \end{aligned} \tag{15}$$

We now obtain the Lagrange dual function by plugging these conditions to the Lagrangian $\mathcal{E}$,

$$
\begin{aligned}
L(\mathbf{f},\mathbf{g},\mathbf{r},\mathbf{s}) = & \min_{\mathbf{P},\boldsymbol{\mu},\boldsymbol{\nu},\tilde{\boldsymbol{\mu}},\tilde{\boldsymbol{\nu}}} \mathcal{E}(\mathbf{P},\boldsymbol{\mu},\boldsymbol{\nu},\tilde{\boldsymbol{\mu}},\tilde{\boldsymbol{\nu}};\mathbf{f},\mathbf{g},\mathbf{r},\mathbf{s}) \\
= & \langle \mathbf{f},\boldsymbol{\alpha}\rangle + \langle \mathbf{g},\boldsymbol{\beta}\rangle - \epsilon_p \langle e^{\frac{\mathbf{f}\oplus\mathbf{g}-\mathbf{C}}{\epsilon_p}}, \mathbf{1}^{m\times n}\rangle_{\mathrm{F}} \\
& + \min_{\boldsymbol{\mu}} \epsilon_\mu H(\boldsymbol{\mu}) - \langle \mathbf{f}+\mathbf{r},\boldsymbol{\mu}\rangle + \min_{\boldsymbol{\nu}} \epsilon_\nu H(\boldsymbol{\nu}) - \langle \mathbf{g}+\mathbf{s},\boldsymbol{\nu}\rangle \\
& + \min_{\tilde{\boldsymbol{\mu}}} \frac{\rho}{2}\|\tilde{\boldsymbol{\mu}}\|^2 - \langle \mathbf{r},\tilde{\boldsymbol{\mu}}\rangle + \min_{\tilde{\boldsymbol{\nu}}} \frac{\rho}{2}\|\tilde{\boldsymbol{\nu}}\|^2 - \langle \mathbf{s},\tilde{\boldsymbol{\nu}}\rangle \\
= & \langle \mathbf{f},\boldsymbol{\alpha}\rangle + \langle \mathbf{g},\boldsymbol{\beta}\rangle - \epsilon_p \langle e^{\frac{\mathbf{f}\oplus\mathbf{g}-\mathbf{C}}{\epsilon_p}}, \mathbf{1}^{m\times n}\rangle_{\mathrm{F}} \\
& - (\epsilon_\mu H)^*(\mathbf{f}+\mathbf{r}) - (\epsilon_\nu H)^*(\mathbf{g}+\mathbf{s}) - (\frac{\rho}{2}\|\cdot\|^2)^*(-\mathbf{r}) - (\frac{\rho}{2}\|\cdot\|^2)^*(-\mathbf{s}),
\end{aligned}
\tag{16}
$$

where $(\epsilon H)^*(y) = \epsilon e^{\frac{y}{\epsilon}}$ and $(\frac{\rho}{2}\|\cdot\|^2)^*(\mathbf{y}) = \frac{\|\mathbf{y}\|^2}{2\rho}$. Similar to the $l_1$ case, gradient based optimization methods can be used given the first order derivatives,

$$
\begin{aligned}
\frac{\partial L}{\partial \mathbf{f}} &= \boldsymbol{\alpha} - e^{\frac{\mathbf{f}}{\epsilon_p}} \odot (e^{-\frac{\mathbf{C}}{\epsilon_p}} e^{\frac{\mathbf{g}}{\epsilon_p}}) - e^{\frac{\mathbf{f}+\mathbf{r}}{\epsilon_\mu}}, \\
\frac{\partial L}{\partial \mathbf{g}} &= \boldsymbol{\beta} - e^{\frac{\mathbf{g}}{\epsilon_p}} \odot (e^{-\frac{\mathbf{C}^T}{\epsilon_p}} e^{\frac{\mathbf{f}}{\epsilon_p}}) - e^{\frac{\mathbf{g}+\mathbf{s}}{\epsilon_\nu}}, \\
\frac{\partial L}{\partial \mathbf{r}} &= -e^{\frac{\mathbf{f}+\mathbf{r}}{\epsilon_\mu}} - \frac{\mathbf{r}}{\rho}, \\
\frac{\partial L}{\partial \mathbf{s}} &= -e^{\frac{\mathbf{g}+\mathbf{s}}{\epsilon_\nu}} - \frac{\mathbf{s}}{\rho}.
\end{aligned}
\tag{17}
$$

Setting $\epsilon = \epsilon_p = \epsilon_\mu = \epsilon_\nu$, using the first order condition, and applying the log-sum-exp trick, we have the following Sinkhorn iteration,

$$
\begin{aligned}
\mathbf{f}^{(l+1)} &\leftarrow \epsilon \log \boldsymbol{\alpha} + \mathbf{f}^{(l)} - \epsilon \log(e^{\frac{\mathbf{f}^{(l)}}{\epsilon}} \odot e^{-\frac{\mathbf{C}}{\epsilon}} e^{\frac{\mathbf{g}^{(l)}}{\epsilon}} + e^{\frac{\mathbf{f}^{(l)}+\mathbf{r}}{\epsilon}}), \\
\mathbf{g}^{(l+1)} &\leftarrow \epsilon \log \boldsymbol{\beta} + \mathbf{g}^{(l)} - \epsilon \log(e^{\frac{\mathbf{g}^{(l)}}{\epsilon}} \odot e^{-\frac{\mathbf{C}^T}{\epsilon}} e^{\frac{\mathbf{f}^{(l+1)}}{\epsilon}} + e^{\frac{\mathbf{g}^{(l)}+\mathbf{s}}{\epsilon}}), \\
\mathbf{r}^{(l+1)} &\leftarrow -\epsilon\omega(\frac{\mathbf{f}^{(l+1)}}{\epsilon} - \log\frac{\epsilon}{\rho}), \\
\mathbf{s}^{(l+1)} &\leftarrow -\epsilon\omega(\frac{\mathbf{g}^{(l+1)}}{\epsilon} - \log\frac{\epsilon}{\rho}),
\end{aligned}
\tag{18}
$$

where $\omega$ is the Wright omega function.

### 2 PDE model for simulated benchmark

#### 2.1 Model

Assume there are $n_l$ ligand and $n_r$ receptor species where each ligand or receptor species might bind to multiple others. Let $[L_i], [R_j], [L_iR_j] : \mathbb{R}^{n_{\mathrm{grid}}\times 2} \to \mathbb{R}$ denote the spatial distributions of ligand $i$, receptor $j$ and the complex formed by their binding, the 1) diffusion and degradation of ligand and 2) binding and dissociation of ligand-receptor complex are modeled in the following equations,

$$\begin{aligned}
\frac{\partial[L_i]}{\partial t} &= D_i \nabla^2 [L_i] - \sum_{j:(i,j)\in I} a_{(i,j)}[L_i][R_j] + \sum_{j:(i,j)\in I} b_{(i,j)}[L_i R_j] - c_i[L_i], \\
\frac{\partial[L_i R_j]}{\partial t} &= a_{(i,j)}[L_i][R_j] - b_{(i,j)}[L_i R_j], \\
\frac{\partial[R_j]}{\partial t} &= - \sum_{i:(i,j)\in I} a_{(i,j)}[L_i][R_j] + \sum_{i:(i,j)\in I} b_{(i,j)}[L_i R_j],
\end{aligned} \tag{19}$$

for $(i,j) \in I$, the index set of ligand and receptor that can bind. For each ligand-receptor pair, ligand $i$ and receptor $j$ that can bind, $a_{(i,j)}$ and $b_{(i,j)}$ are the association rate and dissociation rate between them. For ligand $i$, $c_i$ is its degradation rate and $D_i$ is its diffusivity in space.

#### 2.2 Synthetic data as benchmark

Synthetic benchmark data for validating the collective optimal transport method is generated by simulation of the PDE models described in Eq. (19). Various cases of ligand-receptor binding are considered which have varying numbers of ligand species, receptor species, and ligand-receptor complexes, to account for different complexities and levels of competition among species (SFigures 1-4). For each case, ten independent simulations were carried out. For each simulation, the initial distribution of ligands and receptors were generated by using multiple Gaussian distributions with random bandwidths and random locations in a square domain. All ligand-receptor complexes were initiated to be zero.

For all simulations, a natural boundary condition (no-flux) was assumed and a 2D 50-by-50 grid was used. The simulations were implemented using the py-pde package [5]. The solver was set to ”scipy” which uses the ”scipy.integrate.solve_ivp” utility in SciPy [6] that implements the explicit Runge-Kutta method of order 5(4). The simulation was run until the concentrations of the ligand-receptor complexes reach (numerical) equilibrium. The concentrations of the ligand-receptor complexes depict the spatial distributions of each receptor species occupied by each ligand species. The data analysis methods (optimal transport based method) are then benchmarked based on their performance of reconstructing these spatial distributions.

### References

[1] Cédric Villani. *Optimal transport: old and new*, volume 338. Springer, 2009.

[2] Gabriel Peyré, Marco Cuturi, et al. Computational optimal transport: With applications to data science. *Foundations and Trends® in Machine Learning*, 11(5-6):355–607, 2019.

[3] Marco Cuturi. Sinkhorn distances: Lightspeed computation of optimal transport. *Advances in neural information processing systems*, 26:2292–2300, 2013.

[4] Yurii Nesterov. *Introductory lectures on convex optimization: A basic course*, volume 87. Springer Science & Business Media, 2003.

[5] David Zwicker. py-pde: A python package for solving partial differential equations. *Journal of Open Source Software*, 5(48):2158, 2020.

[6] Pauli Virtanen, Ralf Gommers, Travis E. Oliphant, Matt Haberland, Tyler Reddy, David Cournapeau, Evgeni Burovski, Pearu Peterson, Warren Weckesser, Jonathan Bright, Stéfan J. van der Walt, Matthew Brett, Joshua Wilson, K. Jarrod Millman, Nikolay Mayorov, Andrew R. J. Nelson, Eric Jones, Robert Kern, Eric Larson, C J Carey, İlhan Polat, Yu Feng, Eric W. Moore, Jake VanderPlas, Denis Laxalde, Josef Perktold, Robert Cimrman, Ian Henriksen, E. A. Quintero, Charles R. Harris, Anne M. Archibald, Antônio H. Ribeiro, Fabian Pedregosa, Paul van Mulbregt, and SciPy 1.0 Contributors. SciPy 1.0: Fundamental Algorithms for Scientific Computing in Python. *Nature Methods*, 17:261–272, 2020.
